# Minimal biophysical rules are sufficient for the emergence of computational intelligence at the neuronal scale

**DOI:** 10.64898/2026.03.01.708921

**Authors:** Guanyu Wang, Liang Qi, Kunyang Li, Chenyu Tang, Xuhang Chen, Luigi G. Occhipinti, Arokia Nathan, Ningli Wang, Yu Pan, Peter Smielewski, Ying Wang, Yingyan Mao, Xiaoyu Guo, Shuo Gao

## Abstract

How intelligence emerges from the brain’s complex microscopic physical system is a central question for neuroscience and artificial intelligence. Constrained by the genomic bottleneck that precludes synapse-by-synapse specification, we propose and validate a microscopic structure-function ‘concise-constraint sufficiency’ hypothesis. We develop the Neuro-Informed Generative Connectome (NIGC) framework, and show that connectomes generated under a concise set of biophysical constraints (geometric embedding, node propensity modulation, a global energy budget and maximum-entropy selection) closely match the structural statistics of a measured mouse V1 microcircuit (similarities, 0.997). In parallel, using the generated connectome as the fixed reservoir of an echo state network (ESN), training only a linear readout achieves 90% accuracy on an auditory multi-classification task. Moreover, multidimensional biologically consistent functional phenotypes, such as hierarchical transmission delays and low-dimensional spatiotemporal trajectories, are obtained without fitting functional matrices or time courses. Further, by combining single-constraint ablations, pathological perturbations and cross-modal validation, we clarify how specific structural constraints map onto functional consequences. Together, these results delineate sufficient conditions for computational intelligence emergence at the microscopic scale, and provide an auditable benchmark for first-principles understanding of brain construction.

---

How intelligence emerges remains a central question for neuroscience and artificial intelligence^1^. At macroscopic scales, regional specialisation and network-level coordination have progressively yielded robust regularities of functional organisation, and large-scale connectivity and population activity patterns provide key clues for understanding cognition and behaviour^2^. Yet these regularities do not naturally extend to the microscopic neuronal scale. In a regime of an enormous number of connections and strongly coupled interactions, the mapping between structure and function more readily falls into a high-dimensional and complex space^3^. Consequently, at the neuronal scale, we still lack a principle-driven, mechanistically traceable, and testable route to characterize wiring rules and predict functional consequences. The genomic bottleneck suggests that the genome’s information capacity is orders of magnitude smaller than that of the connectome, which precludes specifying fine-grained microscopic wiring connection by connection by genetic information, and is more likely constrained by compressible generative rules^4^. Turing’s concept of unorganised machines^5^ and his theory of morphogenesis^6^ further indicate that simple local interactions can, in principle, generate coherent structure and rich dynamics. Inspired by these ideas, we propose the testable microscopic structure-function ‘concise-constraint sufficiency’ hypothesis: given geometric-scaling-law embedding and controlled dynamics, a small set of explicit biophysical wiring rules may already be sufficient, empirically, to reproduce key statistical structure of neural connectivity and to support a set of biologically consistent functional phenotypes (Fig. 1a).

**Figure 1.**
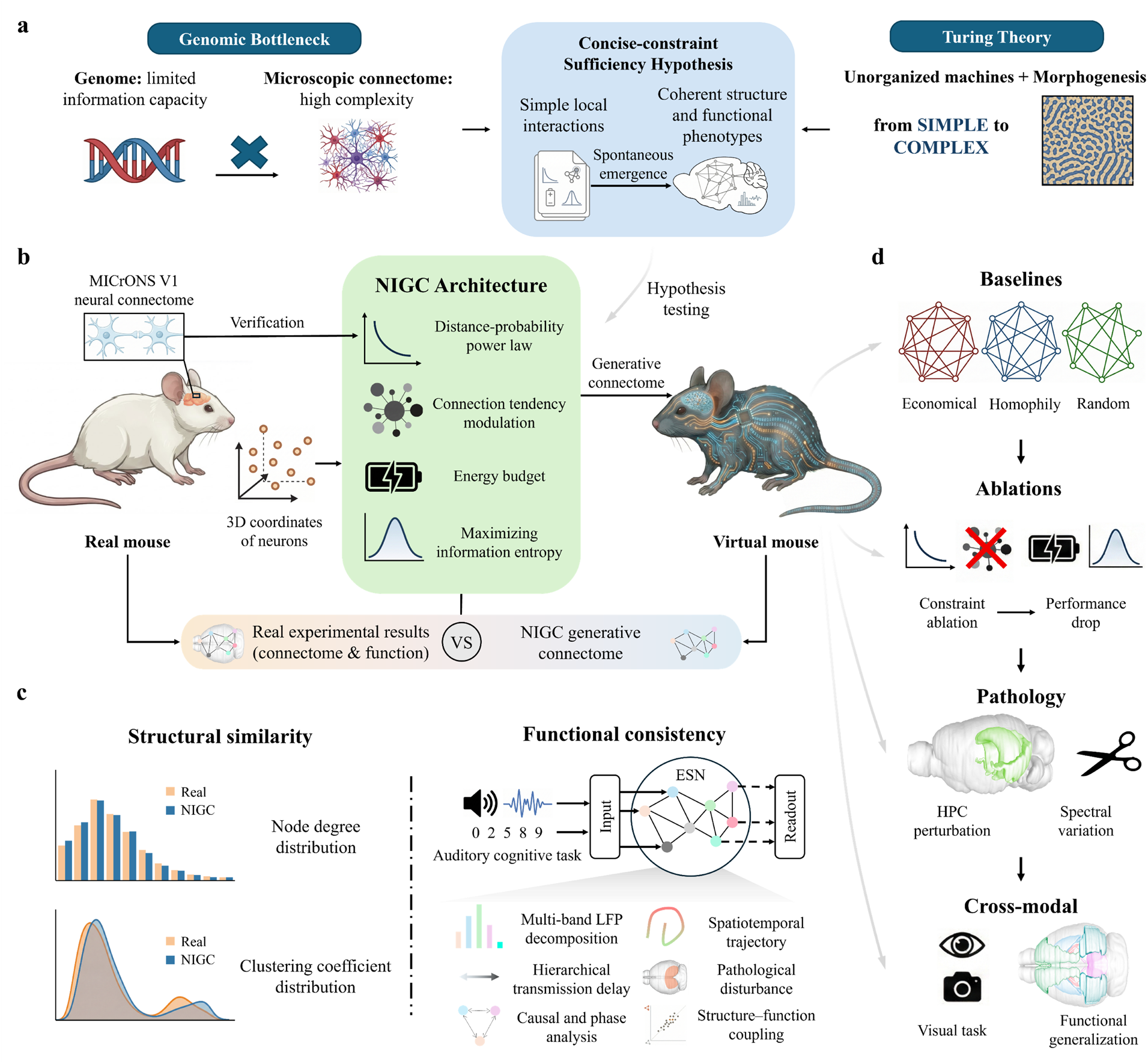
Testing the microscopic structure-function ‘concise-constraint sufficiency’ hypothesis. **a**, Motivation and hypothesis. The genomic bottleneck implies that the genome cannot specify microscopic wiring synapse by synapse, pointing instead to compressible generative rules^4^. Turing’s unorganised machines^5^ and morphogenesis^6^ illustrate how simple local interactions can yield coherent, complex patterns. Inspired by these ideas, we propose the concise-constraint sufficiency hypothesis: whether geometry, together with a small, explicit and ablatable set of biophysical wiring constraints, is sufficient to generate neuronal-scale connectomes that match measured structural statistics and support biologically consistent functional phenotypes under controlled dynamics. **b**, Overview of the sufficiency-test pipeline. Starting from measured neuronal coordinates, MICrONS mouse V1 data^7^ support a power-law distance-connection probability scaling over the observable range, which defines the geometric-scaling-law prior. A concise, explicit and ablatable set of biophysical wiring constraints is then applied to generate a directed neuronal-scale connectome. These constraints include geometric embedding, node-propensity modulation, a global energy budget and maximum-entropy selection, and are implemented here using the Neuro-Informed Generative Connectome (NIGC) framework. **c**, Structural and functional sufficiency tests under controlled dynamics. Structural sufficiency is assessed by benchmarking generated versus measured microcircuit statistics (degree and clustering-coefficient distributions). Functional sufficiency is assessed by instantiating the generated connectome as a fixed reservoir topology in an echo state network (ESN) and training only a linear readout; multidimensional functional phenotypes (spectral fingerprints, hierarchical transmission delays, directed interactions, phase coupling, low-dimensional spatiotemporal trajectories, perturbation effects and structure-function coupling) are quantified under the same unified dynamical regime and compared with biological observations. **d**, Causal and generality validation. Baseline comparisons test whether alternative generative rules (economical wiring, homophily and random) can match the same structural and functional signatures. Single-constraint ablations remove one wiring constraint at a time while keeping the dynamical regime fixed, enabling causal attribution of changes in structure and function to specific constraints. Targeted perturbations selectively rewrite subsets of connections to model structural changes, and test whether these changes produce biologically similar functional consequences under the same dynamical regime. Cross-modal validation applies the same constraint set and analysis pipeline to a visual cognition task to assess generality across sensory modalities.

To rigorously test this hypothesis, we developed the Neuro-Informed Generative Connectome (NIGC) framework (Fig. 1b). The framework aims to reveal the contribution of the structural prior itself to computation without introducing complex cellular dynamics or large-scale learning. Using the synapse-level MICrONS mouse V1 neuronal-scale connectome^7^, we establish a distance-connection probability geometric scaling law and combine it with node propensity modulation^8^, a global energy budget^9^ and maximum-entropy selection^10^.

We then test the hypothesis through sufficiency tests, and through causal and generality validation (Fig. 1c, d). For the former, we show that the generated V1 connectome closely matches the measured microcircuit in the heavy-tailed degree distribution (similarity, 0.997) and clustering. Next, by using an NIGC-generated auditory-pathway connectome as the fixed reservoir of an echo state network (ESN)^11^ and training only a linear readout, we achieve 90% classification accuracy on a spoken-digit classification task^12^ and observe the spontaneous emergence of functional phenotypes including spectral fingerprints, causal interactions and phase coupling, without fitting functional matrices or time courses. For the latter, through controlled ablations and pathological perturbation simulations, we clarify the independent contribution of each constraint to functional consequences, and we further demonstrate cross-modal generalisation with a visual cognition task.

Taken together, we show that a minimal set of biophysical constraints can be sufficient, at the microscopic scale, to generate structural features with computational potential and functional phenotypes that are biologically similar. This provides neuroscience with an auditable structure-function baseline for dissecting disease-related mechanisms, and offers principled support for building more interpretable and transferable neuromorphic intelligence in artificial intelligence.

## Results

### Neuronal-scale distance-connection probability geometric scaling law

To establish the geometric constraint underlying our concise-constraint sufficiency test, we first quantified how neuronal-scale connection probability in the MICrONS mouse V1 microcircuit varies with Euclidean distance. Using the synapse-level MICrONS V1 connectome (Fig. 2a,b), we fitted four distance-decay families (power law^10^, exponential^13^, Gaussian kernel^14^, and logistic^15^) over a common distance range and selected the best model by maximum likelihood and information-criterion comparison. Across the observed range, a power law provided the best fit, with higher log-likelihood and better information-criterion scores than the alternatives (Fig. 2c).

**Figure 2.**
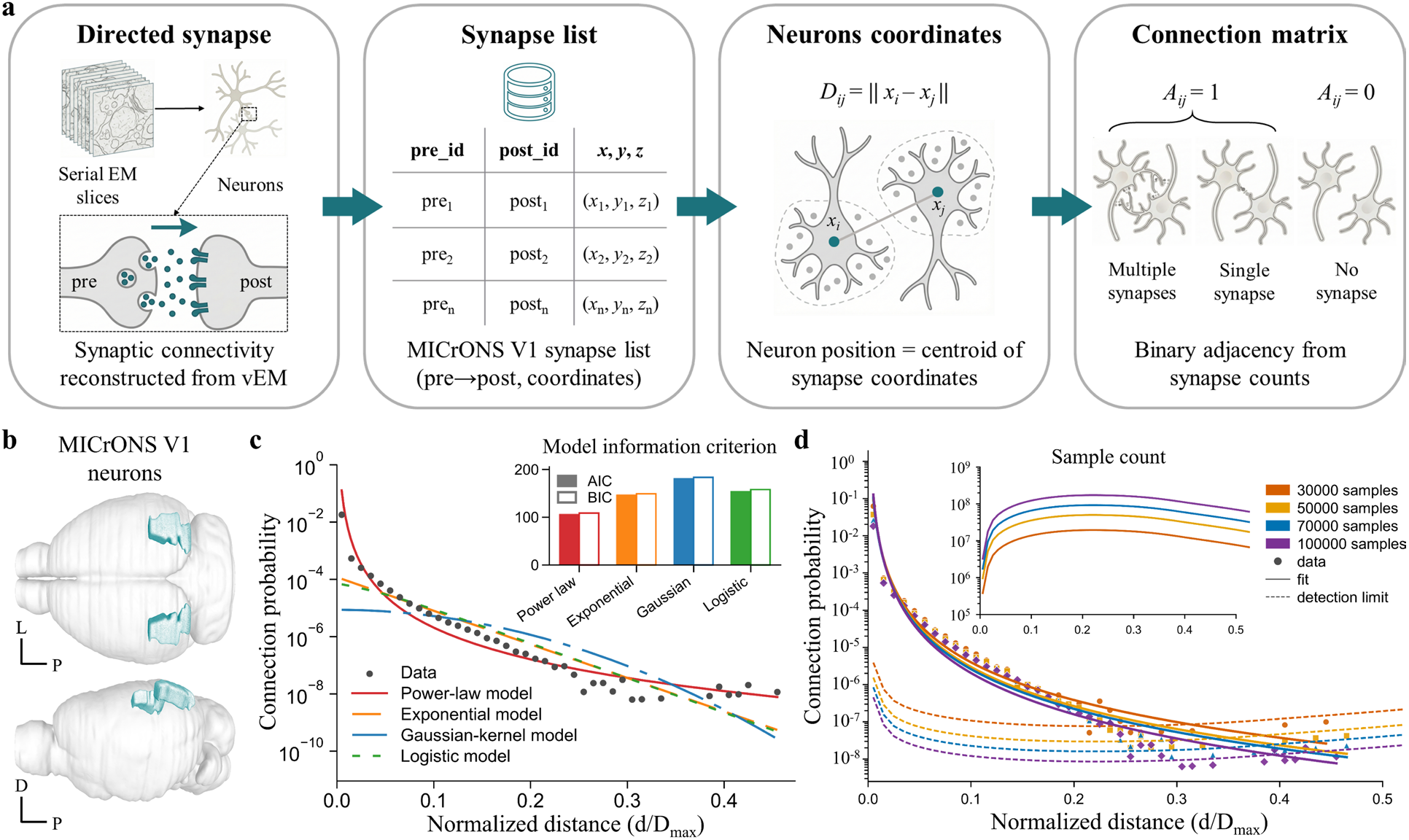
Power-law scaling of the neuronal-scale distance-connection probability in MICrONS mouse V1. **a**, MICrONS V1 data source and preprocessing for neuronal-scale connectivity analysis. Serial EM sections are reconstructed by volume electron microscopy (vEM) to obtain directed synapses (pre→post)^7^. From the MICrONS V1 synapse table, we extracted pre_id and post_id (presynaptic and postsynaptic neuron IDs) and 3D synapse coordinates (*x, y, z*), defined each neuron’s position as the centroid of its associated synapse coordinates, and computed centroid-to-centroid Euclidean distance *D*_ij_. A directed binary adjacency matrix *A*_ij_ was derived from synapse counts, with *A*_ij_=1 if at least one synapse from neuron *i* to neuron *j* is present and *A*_ij_=0 otherwise. **b**, Location of the MICrONS sampling volume in mouse primary visual cortex (V1; blue)^60^. Axes: L, lateral; P, posterior; D, dorsal. **c**, Model comparison for the empirical decay of connection probability as a function of soma-to-soma Euclidean distance. Black dots denote binned empirical probabilities; curves show maximum-likelihood fits of power-law, exponential, Gaussian kernel and logistic models over the same distance range. Top right, AIC/BIC comparisons across the four models. **d**, Subsampling test of long-distance sparsity and apparent downward deviation. The distance-probability relationship is re-estimated under different subsampling scales (30k, 50k, 70k and 100k), together with the corresponding power-law fits and detection limits (dashed lines). Distances in **c** and **d** are normalised by the diagonal length (*D*_max_) of the sampling-volume bounding box (that is, *d*/*D*_max_), where *d* is the Euclidean distance between neuronal centroids (after converting voxel coordinates to physical units).

At the largest distances, a small fraction of estimates showed increased variance and fell slightly below the fitted curve. This deviation is expected from finite-volume boundaries and sampling sparsity: far-distance bins contain few neuron pairs, so estimates approach the noise floor. Across subsampling scales (30k-100k), the deviation remained confined to the largest distances, whereas power-law scaling over short-to-intermediate distances was robust (Fig. 2d). Together, these analyses support power-law scaling for the mouse V1 distance-connection probability relationship over the observed range. We use this power-law distance prior as the geometric constraint in all subsequent analyses.

### Structural similarity of the generative connectome

Building on the distance-probability scaling law, we constructed NIGC by adding three explicit, ablatable wiring constraints: node propensity modulation^8^, a global energy budget^9^ and maximum-entropy selection^10^ (Fig. 3a). Given three-dimensional neuronal coordinates from MICrONS mouse V1, NIGC generates a directed neuronal-scale connectome. We then performed a structural sufficiency test by benchmarking its structural statistics against the measured microcircuit.

**Figure 3.**
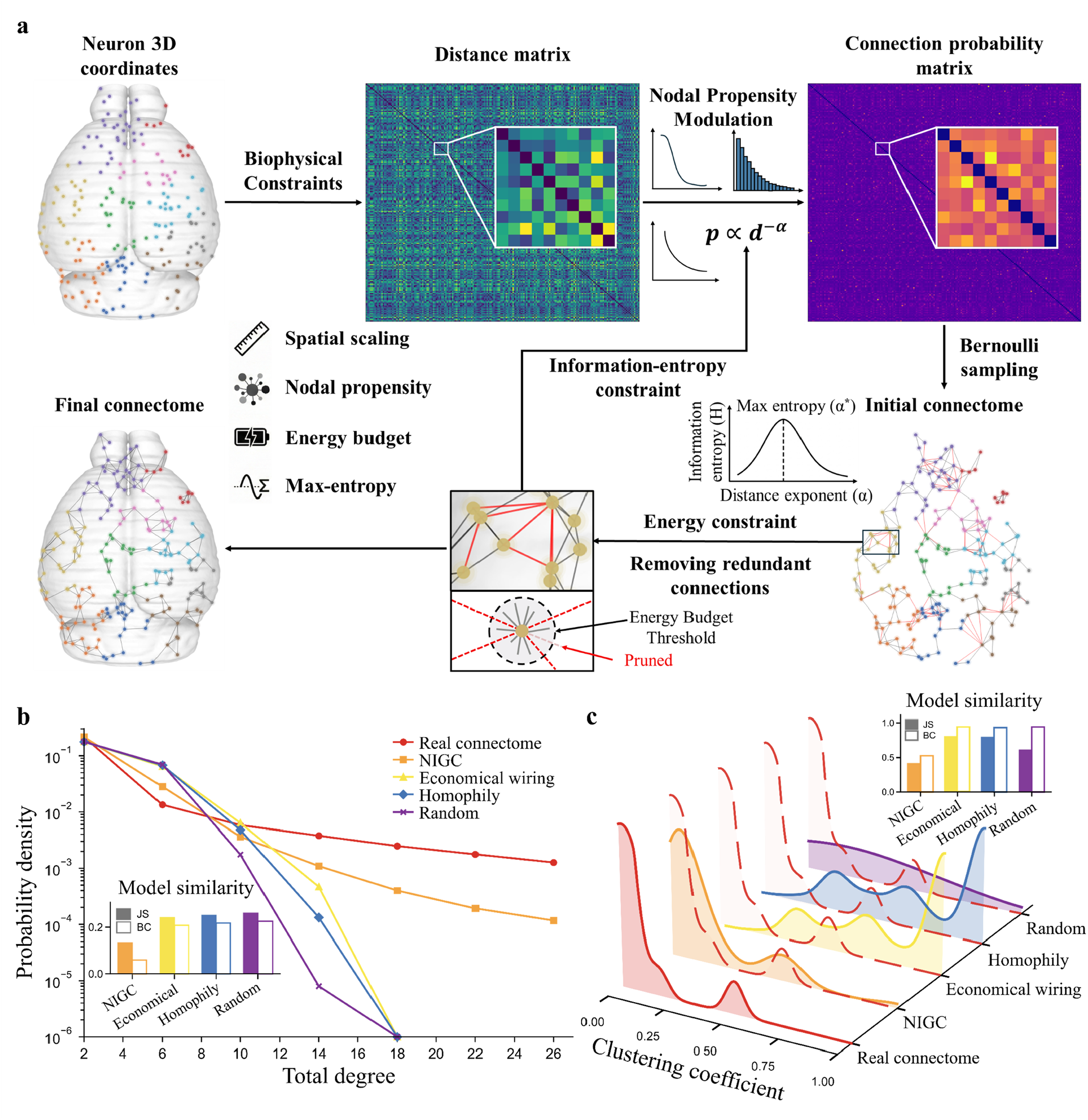
NIGC connectome generation and structural benchmarking against a measured microcircuit. **a**, Schematic of NIGC. Three-dimensional neuronal coordinates yield a pairwise distance matrix, from which baseline connection probabilities are defined by a power-law distance prior. Node propensity modulation is applied to induce degree heterogeneity, yielding a connection-probability matrix. Bernoulli sampling produces an initial connectome, after which redundant connections are pruned under a global energy budget constraint. The generation-and- pruning procedure is then repeated under a maximum-entropy constraint to determine the optimal power-law parameter, yielding the final neuronal-scale connectome. **b**, Degree-distribution comparison. Probability densities of node degree for the measured connectome, the NIGC-generated connectome and three baseline generative models (economical wiring, homophily and fully random). The inset bar chart reports distributional discrepancy metrics (Jensen-Shannon divergence and Bray-Curtis dissimilarity) relative to the measured connectome. **c**, Clustering-coefficient distribution comparison for the same set of networks. The inset bar chart reports Jensen-Shannon divergence and Bray-Curtis dissimilarity relative to the measured connectome. All networks were constructed under matched control conditions, including identical node sets and mean degree.

NIGC reproduces the heavy-tailed degree distribution of the empirical connectome (Fig. 3b; cosine and Pearson similarities, 0.997; JS, 0.133; Bray-Curtis, 0.059). The far tail showed slight compression, with a modest underestimation of the most highly connected nodes. This likely reflects the global energy-budget constraint on in- and out-degrees, compounded by finite-volume sampling limits at long distances (Supplementary Fig. 1i). The clustering-coefficient distribution is also comparable (Fig. 3c; KS, 0.480; JS, 0.413; Bray-Curtis, 0.526), suggesting that NIGC captures both degree heterogeneity and non-saturated local triadic closure.

To test constraint specificity, we benchmarked NIGC against three baselines with matched node set and mean degree: economical wiring, homophily and a mean-degree-matched random network. Economical wiring^9^ and homophily^16^ over-regularised the structure, characterized by inflated local closure while suppressing hub-like degree heterogeneity, whereas the random network produced an approximately Poisson degree profile with clustering concentrated near zero (Fig. 3b,c; Supplementary Tables 2 and 3). None of the baselines simultaneously recovered the heavy-tailed degree range and the moderate, heterogeneous clustering of the measured microcircuit.

We then ablated each core constraint in NIGC. Removing node propensity modulation, the global energy budget or maximum-entropy selection reduced agreement in degree and/or clustering across multiple similarity metrics (Supplementary Fig. 1; Supplementary Tables 4 and 5). Together, these results indicate complementary contributions of the constraints, jointly supporting the degree heterogeneity and the non-saturated level of local closure observed in the measured microcircuit.

### Task-level functional validation in spoken-digit classification

Structural similarity alone does not guarantee functional sufficiency. We therefore tested whether the structurally matched connectomes support executable computation under a fixed, unified dynamical regime when used as recurrent substrates. Specifically, we instantiated each generated connectome as the fixed reservoir topology of an echo state network (ESN)^11^ and trained only a linear readout for spoken-digit classification^12^.

We generated a neuronal-scale connectome for 13 regions (CN, Pons, IC, ACx, HPC, MGB, TRN, LP, SP, PL, IL, OFC and FP) spanning the auditory pathway^17-19^ and prefrontal network and used it as the fixed reservoir in an ESN (Fig. 4a,b; Supplementary Fig. 13). On the Spoken Arabic Digit classification task, the ESN reached 90% test accuracy without any task-driven optimisation of reservoir weights; only the linear readout was trained, indicating that the structural prior can support temporal classification. For context, reported ESN performance on this benchmark typically falls in the ≈ 0.88-0.98 accuracy range across reservoir configurations^20^. Our 90% test accuracy sits within this regime using a NIGC-generated, fixed recurrent reservoir with only a trained readout.

**Figure 4.**
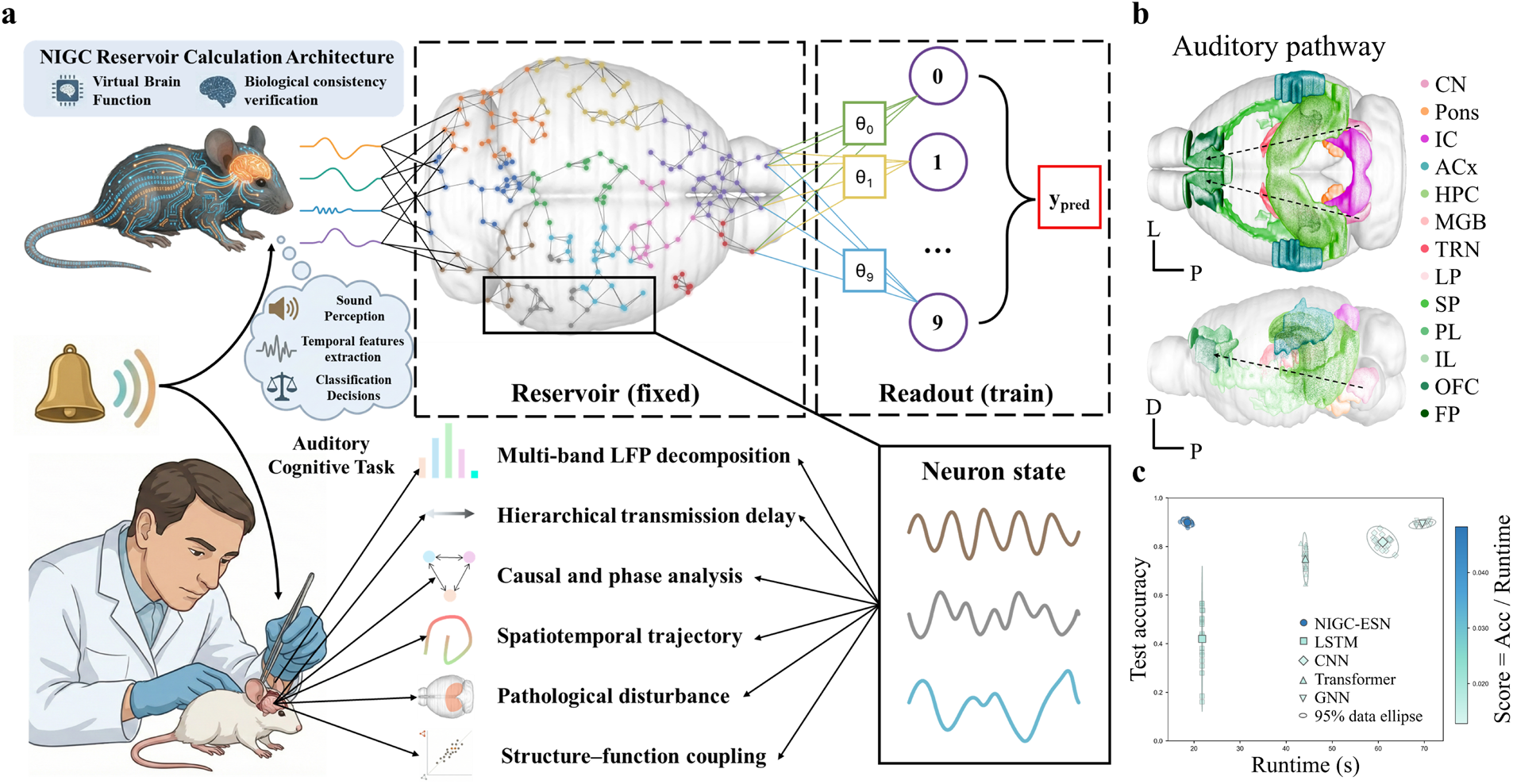
Reservoir computing with NIGC connectomes for task-level functional validation. **a**, NIGC-ESN architecture. The neuronal-scale connectome generated by NIGC is used as the fixed recurrent weight matrix of the reservoir. Spoken Arabic Digit stimuli drive reservoir dynamics, and the readout layer is trained and tested on 10 digit classes while training only the readout parameters. Using the same reservoir responses, functional phenotypes are quantified at multiple levels: basic (multiband local field potential (LFP) spectral fingerprints, hierarchical transmission delays, cross-regional directed interactions and phase coupling), intermediate (low-dimensional trajectories) and higher-level (perturbation effects and structure-function coupling), and compared with relevant biological findings. **b**, The 13 brain regions along the auditory pathway used for functional analysis (colour-coded)^60^. Axes: L, lateral; P, posterior; D, dorsal. Dashed arrows indicate the auditory-pathway direction, from the sensory endpoint CN toward higher-order PFC regions. Region abbreviations: CN, cochlear nuclei; IC, inferior colliculus; ACx, auditory cortex; HPC, hippocampus; MGB, medial geniculate body; TRN, thalamic reticular nucleus; LP, lateral posterior nucleus; SP, cortical subplate; PL, prelimbic cortex; IL, infralimbic cortex; OFC, orbitofrontal cortex; FP, frontal pole. **c**, Performance-efficiency comparison on the Spoken Arabic Digit classification task. The x-axis shows runtime (s) and the y-axis shows classification accuracy; colour indicates a composite score (accuracy divided by runtime). Small points represent repeated independent runs and large points denote the mean. Error ellipses summarize joint uncertainty in accuracy and runtime. NIGC-ESN is compared with end-to-end baselines (LSTM, CNN, Transformer and GNN).

We compared NIGC-ESN with end-to-end sequential and graph models under matched training settings and parameter budgets. NIGC-ESN matched baseline accuracy with lower training cost, as the reservoir is fixed and only a linear readout is trained (Fig. 4c; Supplementary Tables 6 and 7). The resulting trade-off therefore mainly reflects the structural prior, rather than gains from task-specific learning of reservoir weights or architecture search.

### Dynamical fingerprints of generated connectome

Having established task performance, we next examined whether the same NIGC-generated connectomes express biologically similar mesoscopic dynamical fingerprints under the same unified dynamical regime. Specifically, we quantified four aspects of task-driven regional organisation: spectra, hierarchical delays, directed interactions and phase coupling.

### Spectral fingerprints of multiregion LFP

We quantified band-specific power in regional local field potentials (LFPs)^21^ and identified distinct spectral fingerprints across six representative regions (Fig. 5b,c). Overall, beta power dominated across regions, with delta and gamma contributing next most strongly under the spoken-digit task. Experimental studies report prominent beta-band activity in auditory-prefrontal circuits during auditory processing and feedback modulation^22^. Band ordering and relative strength, however, vary across paradigms and region sets.

**Figure 5.**
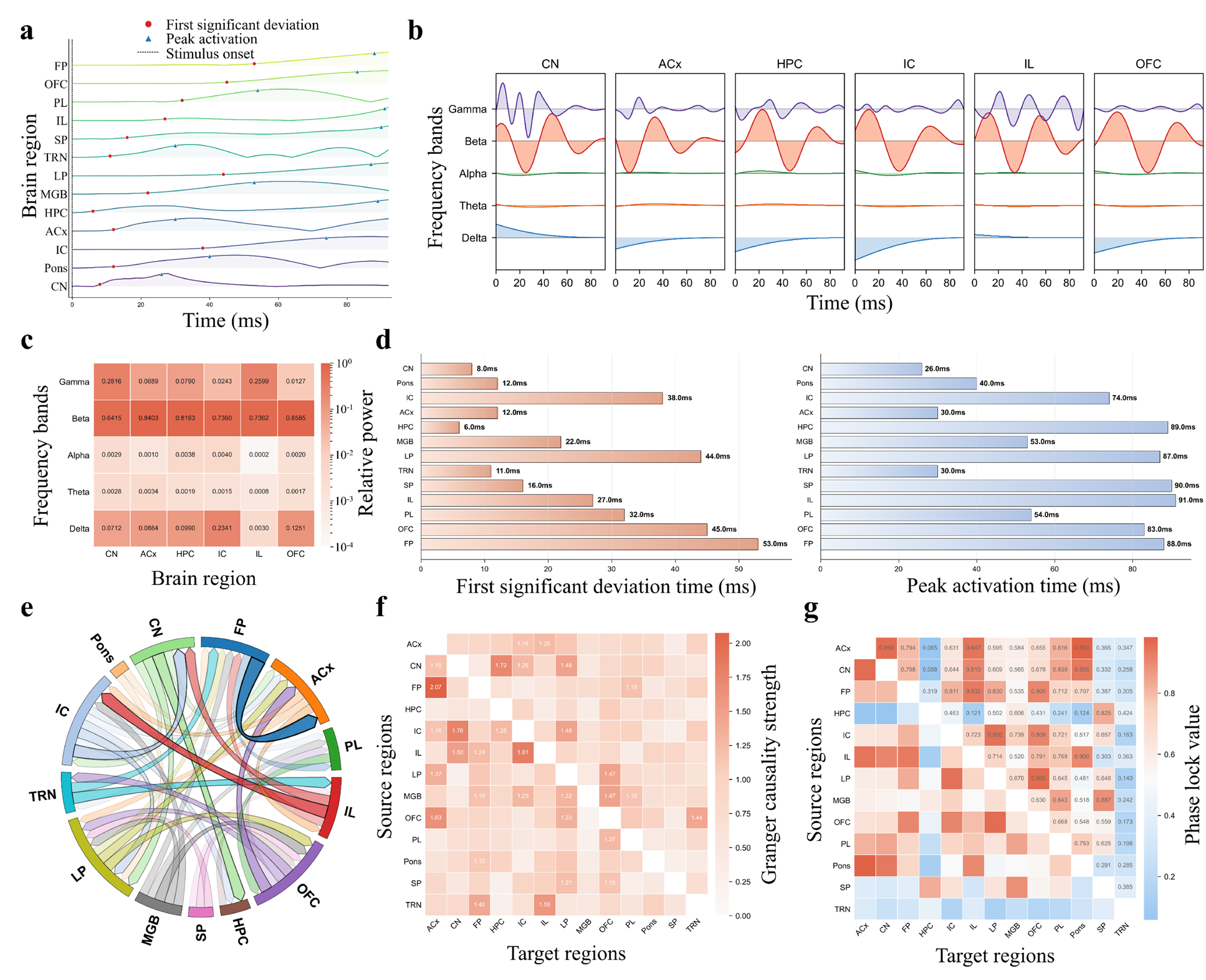
Basic functional phenotypes in the auditory task: spectral fingerprints, hierarchical delays and cross-regional interactions. **a**, Region-level local field potential (LFP) time series for the 13 auditory-pathway-related regions (one row per region). Red circles and blue triangles mark the first time points at which each region reaches 10% and 90% of its peak absolute amplitude, respectively. **b**, Multiband decomposition in representative regions (CN, ACx, HPC, IC, IL and OFC), showing time-frequency components in the δ, θ, α, β and γ bands. **c**, Heat map of relative band power across five bands (spectral fingerprints) for six representative regions (colour indicates relative power). **d**, Summary of onset latencies (10%) and rise-to-peak latencies (90%) across the 13 regions. **e**, Chord-diagram visualisation of cross-regional Granger-causal connections. Edge direction is from source to target and line width indicates causal strength. Networks retain the top 25% strongest edges. **f**, Multivariate Granger-causal strength matrix (source × target) across the 13 regions. **g**, Phase-locking value (PLV) matrix between all region pairs.

IC was beta-dominant (≈0.736) with appreciable delta power (≈0.234). These bands have been linked to predictive/feedback signalling (beta)^23^ and envelope tracking (delta)^24^, consistent with reports of IC involvement in envelope encoding. OFC combined a beta peak with relatively elevated delta power, consistent with reported roles of delta rhythms in envelope-related processing^25^. IL showed relatively elevated gamma power, a pattern often reported in prefrontal cortex during demanding cognitive conditions^26^. CN exhibited elevated gamma power (≈0.282), consistent with reports that early auditory nuclei emphasize fast temporal processing^27^. Because we did not fit spectral features, these patterns arose from the structural prior under the unified dynamics. We therefore interpret the correspondences with prior electrophysiology as qualitative, trend-level similarity rather than strict quantitative matching.

### Hierarchical transmission delays along the auditory pathway

We quantified post-stimulus timing using robust latency metrics^28^ and derived regional response sequences (Fig. 5a,d). Under task drive, responses propagated along an ascending hierarchy: CN activated first (onset ≈8 ms), then propagated through Pons, IC, MGB and LP^19^. ACx showed a primary response within an ≈ 12-30 ms window, whereas PFC regions activated later (≈27-53 ms) and peaked around 83-91 ms^29^. The timing is consistent with the canonical auditory hierarchy (CN→IC→MGB→ACx→PFC) reported in recordings across behavioural and task conditions^19,30^.

ACx and TRN activated with similar short latencies (≈11-12 ms). This pattern is consistent with reports of short-latency thalamic influence on deep auditory cortex (L6) within <10 ms^31^. Cross-regional delays were on the order of a few milliseconds (for example, ≈1 ms for ACx-TRN and ≈5-6 ms for ACx-HPC and IL-PL). Reported myelination-dependent conduction speeds can yield thalamus-to-cortex arrival times in a narrow 1-4 ms window, and our simulated delays are in the same range^32^. TRN peaked early, whereas prefrontal peaks were delayed, matching the expected ordering of thalamic gating and later prefrontal modulation^29^. Overall, the model reproduces millisecond-scale propagation timing consistent with the anatomical auditory hierarchy without fitting functional time courses. Intrinsic time scales under resting-state-like conditions showed a similar hierarchy^33^ (Supplementary Fig. 2b,c), suggesting a link between propagation delays and intrinsic temporal integration.

### Directed interactions along thalamus-cortex-prefrontal pathways

Granger causality indicated directed interactions among the 13 regions, aligned with thalamus-cortex-prefrontal pathways (Fig. 5e). The strongest quartile of edges concentrated along CN-IC-MGB-ACx-PFC and highlighted an ACx-HPC interaction. This aligns with reports of bidirectional ACx-HPC exchange^34^, with direction depending on task phase. In this setting, the ACx-HPC link is among the most prominent directed interactions. ACx→OFC and ACx→FP edges were also among the strongest, consistent with reported sensory-cortical input to OFC in flexible categorization and decision making^35^.

The Granger causality matrix further supports a structured directed network across thalamus, ACx and PFC (Fig. 5f). ACx and thalamic nuclei (MGB, TRN and LP) showed high bidirectional causal strengths (≈0.75-1.37). PFC exhibited bidirectional coupling with thalamic nuclei and ACx (≈0.80-2.07), supporting reciprocal interactions along thalamo-cortico-prefrontal pathways^36^. The pattern is compatible with predictive-coding accounts, in which gamma and beta rhythms have been linked to feedforward and feedback signalling, respectively. Consistent with the spectral fingerprints, beta dominates downstream, whereas gamma is relatively elevated at early relays^37^. Causal strengths were dominated by intermediate edges, with a small high-strength tail (Supplementary Fig. 2d), suggesting a few core pathways superimposed on broader background coupling^38^.

### Rhythmic phase coupling within the auditory pathway

PLV and phase consistency indicate a tightly coupled PFC subnetwork (IL, PL, OFC and FP) (PLV ≈0.668-0.932; phase consistency ≈0.635-0.891) (Fig. 5g; Supplementary Fig. 7g). ACx also showed high phase locking with IL and PL (PLV ≈0.947 and ≈0.816), consistent with beta-dominant long-range coordination^39^. Other long-distance pairs showed weaker PLV synchrony, indicating that rhythmic coordination is concentrated within task-relevant subnetworks. Mutual information highlighted several pathways with nonlinear dependence beyond phase synchrony (Supplementary Fig. 7j), including ACx-LP, IC-PFC and intra-PFC pairs. Consistent with this, supplementary measures (including amplitude-and correlation-based indices, phase stability and power allocation) showed similar pathway-level patterns (Supplementary Fig. 7). Across metrics, several pathways showed co-variation: higher directed Granger coupling tended to coincide with stronger phase locking and higher mutual information. The metrics suggest that directed coupling, phase locking and nonlinear dependence often co-occur along the same pathways^40^.

### From trajectories to system-level coupling

We next asked whether neuronal population activity forms reproducible low-dimensional trajectories. We projected each region’s population states into a low-dimensional space and analysed the resulting trajectories over time. ACx, PL and OFC showed distinct task-structured trajectories (Fig. 6a). Late after stimulus onset, ACx followed an approximately ring-like trajectory, similar in geometry to reported OFF-response trajectories in auditory cortex^41^. Because trajectory features are not fitted, the low-dimensional patterns arise as a consequence of the structural prior and dynamics. In PFC regions, trajectories ramped along a dominant direction and then turned near the decision stage. The ramp-to-turn geometry resembles reported mPFC trajectories in auditory-cue paradigms^42^ and reported late-stage reversals in OFC during tone/noise tasks^43^. Because the tasks differ, we interpret the similarity as qualitative rather than a strict quantitative match. TRN and MGB showed simpler trajectories with prominent turns, consistent with their proposed relay/gating functions^44,45^ (Supplementary Fig. 10a).

**Figure 6.**
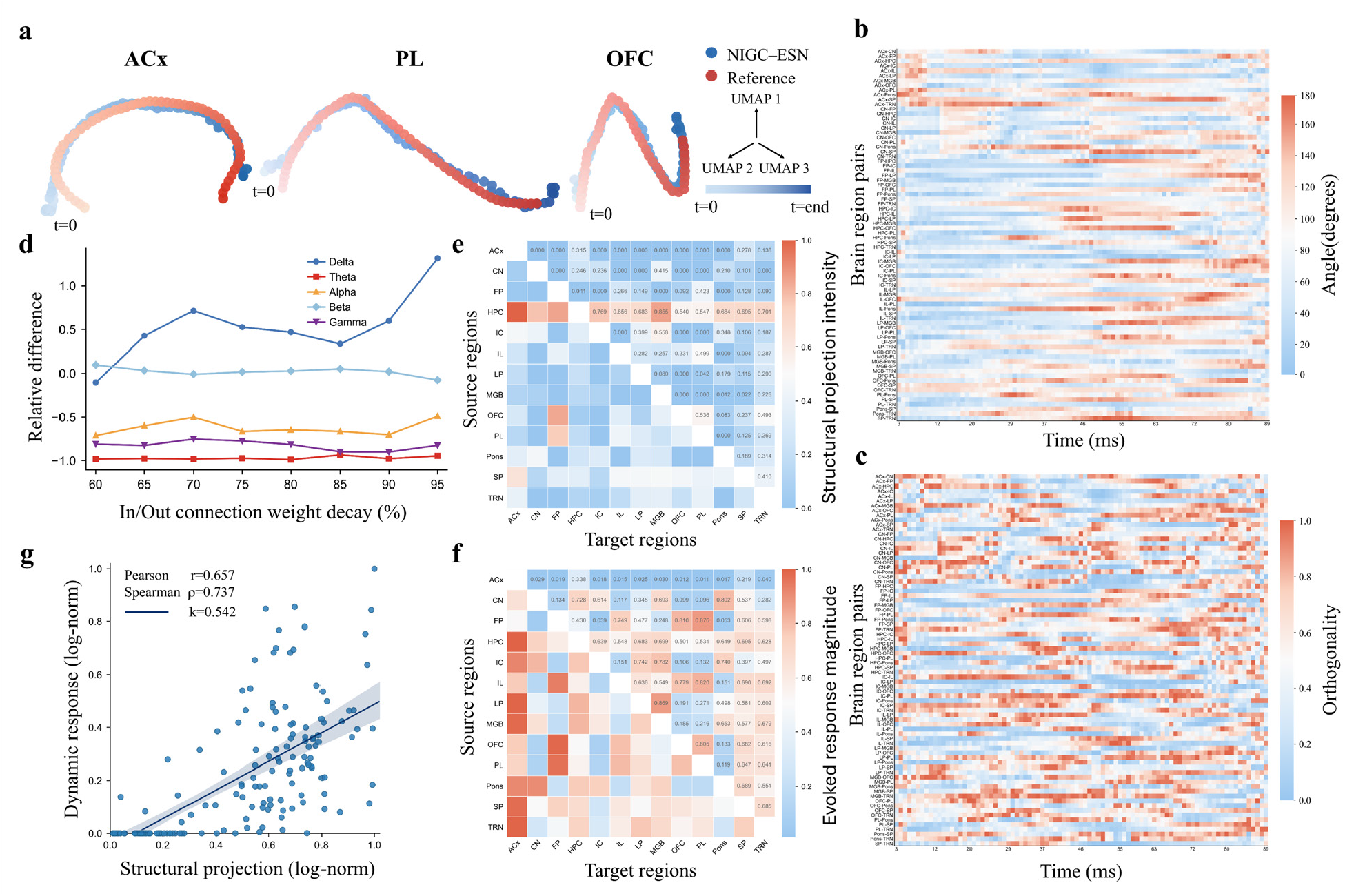
Intermediate and higher-level functional phenotypes: low-dimensional trajectories, perturbation effects and structure-function coupling. **a**, Low-dimensional trajectories in representative regions (ACx, PL and OFC). Task-evoked regional state trajectories are projected into a common embedding space. Blue indicates trajectories from NIGC-ESN and red indicates reference trajectories reported previously (colour intensity indicates time)^41-43^. **b**, Heat map of time-varying angles between trajectories for all region pairs among the 13 regions. The x-axis denotes time steps and the y-axis denotes region pairs; colour indicates trajectory angle. **c**, Heat map of time-varying orthogonality between trajectories for all region pairs. Axes are as in **b**; colour indicates orthogonality strength. **d**, Changes in the HPC rhythmic spectrum under perturbation simulations. After graded attenuation of incoming and outgoing HPC connection weights, changes (relative to the unperturbed condition) in band power are quantified across δ, θ, α, β and γ bands. **e**, Structural projection-field matrix. Each region is treated as a projection source; normalized multi-hop projection strengths to other regions are summarized into a source × target matrix. **f**, Evoked causal response matrix. Brief pulse stimulation is applied to each region, and the response amplitude of each region’s LFP within a time window is recorded to form a stimulus-source × response-target matrix. **g**, Quantification of structure-function coupling. Corresponding source-target entries from **e** and **f** are paired to yield a scatter distribution; Pearson and Spearman correlations quantify consistency between structural projection patterns and evoked causal responses.

We quantified cross-regional coordination using tangent-vector angles and trajectory orthogonality in short windows (Fig. 6b,c). Early trajectories were directionally aligned, with increasing divergence later. Sensory input regions diverged earlier, whereas higher-order regions diverged more slowly, consistent with longer intrinsic time scales higher in the hierarchy^33^. Results were robust to the time-window size over a reasonable range (Supplementary Fig. 10b).

### Pathological perturbations: functional reconfiguration after selective weakening

To probe mechanistic consistency under pathology-like perturbations, we perturbed specific brain regions while keeping structure and dynamics otherwise unchanged. As intra-HPC connectivity was progressively weakened (60% to 95% attenuation of incoming and outgoing HPC weights), the HPC LFP spectrum exhibited a band-selective reconfiguration dominated by a sharp loss of θ (−93.3 to −98.9%) and γ power (−75.2 to −90.0%) alongside a concomitant increase in δ (+42.6 to +131.4%). By contrast, α showed a secondary decrease (−48.9 to −71.2%), whereas β remained near baseline (−7.5% to +9.7%) (Fig. 6d). These trends are consistent with reports of reduced theta/gamma and increased delta activity in hippocampal pathology^46^. In vivo unilateral selective hippocampal ischemia similarly reduces θ and γ components, with θ burst amplitude dropping to 78.0 ± 6.2% of pre-insult levels and low-γ baseline/burst to 61.9 ± 3.3%/57.6 ± 2.8%^47^. These results suggest that NIGC can generate perturbation-induced changes that are mechanistically grounded and may help formulate testable hypotheses.

### Projection-causality: structure-function coupling analysis

We quantified system-level structure-function coupling using a projection-causality comparison inspired by intervention-response paradigms (Fig. 6e-g). Projection strength covaried monotonically with pulse-evoked causal responses across pairs (Pearson’s r = 0.657; Spearman’s ρ = 0.737; Fig. 6g). These correlations suggest that structural projections constrain causal propagation. The magnitude is comparable to correlations reported in intervention-response mapping at broader scales. For example, in mice, axonal tracer-derived anatomical connectivity has been reported to correlate with rs-fMRI coupling profiles with a mean Pearson correlation of r ≈ 0.46^48^. Without fitting functional matrices, pulse-evoked responses still yielded coupling of similar magnitude to mesoscopic reports, suggesting that the NIGC prior captures propagative constraints of plausible magnitude at the system level.

Coupling varied substantially across regions. Source-stratified correlations were high for most regions (median ≈0.816; IC ρ = 0.954), consistent with multi-hop structure shaping outward effects (Supplementary Fig. 2e). Target-stratified coupling was highest for CN/IC and lowest for HPC (Supplementary Fig. 2f). Relay nuclei show tighter structure-causality coupling than hippocampus, suggesting region-dependent contributions of local dynamics^49^. Projection-causality coupling provides an independent quantitative axis for assessing structure-function coupling.

## Stress tests of functional consistency

### Functional comparisons with baselines and constraint ablations

We repeated all functional analyses with matched-parameter baseline connectomes (economical wiring, homophily and fully random) to test dependence on the NIGC prior (Supplementary Figs. 3 and 4). Baselines did not reproduce the joint set of functional fingerprints: economical wiring/homophily weakened coupling and backbone structure, whereas random connectivity increased global synchrony and reduced regional specificity.

We then repeated the functional analyses after ablating each NIGC constraint in turn (power-law distance, node propensity, global energy budget and maximum-entropy selection) (Supplementary Figs. 5 and 6). Each ablation measurably altered the functional signatures in directions consistent with its structural effects. Ablating the global energy budget constraint weakened temporal hierarchy and compressed the propagation ordering. Ablating the power-law distance or maximum-entropy constraint reduced the coherence of the causal backbone and phase coupling along key pathways. Node-propensity ablation reduced regional specificity and weakened hierarchical coordination. Together, the results suggest that the functional fingerprints depend on the NIGC structural prior under the same fixed dynamics, rather than on reservoir dynamics alone.

### Cross-modal generalization: validation of functional consistency in a visual cognition task

We tested generality by switching to a three-class video task along the visual pathway without changing generation rules or dynamics, and reapplying the same analysis pipeline (Supplementary Figs. 11, 12 and 14). The visual task produced hierarchical organisation qualitatively consistent with experimental reports. Spectrally, beta/gamma modulation appeared, and gamma increased along the hierarchy^50^. Temporal dynamics showed staged propagation from early inputs to cortex and then to prefrontal regions. Interactions suggested an ascending backbone centred on visual cortex, with additional top-down influences^51^. Together, these results suggest cross-modal generality of the functional signatures, reducing the likelihood that they arise from fitting to a single task.

## Discussion

Understanding how intelligence emerges at the microscopic neuronal scale is fundamental to explaining how structural alterations reshape function and for formulating testable mechanistic predictions^7^. It offers a powerful causal analysis tool for both neuroscience, particularly in dissecting disease mechanisms, and artificial intelligence, in the quest for interpretable neuromorphic architectures^52^. To date, microscopic structure-function studies have largely followed three paradigms: (1) Task-optimized neural networks^53^, which often treat structural priors as a “black box” obscuring the specific biophysical contributions to intelligence; (2) Connectome generative models^54^, which often remain focused on reproducing structural statistics and more rarely assess multidimensional functional phenotypes under controlled dynamics; and (3) Large-scale biophysical simulations^55^, whose parameter complexity makes it nearly impossible to isolate the independent contributions of the geometric backbone from cellular-level details (see Supplementary Table 8 for details).

These gaps necessitate a move toward first-principles derivation. We therefore propose and validate the testable ‘concise-constraint sufficiency’ hypothesis: that a minimal set of explicit, ablatable biophysical wiring rules can empirically reproduce key neuronal-scale structural statistics and spontaneously yield biologically consistent functional phenotypes without the need to fit functional matrices or time courses. This method renders the generative mechanisms linking structure and function both traceable and separable.

Importantly, our results establish a fundamental null model serving as an auditable structural benchmark for understanding how computational intelligence emerges from minimal biophysical priors. In the current implementation, we deliberately adopted a clean, low-dimensional baseline by omitting secondary biological complexities such as cell-type diversity, laminar specificity^56^, and explicit conduction delays^57^. Furthermore, our use of a continuous-valued ESN reservoir, rather than a spiking neuron model, ensures that the observed functional signatures are strictly attributed to the structural prior rather than the intricacies of cellular-level tuning or synaptic plasticity^58^.

This parsimonious approach allows us to delineate a modular blueprint for future inquiry. They serve as a foundational scaffold onto which additional biological factors can be systematically integrated as incremental constraints to quantitatively measure their specific contributions to computational gain. For instance: (1) Circuit Specialization: Incorporating cell-type and laminar specificity will clarify how local computation is shaped by cortical organization. (2) Temporal Coordination: Introducing explicit conduction delays and spiking dynamics will be critical for explaining inter-areal information routing and rhythmic synchronization. (3) Functional Reorganization: Integrating structural rewiring and plasticity mechanisms as testable rule-level elements will provide an operational path for predicting recovery after injury or learning-driven adaptation.

Ultimately, this methodology enables a rigorous causal decoupling of brain organization principles. By establishing a minimal sufficient structure for computational intelligence, we provide the necessary control to isolate the functional value of advanced biological features as they are layered onto this foundational protocol. This study thus transitions the field from fitting observed neural activity to deriving the underlying physical logic of neural information processing.

## Methods

### Neuronal-scale geometric scaling law of distance-connection probability

We used the MICrONS synapse-level connectome from mouse V1. The synapse table provides pre- and postsynaptic neuron IDs and 3D synapse coordinates in the Neuroglancer voxel system (voxel size 4 × 4 × 40 nm). For each neuron, we converted associated synapse coordinates to physical units and used the centroid as the neuronal coordinate. We computed centroid-to-centroid Euclidean distances *D*_ij_ for ordered neuron pairs (*i, j*). We defined *A*_ij_ = 1 if a synapse from neuron *i* to neuron *j* was present in the dataset and *A*_ij_ = 0 otherwise; self-loops were excluded (Fig. 2a).

For visualisation, we binned distances into logarithmically spaced intervals and computed *p*(*d*) in each bin as the number of connected pairs divided by the number of eligible pairs^10^. Model comparison was conducted on the unbinned pairwise data. We fit four candidate distance-decay families (power law, exponential, Gaussian kernel and logistic) by maximum likelihood and selected models using AIC, BIC and cross-validated log-likelihood. Finite sampling at long distances reduces the number of eligible pairs *N*_K_, pushing *p*(*d*) toward the detection limit 1/*N*_k_; we therefore repeated the analysis across subsampling scales to test robustness to long-distance sparsity.

### NIGC generative model

We used the MICrONS synapse-level dataset from mouse V1 as the structural reference. We sampled approximately 3×10^4^ neurons within the V1 microcircuit to construct an empirical binary adjacency matrix *A*. Given a set of neuron IDs, we queried synaptic connectivity via the MICrONS CAVEclient interface^59^. We represented each neuron’s 3D position by the centroid of its associated synapse locations, yielding a pairwise Euclidean distance matrix *D*.

In the generative model, NIGC initialized connection probabilities using a power-law distance prior, *p*_base_(*d*)∝*d*^-*α*^. We then introduced node propensity modulation to induce degree heterogeneity and imposed a global energy budget to constrain overall wiring cost. We sampled the probabilistic graph and pruned edges to obtain a directed binary connectome.

### Structural sufficiency test

Structural similarity was evaluated by comparing degree and local clustering coefficient distributions between empirical and generated networks. Distributional differences were quantified using the KS statistic, Jensen-Shannon (JS) divergence and Bray-Curtis dissimilarity.

To test which constraints drove structural similarity, we constructed baseline generative models on the same node set with matched key statistics, including economical wiring, homophily and a fully random baseline (matched for mean degree). Within the NIGC framework, we also removed key constraints one at a time and performed parallel comparisons against the full model using the same metrics, thereby quantifying each constraint’s contribution to structural agreement.

### Network construction and task validation

We constructed neuronal populations across 13 auditory-pathway-related regions (CN, Pons, ACx, IC, HPC, MGB, TRN, LP, SP, OFC, IL, PL and FP) using whole-brain 3D neuronal coordinates released by the Blue Brain Project^60^. Because the original coordinate set is large, we uniformly subsampled within each region to obtain a total of N = 3×10^4^ neurons^61^, a scale that remained computationally tractable for distance-dependent wiring, network-statistics evaluation and reservoir simulations under our GPU memory budget^54^, while matching the neuronal-scale regime used throughout the study. We then used the subsampled coordinates as inputs to NIGC to generate a directed neuronal-scale connectome.

To validate task executability and functional sufficiency under a unified dynamical framework, we converted the generated binary adjacency matrix into a weighted reservoir matrix. Existing edges were assigned random weights, and reciprocal connections were strengthened^62^. We then rescaled the reservoir matrix to satisfy the ESN echo-state property by setting its spectral radius^63^. We used CN as input nodes and prefrontal-related regions (IL, PL and OFC) as readout nodes. Internal reservoir connectivity was fixed, and only a linear readout layer was trained.

Task validation used the Spoken Arabic Digit classification dataset. Inputs were 13-dimensional mel-frequency cepstral coefficients (MFCCs), zero-padded to a fixed length. Training and test sets were split with speaker stratification. Model performance was evaluated by test-set classification accuracy. For task-level benchmarks, we also implemented end-to-end sequential and graph-based models (for example, LSTM, CNN, Transformer and GNN) under matched parameter budgets and compared their performance with NIGC-ESN.

### Functional phenotyping and validation

To assess whether the generative connectome supports multidimensional functional phenotypes under a unified ESN regime, we recorded task-driven time series of reservoir neuronal states while keeping reservoir weights and hyperparameters fixed. All functional metrics were defined on region-level signals, and the same analysis pipeline was applied in parallel to NIGC, the three structural baselines and each constraint-ablation condition.

LFP spectral fingerprints. For each region, we averaged neuronal states across neurons at each time step to obtain a region-level LFP time series. We band-pass filtered the LFP into five canonical bands (δ, θ, α, β and γ), computed band power as mean-squared energy, and normalized across bands to yield a relative-power vector.

Hierarchical delays. To characterize propagation order along the pathway, we used the peak absolute LFP amplitude within the analysis window as a reference. Onset latency was defined as the first time the absolute LFP reached 10% of this peak, and rise-to-peak latency as the first time it reached 90%^28^, to reduce sensitivity to initial offsets and local secondary peaks.

Cross-regional causality and phase coupling. Directional interactions were quantified using multivariate Granger causality. Under a vector autoregressive framework, we compared residual variances of restricted and full models and defined causal strength as the log variance ratio (negative values were set to 0)^64^. Phase coupling was quantified using the phase-locking value (PLV): after band-pass filtering the LFP to a target band, we extracted instantaneous phase via the Hilbert transform, computed phase differences and took the magnitude of the time-averaged unit phase-difference vector^65^.

Neuronal population spatiotemporal trajectories. For each region, we applied Uniform Manifold Approximation and Projection (UMAP)^66^ to the neuronal state matrix for nonlinear dimensionality reduction and connected embeddings across successive time steps to form a three-dimensional (3D) spatiotemporal trajectory. Within a local sliding window, displacement vectors approximated the trajectory tangent; we then computed the angle between trajectory directions for each pair of regions and their orthogonality (1-|cos *θ*|) to quantify cross-regional trajectory-level alignment and separability.

Pathological perturbations and structure-function coupling. In perturbation simulations, we uniformly rescaled weights of edges incident to a target region by a specified attenuation factor while keeping the rest of the network structure and dynamics unchanged, and compared relative changes in functional metrics before versus after perturbation. To quantify coupling between structural projections and evoked responses, we reconstructed a cross-regional weighted adjacency matrix from neuronal-scale reservoir weights and computed a structural projection field on the region graph using a k-step random walk^67^. We obtained the functional response matrix by delivering a brief pulse stimulus to each source region and defining each target region’s response as the time integral of the absolute LFP deviation from baseline within the stimulation window^68^. Pearson and Spearman correlations quantified coupling strength between the structural projection matrix and the response matrix^69^.

## Funding

S.G. acknowledges support from the National Natural Science Foundation of China (62571016). Y.P. acknowledges support from the National Natural Science Foundation of China (82572934). Y.W. acknowledges support from the National Key R&D Program of China (2023YFB3208002), the National Natural Science Foundation of China (82571767), and the Scientific Research Foundation of the National Health Commission/Zhejiang Provincial Major Science and Technology Plan Project of Medicine and Health (WKJ-ZJ2412).

## Author contributions

S.G., Y.W. and G.W. conceived the study and jointly designed the overall research framework.

G.W. curated and processed the neuronal-scale mouse connectome and spatial coordinate data, quantified and validated the power-law scaling of the distance-connection probability relationship, designed and implemented the NIGC framework, performed structural statistics benchmarking, constructed NIGC-based ESN reservoirs, and carried out functional simulations and consistency comparisons with experimental findings.

K.L. curated and preprocessed the visual cognition task dataset and conducted the corresponding cross-modal functional simulations and biological-consistency validation.

L.Q. and G.W. prepared, refined, and assembled all figures.

S.G., Y.W., Y.M. and X.G. provided overall guidance and supervision of the research.

G.W., S.G. and K.L. wrote the initial draft of the manuscript.

S.G., Y.W., X.G., C.T., X.C., Y.M., L.G.O., A.N., N.W., Y.P. and P.S. reviewed and edited the manuscript.

All authors discussed the results and their implications and approved the final version of the manuscript.

## Competing interests

The authors declare no competing financial interests.

## Data availability

The MICrONS neuronal-scale synapse table used in this study (Minnie65 Synapse Graph; data release v1300) is publicly available at https://bossdb-open-data.s3.amazonaws.com/iarpa_microns/minnie/minnie65/synapse_graph/synapses_pni_2.csv. Neuronal 3D coordinate data were downloaded from the Blue Brain Cell Atlas (2023 second version) at https://bbp.epfl.ch/nexus/cell-atlas/. The Spoken Arabic Digit classification dataset is available from the UCI Machine Learning Repository (https://doi.org/10.24432/C52C9Q). The custom video classification dataset used for the visual cognitive task is publicly available on ScienceDB at https://doi.org/10.57760/sciencedb.28916. Processed data and additional materials generated in this study are publicly available on Figshare at https://doi.org/10.6084/m9.figshare.31471300.

## Code availability

Custom code implementing the NIGC framework and all downstream analyses is publicly available on GitHub at https://github.com/jiedia/NIGC.

## Notes

https://bossdb-open-data.s3.amazonaws.com/iarpa_microns/minnie/minnie65/synapse_graph/synapses_pni_2.csv

https://bbp.epfl.ch/nexus/cell-atlas/

### Competing Interest Statement

The authors have declared no competing interest.

### Summary of Updates

The author list and author order have been updated. The scientific content, figures, and main text of the manuscript remain entirely unchanged.

