## supplementary information for "Minimal biophysical rules are sufficient for the emergence of computational intelligence at the neuronal scale"

### S1. Structural validation with baselines and constraint ablations

S1.1. Distance–connection geometric scaling law

To quantify the neuronal-scale distance–connection probability relationship, we fitted four standard distance-decay families (power law[1], exponential[2], Gaussian kernel[3] and logistic[4]) over a common distance range:

(1) Power law model:

$$p_k = \beta d_k^{-\alpha} \tag{S1}$$

(2) Exponential model:

$$p_k = \beta \exp\left(-\frac{d_k}{\lambda}\right) \tag{S2}$$

(3) Gaussian kernel model:

$$p_k = \beta \exp\left(-\frac{d_k^2}{2\sigma^2}\right) \tag{S3}$$

(4) Logistic model:

$$p_k = \frac{1}{1+\exp\left(\gamma_0 + \gamma_1 d_k\right)} \tag{S4}$$

Here, $d_k$ is the centre of distance bin $k$; $\alpha$, $\beta$, $\lambda$, $\sigma$, $\gamma_0$ and $\gamma_1$ were estimated from data. We fit each family by maximum likelihood to the unbinned pairwise data ($d_{ij}$, $A_{ij}$), using the same distance range across models, instead of fitting a local regression to the binned curve. This ensures that models are compared under the same data and statistical assumptions.

The fitting and evaluation metrics for the four models are summarized in Supplementary Table 1. Across metrics ($R^2$, log-likelihood, AIC[5] and BIC[6]) , the power law model performed best among the four families. On log–log axes, the power-law fit was best; it also maximized log-likelihood and minimized AIC and BIC after accounting for model complexity. Because information criteria penalize model complexity, this advantage is unlikely to reflect parameter count alone. Instead, information-criterion comparisons under the same data constraints consistently favored the power-law family. Based on these results, we used the power-law family as the geometric prior for the distance–connection probability relationship in subsequent NIGC analyses.

In Fig. 2c of the main text, estimates at the largest distances become sparse and deviate downward, which can resemble a model mismatch if not interpreted in light of sampling limits. To clarify this point, we computed the number of available neuron pairs $N_k$ in each distance bin and the corresponding empirical probability $p_k$. As distance increases, sampling-volume boundaries sharply reduce the number of eligible pairs; in the most distant bins, $N_k$ can be on the order of tens. In this regime, the minimum resolvable probability increment is ~$1/N_k$, comparable to (or larger than) the expected true probability. In other words, these far-distance bins approach the observability limit, and even small random fluctuations can push estimated probabilities in these bins below the values predicted by the power-law fit. Repeating the analysis under different subsampling scales (Fig. 2d) shows that the fitted power-law slope and agreement over short-to-intermediate distances are essentially unchanged. Deviations remain confined to the farthest bins, where sampling is sparse. Therefore, the long-distance anomaly in Fig. 2c is best explained by observability limits induced by volume boundaries and finite sampling, rather than a systematic failure of the power-law family.

To determine the exponent $\alpha$ of the power-law geometric prior, we generated connectomes across candidate values subject to fixed mean degree and energy budget constraints. We then selected the $\alpha$ that yielded the maximum information entropy. This maximization serves as a principled criterion for parameter selection under resource constraints[1,7], consistent with maximum-entropy arguments in neural wiring studies.

Using the V1 geometry, we matched mean degree to the measured connectome and scanned $\alpha$ from 0 to 2 in steps of 0.1. The resulting entropy–$\alpha$ curve is shown in Supplementary Fig. 1a. The curve peaks at $\alpha\approx0.5$ (entropy $\approx$6.27), compared with 5.58 at $\alpha$=0. Entropy then decreases rapidly with $\alpha$, approaching 0 when $\alpha\geq1$. When $\alpha$ is small, long-range edges dominate and the network mixes more uniformly; when $\alpha$ is large, edges concentrate locally and the network approaches a lattice. Intermediate $\alpha$ values maximize structural diversity and entropy. Based on this observation, we used $\alpha$=0.5 as the power-law exponent for V1 generation experiments.

For network construction along the mouse auditory pathway, we similarly scanned $\alpha$ from 0 to 1 in steps of 0.05 and obtained the entropy–$\alpha$ curve for the auditory-pathway connectome (Supplementary Fig. 1b). In this pathway, the information entropy increases with $\alpha$, reaches a peak at $\alpha\approx0.7$ (entropy $\approx$7.18), and then decreases gradually. Compared with single-region V1, the peak shifts to a larger $\alpha$, possibly because, under multi-region coupling, a shallow power-law decay makes connectivity more uniform at the whole-brain scale and reduces inter-regional structural differences. Under the same energy budget, a moderately steeper exponent increases local connection density while retaining a small number of long-range edges, yielding a topology that is locally clustered yet bridged across regions. Taken together, we used $\alpha$=0.7 as the power-law exponent in the auditory-pathway NIGC–ESN model.

#### S1.2. Structural baseline comparisons

We benchmarked NIGC against three baselines with matched node set and mean degree: economical wiring[8], homophily[9] and a mean-degree-matched random network. These baselines used the same node set and mean degree as NIGC and differed only in wiring rules and encoded constraints, providing reference models for assessing the contribution of the NIGC constraint set to structural agreement.

For degree distributions, NIGC matched the measured connectome closely in degree range and tail slope, with a mild compression in the fraction of the highest-degree nodes. By contrast, economical wiring and homophily concentrated degrees within a narrow range and reduced the heterogeneity seen in the measured network, which contains a small number of hub-like high-degree nodes and many low-degree nodes. The fully random network showed a near-Poisson degree distribution, consistent with an Erdős–Rényi graph[10]. Supplementary Table 2 quantifies these differences:

NIGC had the highest cosine similarity to the measured network and the smallest Jensen–Shannon (JS) divergence[11] and Bray–Curtis dissimilarity[12], whereas all three baselines showed larger discrepancies across metrics. Under matched mean degree, these results suggest that the full NIGC constraint set best recovers both the heavy-tailed degree profile and the empirical degree range among the models tested.

Comparisons of clustering-coefficient distributions further highlighted systematic differences among the baselines. The NIGC-generated network matched the measured connectome in the overall shape and tail behaviour of node-wise clustering coefficients. It neither produced the near-zero clustering mass typical of random graphs nor the uniformly elevated clustering of overly regular networks. Across the KS statistic[13], Jensen–Shannon (JS) divergence and Bray–Curtis dissimilarity (Supplementary Table 3), NIGC showed the smallest discrepancy from the measured network, whereas the KS statistic increased for economical wiring and homophily. This likely reflects their tendency to generate excessive local closure: abundant triadic closure and modular structure shift the clustering-coefficient distribution upward, reducing the proportion of medium-to-low clustering nodes observed in the measured connectome. The fully random network can appear closer under some divergence metrics, but its clustering coefficients concentrate in a narrow range near zero, which is inconsistent with the clustering heterogeneity of the measured network. Thus, this apparent closeness largely reflects peaked distributions under certain metrics rather than broad agreement in clustering structure.

Taken together, degree and clustering statistics indicate that each baseline captures a subset of structural features of the measured network. Economical wiring and homophily tend to increase local clustering while reducing degree heterogeneity, whereas the fully random network lacks a heavy-tailed degree distribution and shows minimal local closure. By contrast, under joint constraints (power-law geometry, node propensity, energy budget and entropy-based parameter selection), NIGC reproduces a heavy-tailed degree profile and a moderate clustering level comparable to the measured network. Across metrics, NIGC yields the smallest overall discrepancies among the tested models. These results suggest that nearest-neighbour wiring or node-type-similarity rules alone do not reproduce the joint degree–clustering profile of the measured microcircuit under our settings; adding geometry- and resource-related constraints, as in NIGC, improves agreement across both statistics.

In Fig. 3b, NIGC and the measured network show similar overall degree distributions, but a small separation remains at high degrees. The measured network shows a slightly higher frequency of high-degree nodes and correspondingly lower mass at medium-to-low degrees. To determine whether this difference reflects modelling limitations or sampling bias, we assessed the impact of volume boundaries and finite sampling on degree estimation. Specifically, keeping geometry and constraints unchanged, we subsampled both networks at 10k, 20k and 30k nodes and compared the resulting degree distributions (Supplementary Fig. 1i). Across sampling sizes, NIGC showed only small deviations from the corresponding measured subsamples, whereas the measured distributions consistently retained higher mass at high degrees and lower mass at medium-to-low degrees. This pattern is consistent with sampling-volume truncation: volume boundaries and finite sampling can preferentially reduce the contribution of medium-to-low degree nodes after normalization, thereby increasing the relative high-degree fraction. Therefore, the mild separation at high degrees is consistent with sampling effects in the measured microcircuit (finite-volume boundaries and finite sampling), and does not on its own indicate a systematic failure of the NIGC constraint set under our settings.

S1.3. Structural constraint ablations

Beyond baseline comparisons, we performed one-by-one ablations of key NIGC constraints using the same node set and matched mean degree. Keeping the power-law geometry fixed, we removed node propensity modulation, the global energy budget, and entropy-based parameter selection in turn, and regenerated connectomes. We then evaluated similarity to the measured V1 microcircuit using degree and clustering-coefficient distributions with the same metrics as above (Supplementary Fig. 1c–h). Corresponding cosine similarity, Jensen–Shannon (JS) divergence, Bray–Curtis dissimilarity, and KS statistics are summarized in Supplementary Tables 4 and 5.

For degree statistics, the full NIGC model achieved cosine similarity close to 1 and the smallest JS and Bray–Curtis dissimilarity relative to the measured network, indicating very similar degree range and tail shape. Removing node propensity modulation led to a marked decrease in agreement: cosine similarity dropped from near 1 to ≈0.3, while JS and Bray–Curtis increased several-fold. The heavy tail diminished and the distribution contracted towards intermediate degrees, producing a narrow-peaked profile. In contrast, removing the energy budget constraint had a smaller direct impact on the degree distribution, with metrics remaining largely unchanged. Removing the entropy-based parameter-selection constraint produced the largest change: the degree distribution largely lost its heavy-tailed profile, cosine similarity fell toward 0, and both JS and Bray–Curtis increased toward 1. The resulting curve formed a sharp unimodal peak around intermediate degrees, lacking both high-degree hubs and the broad low-degree mass. Together, these results suggest that node propensity modulation is a major contributor to degree heterogeneity. Entropy-based parameter selection helps set the power-law exponent at an intermediate regime that balances short- and long-range edges; removing either constraint drives the degree distribution towards a narrower, more homogeneous profile.

Clustering-coefficient distributions showed a complementary pattern under ablations. Under the full NIGC model, the clustering-coefficient distribution matched the measured network across the range and yielded the smallest KS statistic and JS among conditions. It preserved a large population of medium-to-low clustering nodes while maintaining a small subset of highly clustered local motifs. After removing node propensity modulation, clustering-related metrics worsened only mildly, suggesting that this constraint mainly shapes degree heterogeneity and has a limited effect on local triadic closure. By contrast, removing the energy budget substantially reduced clustering structure: the KS statistic, JS and Bray–Curtis dissimilarity increased, and the clustering-coefficient distribution shifted toward a narrow peak close to that of a random network, with strongly reduced triadic closure. This suggests that the global energy budget constrains wiring length and total edge count and influences the balance between local closure and longer-range edges. When entropy-based parameter selection was removed, the clustering distribution also deviated from the measured network. Although some metrics degraded less than in the energy-ablation case, the distribution became a narrow unimodal peak. Together with the homogenized degree distribution, this is consistent with a quasi-regular structure with intermediate degrees and constrained clustering values.

Taken together, degree- and clustering-based statistics indicate that these constraints play distinct yet complementary structural roles. Node propensity modulation shapes degree heterogeneity; the global energy budget influences the balance between local closure and longer-range edges; and maximum-entropy selection favours power-law parameters associated with higher structural diversity under matched resource constraints. When removed one at a time, different statistics shift away from the empirical range and towards random-like or more regular-like extremes. Thus, the geometric and related constraints in NIGC are complementary under our evaluation, forming an empirically compact constraint set. When jointly imposed, these constraints yield the closest agreement with the measured microcircuit in both heavy-tailed degree structure and moderate clustering, providing the structural substrate used in subsequent task- and function-level analyses.

### S2. Auditory-task functional validation

#### S2.1. Task-performance comparison under parameter-budget alignment

To quantify the performance–efficiency trade-off of NIGC–ESN on spoken-digit classification, we performed two budget-aligned comparisons on the Spoken Arabic Digit dataset[14]. The dataset comprises 10 digit classes. Each sample is represented as a 13-dimensional sequence of mel-frequency cepstral coefficients (MFCCs) with a maximum length of 93 frames. We zero-padded all sequences to 93 frames and used a speaker-stratified split with 660 training and 220 test samples. All models used the same preprocessed MFCC inputs and were trained for 200 epochs with multiclass cross-entropy and Adam. Optimization hyperparameters were held fixed across models, except for architecture settings controlling width (and thus parameter count).

In the first comparison, we matched the trainable-parameter budget to the NIGC–ESN readout and compared performance across models. In NIGC–ESN, the only trainable parameters are the linear readout weights and biases ($\approx 10^4$ parameters) applied to the prefrontal readout nodes. We tuned the hidden-state size and depth of the long short-term memory network (LSTM) to match this budget; we similarly adjusted the channel width of the convolutional neural network (CNN), the number of attention heads of the Transformer, and the graph-convolution width of the graph neural network (GNN). For each model, we ran 20 independent trainings with the same data split and recorded test accuracy and wall-clock training time. Under the trainable-parameter budget (Fig. 4c; Supplementary Table 6), NIGC–ESN achieved a mean test accuracy of $0.898 \pm 0.009$, whereas LSTM, CNN, Transformer and GNN achieved $0.419 \pm 0.119$, $0.818 \pm 0.026$, $0.749 \pm 0.043$, and $0.892 \pm 0.008$, respectively. The corresponding mean wall-clock training times were 18.6 s (NIGC–ESN), 21.8 s (LSTM), 61.0 s (CNN), 44.2 s (Transformer) and 69.6 s (GNN). Under this budget, NIGC–ESN achieved the highest mean test accuracy while requiring less training time than the CNN-, Transformer- and GNN-based baselines.

In the second comparison, we evaluated performance under a matched total-parameter budget, allowing end-to-end models parameter counts comparable to NIGC–ESN. We counted the fixed ESN reservoir weights toward the parameter budget and defined the effective parameter count of NIGC–ESN as the sum of non-trainable reservoir parameters and trainable readout parameters. We then rescaled the model width of each end-to-end baseline so that its total parameter count matched this budget. Under this configuration, NIGC–ESN trained only the same linear readout, whereas the LSTM, CNN, Transformer and GNN were optimised end to end. Across 20 repeats, NIGC–ESN maintained a test accuracy of $\approx 0.898$, comparable to the CNN, Transformer and GNN at this budget, while requiring less training time (Supplementary Fig. 2a; Supplementary Table 7).

To summarize accuracy relative to runtime, we defined an accuracy-per-second score as the average test accuracy divided by the average runtime[15] across the 20 repeats:

$$\text{score} = \frac{\overline{Acc_{test}}}{T} \tag{S5}$$

This score reports the mean test accuracy achieved per second of wall-clock training time. Under the trainable-parameter budget, the NIGC–ESN score was 0.048, whereas the LSTM, CNN, Transformer and GNN scores were 0.019, 0.013, 0.017 and 0.013, respectively. Under the total-parameter budget, the NIGC–ESN score remained 0.048; CNN, Transformer and GNN scored 0.016, 0.006 and 0.010, and LSTM scored 0.0008. Thus, under both budget definitions, NIGC–ESN achieved the highest accuracy-per-second score (Fig. 4c).

Together, these budget-aligned comparisons show that at a matched trainable-parameter budget, NIGC–ESN achieves higher test accuracy and shorter training time than the end-to-end baselines. When the fixed reservoir is also counted toward a total-parameter budget, end-to-end models can approach similar accuracy but at higher training-time cost. These results indicate that an NIGC-generated neuronal-scale connectome supports high task accuracy while improving the accuracy–speed trade-off under constrained computational resources.

S2.2. Functional baseline comparisons

We performed all functional analyses within the auditory-pathway NIGC–ESN architecture and compared them with three structural baselines. Specifically, we kept region-wise neuron counts, spatial coordinates, input/output mapping and the ESN update rule fixed, and replaced the NIGC-generated neuronal-scale connectome with each baseline topology (economical wiring, homophily and fully random). All other dynamical settings (input weights, activation function and readout architecture) were kept identical to those of NIGC–ESN. This design tests, under the same fixed, unified dynamical regime, how structural priors shape task performance and multidimensional functional phenotypes (Supplementary Figs. 3 and 4).

For task performance, we conducted independent hyperparameter scans for the three baselines. Under the same task setup and network scale as NIGC–ESN, we performed a grid search over ESN hyperparameters (including spectral radius and leak rate) and selected, for each baseline, the configuration with the best validation performance. We then trained and tested each baseline on the Spoken Arabic Digit dataset under its best configuration and compared it with the best NIGC–ESN setting. Even under their respective optimal spectral radius and leak rate, the economical wiring, homophily and fully random baselines achieved test accuracies of 0.450, 0.385 and 0.572, respectively, compared with 0.900 for NIGC–ESN. The three baselines and NIGC–ESN used the same reservoir size, readout structure and number of training epochs, leading to comparable wall-clock training time per run. The remaining performance differences therefore reflect differences in reservoir topology under matched training conditions.

For spectral fingerprints, the three baselines altered several region-specific patterns observed in NIGC–ESN. Using relative band power of regional local field potentials (LFPs), NIGC–ESN reproduced region-specific patterns in IC, CN and OFC reported previously. In the economical wiring and homophily networks, these patterns were attenuated: $\delta$ power in IC and OFC decreased, and the economical wiring model also showed reduced $\gamma$ power in CN. These spectral shifts are consistent with reduced low-frequency ($\delta$) components in IC/OFC and reduced high-frequency ($\gamma$) components in CN under the same task drive. In NIGC–ESN, the $\theta$ component in HPC was relatively weak under this spoken-digit classification task. Although $\theta/\gamma$ rhythms are often associated with hippocampal activity[16], this task does not explicitly require navigation or spatial memory[17]. The absence of a prominent θ peak in this setting may therefore reflect task demands rather than a generic limitation of the model.

For hierarchical delays, the fully random network showed a flattened temporal order: most regions responded within <10 ms after stimulus onset, and thalamic relays along the ascending pathway (for example, MGB), as well as HPC, peaked later than PFC. Consequently, the CN→IC→MGB→ACx→PFC delay ordering was not preserved in the fully random network, consistent with a loss of staged propagation under matched dynamics. In the homophily network, TRN showed minimal activation throughout the task, with LFP amplitude and power near the noise floor. This contrasts with its established involvement in thalamocortical gating and inhibition of MGB[18]. In NIGC–ESN, TRN showed an early onset and a short rise-to-near-peak time (onset ≈11 ms; near-peak ≈30 ms), in line with thalamocortical gating[19]. PFC regions peaked later, consistent with delayed recruitment of higher-order regions under the same task drive[20]. Together with the β-dominant pattern in prefrontal regions reported in the main text, these latency patterns provide a consistent cross-metric comparison across spectrum and timing.

For cross-regional causal interactions and phase coupling, the three baselines showed markedly different patterns from NIGC–ESN under matched dynamics. From regional LFP time series, we constructed multivariate Granger-causality matrices and computed the phase-locking value (PLV) between region pairs (within the task-relevant frequency bands). Relative to NIGC–ESN, which showed a structured feedforward–feedback pattern along CN–IC–MGB–ACx–PFC and prominent β/γ coupling between PFC and ACx, the economical wiring and homophily networks yielded sparse Granger-causality matrices, with near-zero causal strengths for many region pairs. This contrasts with reports of broad task-related activity distributed along the CN–IC–MGB–ACx–PFC pathway[21]. By comparison, the fully random network produced a "hotspot"-like Granger pattern, with strong links scattered across region pairs without a clear pathway organization. PLV analysis provided a complementary pattern. In the economical wiring and homophily networks, ACx–PFC PLV decreased and PFC-internal pairs did not show stable high β-band coupling. In the fully random network, PLV increased broadly across many region pairs, whereas the concentrated within-PFC coupling seen in NIGC–ESN was reduced. Overall, without the NIGC structural priors, the baselines tended toward either weak inter-regional coupling or broadly distributed phase locking—patterns that differ from hierarchical rhythmic organization[22].

For low-dimensional spatiotemporal trajectories, we embedded single-region population activity into a three-dimensional space using Uniform Manifold Approximation and Projection (UMAP) and compared trajectory geometry over the task timeline. Relative to the ring-like or "ramping–turning" trajectories observed in NIGC–ESN (qualitatively similar to prior reports), all three baselines showed altered trajectory geometry. In the economical wiring baseline, ACx, PL and OFC trajectories were compressed into a narrower region with reduced spatial extent and curvature. Late and early time points were closer in the embedding, and temporal progression was less geometrically separated. In the homophily baseline, some time points collapsed into small clusters and trajectories became fragmented into sparse point sequences over certain periods, reducing continuity across consecutive task epochs. In the fully random baseline, trajectories showed increased jitter and reversals, and the activity cloud was more diffuse, with early and late time points more intermingled and reduced time-ordered structure. Together, these results show that, under identical dynamical rules, changing reservoir topology alone can markedly alter the low-dimensional organization of population activity, yielding more compressed, fragmented or diffuse trajectories.

For pathological perturbations, we used the same hippocampal (HPC) attenuation protocol as in the main text and examined responses over progressively stronger attenuation levels. Specifically, we attenuated the weights of edges incident to HPC (incoming and outgoing; 60–95% attenuation in 5% steps), re-ran the task at each level, and quantified changes in HPC LFP band power (δ, θ, α, β and γ) relative to the unperturbed condition. For NIGC–ESN, a consistent pattern emerged across the attenuation range: as intra-HPC connectivity was weakened, θ and γ power decreased, whereas δ power increased; α and β changed less. This δ increase together with θ/γ decreases is qualitatively consistent with trends reported in hippocampal lesion and injury studies and was stable across attenuation magnitudes. By contrast, the three baselines showed less stable spectral changes under the same attenuation protocol. In the economical wiring and homophily networks, δ and γ power sometimes increased together or δ decreased; the fully random network showed frequent sign changes in δ, θ and γ across attenuation levels. These results show that, with dynamical rules fixed, reservoir topology alone can substantially change the spectral response to perturbation, ranging from stable directionality to patterns that are less consistent with reported experimental trends.

Across task performance, spectral fingerprints, hierarchical delays, directed interactions/phase coupling, low-dimensional trajectories and perturbation responses, the baselines deviated from NIGC–ESN under matched ESN dynamics in one or more dimensions. Together, these comparisons indicate that the NIGC constraint set supports a more stable alignment across functional metrics under a fixed ESN regime.

#### S2.3. Functional constraint ablations

We conducted functional ablation comparisons within the auditory-pathway NIGC–ESN architecture. We ablated each core NIGC constraint—power-law distance prior, node propensity modulation, the global energy budget and maximum-entropy selection—one at a time and generated the corresponding ablated connectomes. Specifically, we kept neuron counts, 3D coordinates, input/output mapping and ESN dynamics fixed, and altered only the corresponding structural prior in the generator. For the geometric ablation, we replaced the power-law distance prior with an exponential distance-decay model commonly used in region-scale generative settings. For the other three ablations, we removed node-level propensity modulation, the global energy budget constraint, or the information-entropy maximisation constraint (Supplementary Figs. 5 and 6).

For task performance, we selected hyperparameters within the same spectral-radius and leak-rate search ranges used for the full NIGC–ESN and trained and tested each ablated model on the spoken-digit classification task. Relative to the NIGC–ESN baseline accuracy (≈0.900), accuracies under power-law geometry ablation, energy-budget ablation and entropy-maximisation ablation were 0.689, 0.474 and 0.711, respectively, whereas ablating node-level propensity modulation yielded 0.914. Reservoir size and the training protocol were held constant, and runtime differences were minimal. The performance changes therefore reflect differences in structural priors under matched training conditions.

For spectral fingerprints, all four ablations reduced δ-band power in the auditory relay nucleus IC, attenuating the δ/β coexistence pattern observed in NIGC–ESN. In prefrontal regions, removing either power-law geometry or node-level propensity modulation decreased δ power in OFC, making β/γ components relatively more prominent and altering a δ–β mixed pattern reported in auditory tasks. Removing either power-law geometry or the information-entropy maximisation constraint reduced γ power in IL, shifting IL toward a more low-frequency-dominated spectrum. Together, these results suggest that power-law geometry and statistical constraints contribute to region-specific spectral fingerprints in the auditory–prefrontal circuit, and that removing any one of them produces detectable changes.

Hierarchical-delay analysis further supported a prominent role of the energy budget in maintaining a plausible propagation order. When the global energy budget constraint was removed, long-range connections were no longer penalized, thalamic relays were effectively bypassed, and PFC peaks occurred earlier than IC/MGB; overall response onsets shifted earlier and the ascending hierarchy was largely flattened. By contrast, removing power-law geometry, node-level propensity modulation or information-entropy maximisation perturbed the delay structure while broadly preserving pathway order. This suggests that the energy budget suppresses long-distance shortcuts that can compress propagation timing, whereas the other constraints more strongly regulate local statistics and connection diversity.

Cross-regional causal interactions and phase coupling provided complementary evidence for constraint-specific effects. After removing power-law geometry or information-entropy maximisation, the Granger-causality matrix was sparser, long-range feedforward/feedback pathway strengths decreased, and PLV between ACx and PFC decreased, consistent with a weakened pathway backbone. Removing the global energy budget constraint produced a pattern resembling the fully random network: TRN received concentrated causal inputs, PLV increased broadly across region pairs with reduced regional specificity, and the network tended toward a globally coupled regime. By comparison, removing node-level connection propensity modulation mainly reduced intermediate-scale heterogeneity, yielding a more uniform causal graph.

At the level of low-dimensional spatiotemporal trajectories, each ablation produced systematic changes in trajectory geometry. When power-law geometry was removed, the ACx trajectory retained curvature but did not return to the initial neighbourhood, drifting along an open arc. PL and OFC trajectories became flatter, with a shortened late "rewrapping" segment. Separation among the three regions in embedding space also decreased. When node-level propensity modulation was removed, ACx and PL trajectories became nearly one-dimensional with minor kinks. In OFC, the late turning segment was compressed, and cross-regional trajectory angles decreased. When the global energy budget constraint was removed, the ACx trajectory deviated from the ring-like pattern, forming a monotonically extending open arc, and trajectory lengths in higher-order regions were shortened. When information-entropy maximisation was removed, trajectories in all three regions collapsed into short, nearly straight paths with increased overlap between early and late states. Overall, relative to NIGC–ESN, all four ablations showed shorter trajectories, weaker rewrapping and reduced cross-regional directional separation. The largest changes were observed after removing the global energy budget or information-entropy maximisation constraint.

For pathological perturbations, we examined how each constraint affects hippocampus-related spectral reconfiguration under the same attenuation protocol. Repeating the same attenuation experiment as in NIGC for all four ablated networks revealed that spectral reconfiguration became unstable: in some conditions, δ power increased excessively and even reversed at high attenuation levels; in others, θ and γ increased with attenuation, or fluctuated non-monotonically across attenuation ratios. α and β bands also exhibited enhancement patterns inconsistent with experimental trends. These results suggest that disrupting any single component among power-law geometry, node-level propensity modulation, the energy budget and information-entropy maximisation reduces the stability and interpretability of the lesion-response signature (δ increase with θ/γ decreases).

Taken together, the functional ablation experiments show that power-law geometry, node-level connection propensity modulation, the global energy budget and information-entropy maximisation jointly contribute to functional phenotypes under identical ESN dynamics. The most consistent alignment across task performance and multilevel functional metrics was observed when all four constraints were present. Removing any single constraint produced measurable changes in at least one class of functional metrics, broadly consistent with the constraint's role in the structural analyses.

S2.4. Intrinsic dynamical properties

Beyond the hierarchical-delay analysis, we examined the intrinsic timescales of the auditory-pathway NIGC–ESN in the absence of task-related inputs. Specifically, with the NIGC-generated neuronal-scale connectome and ESN parameters fixed, we injected small-amplitude Gaussian white noise (mean 0, s.d. = 0.02) into each reservoir neuron and let the network evolve for 2 s to approximate resting-state dynamics[23]. We then averaged neuronal activity within each region to obtain a region-level LFP time series[24] and computed the autocorrelation function and intrinsic timescale for each region.

For each region-level LFP, we computed the autocorrelation function and defined the intrinsic timescale, τ, as the integral of the positive-lag autocorrelation (area under the curve). We then normalized τ across regions to obtain relative timescales. Compared with approaches that estimate timescales using an exponential decay constant or the half-maximum width of the autocorrelation function[25], our definition is fitting-free and still captures relative differences in autocorrelation-decay rate.

Under this noise-driven resting-state condition, autocorrelation functions decayed smoothly from 1 at zero lag, with systematic differences in decay rate (Supplementary Fig. 2b,c). When ordered from short to long timescales, Pons, LP, and early auditory brainstem/relay regions (CN and IC) showed the fastest autocorrelation decay, with normalized timescales of 0.4–0.5. The auditory thalamic nucleus MGB and relay/gating nuclei (SP and TRN) were intermediate. By contrast, ACx, HPC, and the prefrontal-related regions IL and FP showed the slowest decays and the largest relative timescales, with ACx having the largest τ. Along CN→IC→MGB→ACx, timescales increased approximately monotonically. Within prefrontal cortex, IL and FP timescales were longer than those of relay/gating nuclei and early auditory regions. This gradient is consistent with the direction of the hierarchical delays along the same route under task conditions, suggesting a progression from fast sensory pathways to slower associative pathways under both noise-driven and task-driven regimes.

S2.5. Distribution of causal strengths and similarity metrics

We summarized Granger causality strengths among the 13 auditory-related regions and compiled a histogram of all directed edges (Supplementary Fig. 2d). The distribution is slightly left-skewed (mean ≈0.816; median ≈0.783): most edges are of moderate strength, with a small high-strength right tail. Right-tail edges were concentrated in task-relevant pairs (e.g., ACx–FP, CN–IC, CN–HPC), forming a strong-influence backbone, whereas the remaining edges contributed more diffuse moderate coupling. Overall, this pattern is consistent with a small number of strong directed interactions embedded in a broader background of moderate coupling[26].

In the rhythm analysis, the main text emphasizes PLV as summaries of cross-regional coupling. To further characterize rhythmic relationships, we computed a panel of metrics on the same LFP rhythmic signals—Pearson/Spearman correlation, cosine similarity, amplitude and derivative correlation, energy ratio, phase consistency and phase s.d., wavelet correlation and mutual information (Supplementary Fig. 7). These metrics provide complementary descriptions spanning linear and rank association, directional similarity, amplitude and rate coupling, energy asymmetry, phase stability, time–frequency covariation and nonlinear dependence.

Pearson, Spearman, and cosine similarity showed consistent patterns: FP, HPC, IC, IL, LP, and MGB tended to be mutually similar, whereas relationships with early relay structures (CN, SP, and Pons) were weaker or negative. In particular, FP–HPC, FP–IC, and HPC–IC showed stable positive association across all three metrics, consistent with a tightly coupled subset of task-evoked rhythms. By contrast, ACx was negatively associated with CN/IC/MGB but positively associated with PL/IL/Pons. Spearman and cosine metrics mirrored Pearson, suggesting that these relationships extend beyond linear covariation to monotonic and directional concordance.

Amplitude correlation was strongest among FP, HPC, IC, IL, LP, and MGB (e.g., ≈0.8–0.9 for FP–HPC, FP–IC, and HPC–IC), consistent with coordinated amplitude co-fluctuations. By contrast, amplitude correlations between Pons/SP/TRN and cortical or hippocampal regions were lower. Derivative correlation yielded a more diffuse pattern: a few pairs (e.g., FP–HPC and FP–IC) remained strongly positive, whereas many pairs were moderate or near zero, consistent with limited synchrony in rate-of-change across regions at the task timescale.

The energy-ratio matrix spanned a wide dynamic range (log scale), with a small number of pairs showing very large ratios. Overall, FP, HPC, and IC more often carried higher energy than their partners, whereas IL tended to be lower-energy across comparisons, yielding an asymmetric pattern. Phase consistency and phase standard deviation provided complementary views of phase stability. Pairs such as ACx–CN, ACx–IL, ACx–Pons, FP–HPC, and FP–IC showed higher phase consistency and lower phase s.d., consistent with relatively stable phase relationships over the task epoch. By contrast, pairs involving TRN and SP tended to show lower phase consistency and higher phase s.d., consistent with less stable phase locking in the analyzed bands.

Finally, wavelet correlation captured time–frequency covariation and broadly mirrored the linear similarity measures, with higher values concentrated among FP, HPC, IC, IL and LP and weaker covariation involving Pons, SP and TRN. Importantly, mutual information (MI) complemented this picture by revealing dependencies that need not be explained by linear correlation or phase synchrony alone: several pathways—including ACx–LP, IC–PFC and intra-PFC pairs—showed elevated MI alongside strong wavelet-based coupling, indicating that these interactions preserve tighter statistical dependence at the level of nonlinear structure and time–frequency coordination. Taken together, these complementary metrics align with the PLV/MI patterns emphasized in the main text, highlighting stronger rhythmic coordination within a subset of regions and weaker or more variable coupling involving brainstem and selected thalamic nuclei.

### S3. Low-dimensional representations and spatiotemporal trajectory geometry

#### S3.1. Dimensionality selection for PCA/UMAP

To construct interpretable spatiotemporal trajectories at the neuronal population level, we selected a low-dimensional embedding that retains the key structure of population activity. As a linear baseline, we ran principal component analysis (PCA) for each of the 13 regions and computed the cumulative explained-variance ratio to gauge effective linear dimensionality (Supplementary Fig. 8). Across regions, the first three principal components captured ≥95% of the PCA-explained variance, consistent with a pronounced low-rank structure in population states. Because population-state dynamics can be nonlinear and locally manifold-like[27], PCA is best viewed as a lower bound on effective dimensionality. We therefore used Uniform Manifold Approximation and Projection (UMAP)[28] and quantified local neighbourhood preservation under nonlinear embeddings.

We selected the UMAP dimensionality using neighbourhood-preservation criteria quantified by three complementary metrics. For each region, we embedded the high-dimensional neuronal states into target dimensions d=2–10. We then computed Trustworthiness, Continuity[29] and the k-nearest-neighbour (kNN) retention rate (overlap between high-dimensional and low-dimensional kNN sets)[30]. We tested sensitivity to neighbourhood size (k=15 or 30) and distance metric (Euclidean or cosine) by repeating the evaluation under each setting (Supplementary Fig. 9).

Under the default setting (Euclidean distance; k=30), all three metrics were consistently high across d=2–10 and saturated early. The cross-regional mean Trustworthiness ranged from 0.9849 to 0.9865, Continuity from 0.9899 to 0.9919, and kNN retention from 0.9240 to 0.9283. Beyond d=3, gains were negligible (maximum increase <0.001 for Trustworthiness and Continuity; maximum absolute change ≈0.003 for kNN retention), indicating a plateau. The same pattern held across regions: Trustworthiness and Continuity were tightly distributed, and increasing d did not systematically improve or stabilize the metrics (Supplementary Fig. 9). Thus, larger d primarily increases visualization complexity and downstream computational cost, with minimal gains in neighbourhood-preservation metrics.

These conclusions were robust to neighbourhood size and distance metric. With Euclidean distance and k=15, all three metrics plateaued after d=3 and were slightly higher in magnitude. With cosine distance (k=15 or 30), metrics were lower in magnitude than with Euclidean distance, but still saturated after d=3. Accordingly, we used a 3D UMAP embedding for all subsequent trajectory construction and cross-regional geometric analyses. This choice is consistent with PCA-based effective dimensionality and with the plateau in neighbourhood-preservation metrics, balancing fidelity, interpretability and computational cost.

#### S3.2. Spatiotemporal trajectory analysis

Using a 3D UMAP embedding, we projected each region's population states in temporal order and connected successive points to form continuous trajectories (Supplementary Fig. 10a). To reduce visualization bias from cross-regional scale differences, we rescaled each region's trajectory coordinates to [0, 1] using that region's minimum and maximum values. Accordingly, we analysed trajectory geometry, including linearity, curvature, looping and stage-like plateaus, rather than absolute position or axis semantics.

Overall, the trajectories exhibited two broad geometric motifs. Some regions (for example, TRN and FP) showed an approximately linear progression along a dominant direction, consistent with limited curvature. Others (for example, MGB, HPC and several subcortical nuclei) showed bends or loops, consistent with stage-like transitions in the low-dimensional state space. Because UMAP embeddings are invariant to global rotation/reflection, we interpret geometry (loops, bends and turns) rather than axis-specific trends.

The TRN trajectory formed a smooth, near-straight segment with minimal curvature and looping. This geometry suggests dominance of a single low-dimensional mode that tracks temporal progression. Given TRN's inhibitory gating role in thalamo–cortical circuits, low-dimensional modulation is compatible with its reported control over cortical state and thalamic transmission[31].

Compared with TRN, MGB showed an inflection and then progressed along a second direction, suggesting stage-like structure. The bend may reflect a change in dominant response components (e.g., onset versus offset), which reshapes the embedding dynamics. Reported ON/OFF pathway dissociation in auditory thalamus is compatible with separable onset and offset components in low-dimensional space[32]. Accordingly, the bend is consistent with a transition between two relatively orthogonal subspaces rather than a one-dimensional expansion–contraction.

Downstream regions (CN, Pons and SP) showed more curved trajectories, often with an initial decline/lateral shift and a late rise or turn. CN heterogeneity (cell types and coding mechanisms) may underlie more complex low-dimensional folds and stage-like segments[33]. The late jump in SP is compatible with an OFF component, as OFF responses often concentrate near stimulus termination[34]. By contrast, Pons showed larger rotations and multi-segment advances. This geometry is compatible with the multimodal modulatory role of brainstem circuits in arousal regulation and rapid transmission[35].

IC and LP showed a single large-amplitude bulge: a gradual rise, a high-curvature/high-amplitude segment, a partial return and stabilization near the endpoint. IC acts as a convergence/integration node in the ascending pathway and is sensitive to temporal-envelope processing, inhibition–excitation balance and acoustic-feature integration. Accordingly, this geometry is compatible with a transition from input-driven dynamics to a stabilized, integrated state[36]. LP showed a similar outward–return pattern, compatible with its cortico-thalamic integrative role in prediction-error processing and bidirectional top-down/bottom-up interactions[37].

Among regions, HPC showed the strongest tendency toward looping/closure and the longest smooth segment, resembling an annular or curved-tubular manifold. This geometry is compatible with low-dimensional coding in hippocampus during learning[38], where task variables can form structured manifolds and population activity can undergo representational decorrelation and reorganization. Here, the loop-like trajectory suggests reversible or quasi-periodic traversal on a manifold, rather than a strictly periodic oscillation.

Finally, IL and FP combined monotonic progression with a late turn or plateau, consistent with the ramp-to-turn geometry reported for prefrontal population dynamics[39]. In this interpretation, the smooth segment is consistent with temporal integration, whereas the terminal bend marks a transition to a readout/decision-related subspace.

Overall, Supplementary Fig. 10a summarizes region-specific dynamics: TRN/FP are dominated by a single mode, whereas bends/loops in MGB, HPC and several subcortical nuclei indicate stage structure and inter-subspace transitions. These patterns motivate the quantitative metrics used below and support the use of a 3D embedding that balances interpretability and neighbourhood-structure fidelity.

S3.3. Cross-regional trajectory geometry

Using a three-time-step sliding window, we quantified cross-regional trajectory alignment by the angle $\theta_{ij}(t)$ between local tangent vectors and the corresponding orthogonality metric, $O_{ij}(t)=1-|\cos\theta_{ij}(t)|\in[0, 1]$ (Fig. 6b,c).

Along CN→IC→ACx, early time points showed aligned progression within the chain: ACx–CN, CN–IC and ACx–IC had mean angles of ≈22–34° and orthogonality of ≈0.09–0.19. Together, these values indicate stronger cross-regional alignment of trajectory tangents during the initial stimulus-driven stage. Later time points showed increased divergence within CN→IC→ACx: ACx–CN and ACx–IC reached orthogonality ≈0.66 and ≈0.73, with angles ≥90°, indicating reduced alignment of trajectory directions after the early propagation stage.

At the whole-network level, the heat map showed time-resolved structure in cross-regional alignment and divergence. Early time points showed preferential divergence around CN and ACx: angles exceeded 90° for ≈2/3 of CN-related pairs and ≈1/2 of ACx-related pairs. By contrast, early alignment was stronger among FP/IL/PL and HPC (mean angles ≈24–36°, orthogonality ≈0.09–0.19), followed by later divergence (FP–HPC ≈110°, orthogonality ≈0.63). This temporal ordering is compatible with reports of longer intrinsic time scales in higher-order cortical areas[25].

S3.4. Time-window effects

We evaluated temporal-scale sensitivity by recomputing angle and orthogonality with different sliding-window lengths and comparing the resulting angle–orthogonality relationships across samples (Supplementary Fig. 10b).

With a 5-time-step window, the scatter closely followed the cosine-implied mapping between angle and orthogonality. The tighter clustering around the theoretical curve suggests that short windows better approximate local tangents and thus capture transient alignment changes with higher temporal resolution. Quantitatively, deviations from the theoretical mapping were reduced with the 5-time-step window (mean deviation ≈25% of that with the 25-time-step window).

With a longer window (25 time steps), the angle–orthogonality relationship became more dispersed. First, the dynamic ranges of both angle and orthogonality narrowed, with fewer extreme values. Second, orthogonality values were more variable at similar angles, consistent with longer windows averaging across multiple transient phases. As a result, directions estimated from longer concatenated segments can deviate from instantaneous tangents, reducing sensitivity to rapid turns in the trajectories. This is consistent with reports that decoding accuracy often peaks within transient millisecond-scale windows after stimulus onset[40]. Together, shorter windows are better suited to capturing rapid cross-regional coordination and decoupling during the stimulus-evoked transient phase, whereas longer windows provide a coarser summary of slower geometric variation at the cost of reduced transient sensitivity.

### S4. Visual-pathway connectome and video task

#### S4.1. Construction of the visual-pathway connectome

We constructed neuronal populations across 12 regions along the mouse visual pathway (VC, SC, LP, TRN, LGN, OPN, PPC, MC, PL, OFC, IL and DS[41–45]; abbreviations defined in Supplementary Fig. 14) using whole-brain 3D neuronal coordinates released by the Blue Brain Project[46]. Because the full coordinate set contains on the order of $10^7$ entries, we uniformly subsampled ≈0.1% of neurons within each region. The sampled coordinates were then scale-normalized and used as input to NIGC to generate a neuronal-scale visual-pathway connectome.

To support downstream tasks, we converted the generated binary adjacency matrix into a weighted reservoir matrix. Existing edges were assigned random weights (log-normally distributed), and reciprocal connections were strengthened relative to unidirectional connections to reflect reported microcircuit trends. To satisfy the ESN echo-state property (that is, to avoid divergence or excessive decay), we linearly rescaled the reservoir matrix by setting its spectral radius within a predefined target range, yielding the final reservoir weight matrix $W_{\mathrm{res}}$.

#### S4.2. Three-class video task for cross-modal generalization

To test cross-modal generalization, we switched from the auditory pathway to a three-class video task along the visual pathway and reapplied the same analysis pipeline. Consistent with the main text, we kept the reservoir connectivity fixed (no task-specific optimisation) and relied on the neuronal-scale topology generated under the same geometric and biophysical constraints. We then assessed task performance and used the task-driven regime for subsequent comparisons of basic functional readouts (spectral fingerprints, hierarchical delays and cross-regional coordination).

Specifically, we collected three action categories (“approaching”, “moving away” and “pausing/observing”), with 100 videos per category, and used a class-balanced split (80% training; 20% testing). We chose these actions because they differ in spatiotemporal cues (for example, optical-flow direction and speed distribution, changes in target scale and the intermittency of motion energy). This provides separable dynamic structure with a modest sample size while reducing confounds from complex semantics or backgrounds. For implementation, each video was converted into a frame sequence, and early-stage nodes (SC and LGN) served as input nodes. We then applied a linear readout to population states from higher-order cortical and downstream regions (PL, IL, OFC, PPC, DS and MC) for classification. This design enables a controlled comparison of task executability and functional signatures across pathways under the same ESN dynamics.

#### S4.3. Biologically inspired retina-like visual front-end and feature encoding

In modelling the visual pathway, we did not feed raw pixel intensities directly into the reservoir. Instead, we used a retina-like front end: each video frame was mapped to spatiotemporal responses that emulate retinal ganglion cells (RGCs), which were then projected to the superior colliculus (SC) and lateral geniculate nucleus (LGN) as inputs to the visual pathway. This choice was motivated by two considerations. Physiologically, SC and LGN do not receive raw pixel intensities; instead, they receive preprocessed retinal signals[47] shaped by classic operations such as centre–surround antagonism, ON/OFF pathway decomposition[48], multiscale integration and temporal high-pass filtering. From a modelling perspective, we designed the visual drive to approximate key statistical properties of mouse visual processing while avoiding additional trainable degrees of freedom. This places the primary biological-consistency burden on the generative connectome rather than on a task-specific front end.

We therefore used a bank of biologically inspired fixed filters (rather than a learnable convolutional front end or high-dimensional deep features) to approximate retinal spatiotemporal processing[49]. Relative to an end-to-end trained CNN, this fixed-filter scheme offers practical benefits for our evaluation. First, it restricts the front-end degrees of freedom to a small set of interpretable parameters (e.g., spatial scale, centre–surround ratio and a temporal constant), reducing label overfitting under limited sample sizes and limiting front-end learning as a confound in testing fixed geometric and biophysical constraints. Second, the filter forms map directly to classic retinal physiology (RGC receptive fields and transient/sustained response types). This facilitates qualitative comparisons of spectral fingerprints, hierarchical delays and cross-regional coordination at SC/LGN and higher cortex with electrophysiological findings, without an additional remapping from deep feature space. Finally, a fixed-filter front end improves methodological symmetry between visual and auditory pathways: both use a simple, task-agnostic feature mapping to drive the same NIGC–ESN dynamics, enabling more controlled comparisons across pathways.

In implementation, we converted each video frame to grayscale and downsampled it to a low-resolution retinal grid (e.g., $H \times W$). On this grid, we constructed a multiscale bank of centre–surround difference-of-Gaussians (DoG) filters to approximate RGC receptive-field spatial profiles[50]. For each spatial scale $s$, we defined a pair of Gaussian kernels with standard deviations $\sigma_{\mathrm{centre}}=s$ and $\sigma_{\mathrm{surround}}=\alpha s$ ($\alpha>1$ is a fixed ratio), and took their difference to obtain a DoG kernel. A positive DoG approximates an ON-centre/OFF-surround receptive field, whereas a negative DoG approximates an OFF-centre/ON-surround receptive field. Thus, at each spatial location we obtained multiscale paired ON/OFF filter responses, yielding a multiscale encoding of local luminance contrast.

Along the temporal dimension, we further applied a first-order temporal high-pass filter to the spatial-filter outputs. Specifically, the difference between spatial responses of adjacent frames approximated the transient component, whereas the original (or lightly smoothed) response provided the sustained component, capturing sensitivity to luminance changes and motion edges[51]. We then applied a biologically motivated nonlinear transform to each response: half-wave rectification (floor at 0), followed by a saturating nonlinearity (for example, tanh) to compress extreme values and yield a bounded, sparse, positively skewed firing-rate-like signal. This linear–nonlinear structure follows the classical LNP framework for sensory-neuron responses and better approximates RGC population firing patterns than purely linear filter outputs[52].

In terms of feature form, for a given video we ultimately obtained a 4D tensor $\mathbf{R}(t, x, y, c)$, where $t=1,\ldots, T$ indexes time steps (frames), $x=1,\ldots, H$ and $y=1,\ldots, W$ index retinal spatial positions, and $c=1,\ldots, C$ indexes combinations of spatial scales and ON/OFF channels. After flattening the spatial and channel dimensions, each $(x, y, c)$ triplet can be treated as a virtual RGC unit, whose temporal response defines its activity time course. Each video can then be represented as a feature matrix of size $T \times N_{\text{retina}}$, with $N_{\text{retina}} = H \times W \times C$. When coupling to NIGC–ESN, we defined a Gaussian receptive field over the retinal grid for each SC and LGN input neuron. We then computed a distance-weighted sum of virtual RGC responses within that field, constructing an effective transmission function from the retina to SC/LGN. This mapping preserves multiscale, ON/OFF and motion-sensitive properties while implementing a topographic projection with overlap and redundancy[53]. It therefore provides a biologically inspired input basis for simulating visual-stimulus-driven pathway dynamics on the generated connectome.

S4.4. Functional validation in the visual cognitive task

Using the same NIGC–ESN dynamical regime and analysis pipeline as in the auditory task, we assessed whether the visual-pathway connectome supports comparable functional signatures during the three-class video-classification task.

(1) Spectral fingerprints: band composition of multi-regional LFPs in the visual task

We decomposed local field potentials (LFPs) from key nodes (SC, LGN, VC, MC, IL and OFC) into band power (δ, θ, α, β and γ). Under this analysis window and LFP definition, task-related power was concentrated almost entirely in the β and γ bands (Supplementary Fig. 11b,c). Across regions, β and γ accounted for nearly all relative power, with δ/θ/α together contributing only a few per thousand. This pattern indicates a predominantly high-frequency regime under the current analysis window. Specifically, β/γ fractions were ≈0.76/0.24 in SC, ≈0.65/0.35 in LGN, and ≈0.60/0.39–0.40 in VC and MC; OFC showed a slight γ dominance ($\beta \approx 0.47$, $\gamma \approx 0.53$), whereas IL was strongly γ-dominant ($\beta \approx 0.10$, $\gamma \approx 0.90$). Given the short window, lower-frequency components contribute minimally under the present definition; Below we focus on β/γ fingerprints and their regional differentiation.

This β/γ-dominant pattern is broadly consistent with reports implicating β/γ oscillations in visual sensory processing and inter-areal coordination. Studies in mouse V1 suggest that local inhibitory microcircuits can modulate β and γ relatively independently and that visual stimulation or experience can enhance β/γ oscillations[54]. Simultaneous recordings from dLGN–V1 further indicate that narrow-band γ can be selectively enhanced during visual stimulation and may relate to thalamo–cortical information transfer[55]. In our simulations, both LGN and VC exhibited pronounced γ components, consistent with thalamo–cortical interactions accompanied by high-frequency enhancement. β also remained substantial in early entry regions (SC/LGN), suggesting that, under the present model and task configuration, the functional division of labor between β and γ may shift in a paradigm-dependent manner. We examine this possibility further with the directional-causality and phase-coupling analyses below.

High-frequency features comparable to prior reports also emerged in midbrain and prefrontal structures. Narrow-band γ oscillations have been recorded locally in attention-related midbrain networks, suggesting that midbrain visual structures can generate fast rhythms that support spatial selection[56]. In our model, SC was slightly more β-dominant but retained a stable γ component, consistent with reports of high-frequency selectivity in midbrain circuitry. In OFC and IL, γ power was strongest, with IL approaching a γ-dominant profile. This is broadly consistent with reports linking prefrontal γ activity to executive control and attention in mice[57]. Together, these spectral fingerprints formed a hierarchical gradient from mixed β/γ activity at early visual entry to stronger γ activity in prefrontal regions. This pattern is consistent with high-frequency–dominated inter-areal coordination in early pathways and greater involvement of prefrontal regions in integration and decision-related computations[58].

(2) Hierarchical delays: temporal organisation of information flow along the visual pathway

Using the same simulations, we computed each region's onset latency (time to first reach 10% of its peak) and near-peak latency (time to first reach 90% of its peak) relative to stimulus onset (Supplementary Fig. 11a,d). For first deviation, SC responded at 0 ms (defined as time zero for visual input), followed by LGN at 1 ms; LP at 7 ms, DS at 8 ms and TRN at 9 ms; OPN and OFC at 11 ms; PL at 17 ms; and VC, MC, IL and PPC mainly at 30–33 ms. For peak timing, SC peaked at 11 ms; LP and DS peaked at 28 ms and 30 ms, respectively; and many thalamic, cortical and prefrontal regions—including LGN, VC, TRN, MC, PPC, OFC and IL—peaked mainly at 50–54 ms. Several regions (e.g., LGN, TRN, OPN and OFC) showed an early onset deviation but a late peak, suggesting a later amplification or re-entrant process beyond the initial stimulus drive. Overall, the mean delay between neighbouring regions was 2.9 ms, suggesting that the network was not dominated by a small number of long-range edges that would produce an almost synchronous global response; instead, it preserved a recognizable hierarchical propagation gradient and staged recruitment.

More specifically, SC showed the earliest deviation from baseline and reached its peak rapidly, after which thalamic structures such as LGN and LP became active. The first significant response in VC occurred substantially later than in LGN or LP, suggesting that visual information need not propagate solely along a single serial LGN→VC→higher-cortex chain under this task setting. Instead, an SC-driven parallel route appeared to engage the broader network early. This trend is consistent with the view that SC can encode salient, low-spatial-frequency cues with shorter latencies, potentially enabling screening earlier than V1. It is also consistent with the role of LP (pulvinar) as a relay that rapidly forwards SC signals to higher circuits[59].

DS also exhibited a task-related LFP shift at 8 ms, consistent with an early subcortical bypass in which activity can reach striatal circuits before the canonical cortical pathway fully unfolds. In prefrontal-related regions (PL, IL and OFC), the first deviation was not uniformly the latest (for example, OFC could deviate relatively early), but peak responses were largely concentrated after 50 ms. This timing suggests later integration of sensory–motor information and stabilization of selection-related network states. This is broadly consistent with the view that PFC interacts with visual cortex–parietal circuits via β/γ rhythms during visual tasks, acting as a hub for top-down modulation and behavioral selection control[60].

Overall, during the visual task, the generated connectome exhibited staged propagation, with early subcortical fast drive, parallel multi-pathway engagement, and late convergence of peak responses across regions. Early nodes such as SC, LGN, LP, DS and TRN formed a main entry and branching set, whereas a later window showed broad peak co-occurrence across VC, PPC, MC and prefrontal regions. Together with the auditory-task analysis, these results suggest that the same generative-connectome scaffold can support modality-specific spectral and temporal patterns under shared structural constraints.

(3) Cross-regional causality and phase coupling

We next quantified dynamic coordination during the three-class video classification task using Granger causality and phase coupling computed from inter-areal LFPs. We constructed a directed information-flow network using Granger causality and visualized chord diagrams retaining the strongest 50% of connections to highlight core pathways (Supplementary Fig. 11e,f). Overall, the network shows a sensory–parietal–motor and decision hierarchy, with VC as a key hub. Among all edges, VC→PPC showed the strongest directed influence (Granger = 2.29). VC also showed strong directed influences on OPN, MC and TRN, consistent with distribution of sensory information toward higher-order integration and action-preparation circuits. This organization is compatible with bottom-up transfer of sensory evidence to parietal cortex, followed by coupling to action-related circuits. It is also consistent with reports that feedforward and feedback flows in the visual hierarchy can show frequency-specific directionality, with distinct oscillatory channels mediating feedforward–feedback communication[61].

In the subcortical-to-cortical direction, both SC→VC and LGN→VC showed strong causal influences (1.72 and 1.68, respectively) and remained among the strongest edges in the chord-diagram subnetwork. This is consistent with an SC-linked fast pathway contributing to cortical dynamics, in addition to the canonical LGN→V1/VC route, under the present task setting. This pattern is broadly consistent with reports that SC contributes to visual detection and orienting through multiple projections and can influence cortical processing[62]. Prefrontal-related regions also exhibited top-down influences: OFC→PL was among the strongest edges (Granger = 2.20), and OFC→VC was also substantial (1.49), consistent with modulation from value/executive signals through medial prefrontal circuits toward sensory processing. This directionality is broadly consistent with accounts in which orbitofrontal cortex conveys choice-related information to striatal and decision circuits and can influence behavioral output[63].

Phase-coupling analysis further quantified oscillatory coordination along these directional pathways. The PLV matrix (Supplementary Fig. 11g) showed two particularly strong synchrony modules. The first was tight phase locking among LGN, SC, OPN and TRN (e.g., LGN–OPN = 0.968, LGN–SC = 0.967, LGN–TRN = 0.959, SC–TRN = 0.881, OPN–TRN = 0.960), consistent with precise temporal alignment within early visual and thalamic gating circuits during the task. The second was high synchrony among VC, PPC and MC (extending to IL; e.g., MC–PPC = 0.987, MC–VC = 0.972, PPC–VC = 0.942, IL–VC = 0.949), consistent with coordinated routing of sensory evidence toward action-preparation networks. This is broadly consistent with frameworks in which phase coherence enables selective inter-areal information transfer[64] and with reports that task or attentional states can induce long-range high-frequency coupling between prefrontal and visual cortices[65].

Across Granger causality, PLV and MI, we observed consistent pathway-level patterns: directed drive from SC and LGN to VC co-occurred with strong synchrony within the early visual module, and the VC→PPC→MC chain was prominent across causal strength, phase locking and mutual information. Together, this cross-metric coherence suggests that, during task execution, the model exhibited coordination motifs that are qualitatively consistent with experimental reports on biological visual systems, including subcortical fast drive to cortex, selective integration, and top-down influences from cortical–parietal and prefrontal circuits.

(4) Population trajectories and low-dimensional geometry

The analyses above indicate that NIGC–ESN supports task-related spectral fingerprints, hierarchical delays and inter-areal directed interactions under visual drive. We next asked whether population-level representation geometry at single-neuron resolution shows comparable structure, by analysing low-dimensional spatiotemporal trajectories of regional population activity.

Using the same strategy as in the auditory task, we used a three-dimensional UMAP embedding to construct regional spatiotemporal trajectories (Supplementary Fig. 12a), enabling direct cross-task comparison of trajectory geometry. In this visual task, UMAP trajectories across the 12 regions typically showed a pronounced progression from an initial state along a continuous path toward a terminal state, often with local wandering near the endpoint. This pattern suggests stage transitions in task-driven population states and the emergence of stable, attractor-like neighbourhoods near decision or output phases. This behavior is consistent with hierarchical processing frameworks in which activity progresses from stimulus drive to evidence integration and then to selection, traversing distinct state regions and forming stable low-dimensional structure late in the trial[66].

Trajectory geometry differed systematically across regions. Sensory and feedforward-related regions (VC, SC, LGN and OPN) showed more monotonic trajectories with stable directions, resembling near-one-dimensional or weakly curved progressions, consistent with dynamics dominated by stimulus drive and feedforward transmission. Although visual cortex responses can contain rich high-dimensional detail, stimulus encoding can unfold along constrained low-dimensional axes that remain trackable in embedding space[67]. By contrast, higher-order integration, decision and execution-related regions (PPC, PL, IL, OFC, MC and DS) exhibited more curved and complex trajectories (multiple inflection points, local detours and partial reversals), consistent with recurrent integration, contextual modulation and contributions from action-related variables. Similar nonlinear decision geometry has been characterized in perceptual decision-making, where decision-related areas evolve along curved manifolds and yield geometry-dependent readout patterns across task contexts[68]. In addition, thalamic regions such as TRN and LP traversed larger-scale, arc-like trajectories and differed from the more monotonic progression of cortical sensory areas, consistent with roles in gain control, selective routing and state gating in thalamo–cortical loops. In particular, the prominent arc-like trajectory of TRN is consistent with a gating role that controls when specific information channels are engaged, rather than simply mirroring sensory representations.

The time-resolved heatmap of cross-regional tangent angles (Supplementary Fig. 12b) showed clear temporal modulation: some epochs exhibited broadly smaller angles (stronger alignment), whereas others exhibited larger angles (greater separation). This pattern is consistent with stage-dependent coordination: during some stages, multiple regions may be jointly driven and show transient alignment, whereas at other times region-specific computations dominate and directions diverge. This interpretation is broadly consistent with the “communication subspace” framework, in which cortical areas exchange information selectively through low-dimensional subspaces rather than coupling across all dimensions[69].

The time-resolved heatmap of cross-regional trajectory orthogonality (Supplementary Fig. 12c) further showed that many region pairs maintained medium-to-high orthogonality across epochs, consistent with population-state evolution unfolding in approximately orthogonal low-dimensional subspaces across regions. Such geometric organization is compatible with multiplexed processing, with sensory coding, evidence accumulation and action-related variables evolving in partially separated low-dimensional subspaces. This separation can, in principle, reduce interference between concurrent computations and aid downstream linear readout. Related mechanisms have been reported in population dynamics of cognitive tasks; for example, prefrontal circuits can map task variables into different dynamical subspaces across contexts, enabling flexible computations within the same network[70]. Likewise, mixed selectivity and subspace separation are thought to be important for linear separability and robustness in complex tasks[71].

### S5. Supplementary Methods

#### S5.1. Data sources and preprocessing

We used four datasets for structural calibration, connectome construction, and task-based functional validation. Neuronal-scale connectivity for structural calibration was taken from the MICrONS consortium minnie65_public dataset (v1300), which provides electron-microscopy reconstructions and synapse-level connectivity for mouse V1[72]. Whole-brain neuronal coordinates were taken from the Blue Brain Cell Atlas (second release, 2023), which includes refined annotations for inhibitory neurons[46]. For the auditory task, we used the Spoken Arabic Digit dataset (UCI Machine Learning Repository)[14], comprising 8800 utterances from 88 native speakers (44 male and 44 female; 10 digits repeated 10 times per speaker), sampled at 11,025 Hz (16-bit). We segmented the recordings into Hamming-windowed frames and extracted 13-dimensional MFCC features. For the visual task, we used an in-house video dataset (smartphone-recorded; 1080 × 1920 pixels, 30 fps) covering three representative action scenarios.

Because the raw connectome tables exceeded single-GPU memory, we streamed the CSV in contiguous blocks (skiprows with fixed nrows; 30,000–100,000 rows per chunk), rather than randomly sampling records. Contiguous reads preserve record adjacency and tend to retain spatial contiguity in the sampled population.

We normalised spatial coordinates by the diagonal length of the sampled volume's bounding box. We extracted 3D coordinates for pre- and postsynaptic sites and used their extrema to define the sampled volume bounds. We defined the maximum possible distance as the bounding-box diagonal:

$$D_{\max} = \sqrt{(x_{\max} - x_{\min})^2 + (y_{\max} - y_{\min})^2 + (z_{\max} - z_{\min})^2} \quad \text{(S6)}$$

We divided all coordinates by $D_{\max}$ to map positions to the unit interval [0, 1]. To define neuronal centroids, we averaged the 3D coordinates of all synaptic sites associated with each neuron ID. We computed the distance matrix from the normalised centroids and used the same neuron set to assemble the connectivity matrix. Connection probability was defined, for each distance bin, as the ratio of observed synaptic connections to the total number of possible neuron pairs. During model fitting, to avoid numerical instability in log-space regression, we excluded bins with zero or undefined connection probability and estimated parameters on the remaining bins.

#### S5.2. NIGC algorithmic details

We implemented NIGC and the baseline models in Python (CuPy) to enable computations at N≈30,000. We initialised $P$ from a power-law distance prior, $P_{ij}^{\text{base}} \propto d_{ij}^{-\alpha}$. We then applied node propensity modulation[73] using a rank-based weight vector $w$, updating $P_{ij} \leftarrow P_{ij}^{\text{base}} w_i w_j$. We monitored the skewness of the row-sum distribution of $P$, and when skewness exceeded $\gamma_{th}$=2.0 we down-weighted probabilities associated with the top 25% most connected nodes by a multiplicative factor $\eta$=0.5, after which $P$ was clipped to [0, 1].

We instantiated the adjacency matrix in two steps. First, we sampled candidate edges by retaining ($i$, $j$) when $R_{ij}<\lambda P_{ij}$ for $R_{ij}\sim U(0, 1)$, where $\lambda$ is a sparsity scaling factor used to match the target connection density. Second, we pruned candidate edges under a local wiring-length budget $E_{max}$ by keeping the nearest edges for each neuron until the cumulative wiring length reached the threshold[8]. We set $E_{max}$ proportional to the target mean degree via $E_{max}=d_{global}k_{target}$. We selected $\alpha$ by maximising a structural-entropy score $H=-\Sigma p_{ij}\log p_{ij}$ defined on a mixed distribution of first- and second-order neighbours[1]. We scanned $\alpha$ over 0.0–2.0 and chose the value that maximised $H$.

The NIGC procedure is summarised in Algorithm 1.

**Algorithm 1**: Neuro-Informed Generative Connectome (NIGC) generation

**Input:** Neuron coordinates $X\in R^{N\times 3}$, Target average degree $k$
**Output:** Adjacency matrix $A\in\{0, 1\}^{N\times N}$

Compute pairwise Euclidean distance matrix $D_{ij}=||x_i-x_j||_2$
Initialize geometric probability $P=D^{-\alpha}$
**Propensity Modulation:**
Generate weights $w$ such that $w_i\propto i^{-0.8}$
Update $P\leftarrow P\odot(ww^T)$
**Adaptive Balancing:**
Compute degree skewness $\gamma_1$ of $P$
If $\gamma_1$>threshold, apply penalty to high-degree hubs in $P$
**Sampling & Pruning:**
Sample candidate edges $A_{cand}\sim$Bernoulli($P$)
Calculate energy threshold $W$=mean($D$)×$k$
For each node $i$ in parallel:
Sort edges ($i$, $j$) by distance $D_{ij}$ ascending
Compute cumulative wire length $L_{cum}$
Keep edge ($i$, $j$) if $L_{cum}\leq W$, else remove
Return $A$

For comparison, we implemented three baseline generative models.

(1) Economical wiring model[8]. We defined an edge score based on a cost–benefit trade-off:

$$S_{ij}=\beta\cdot \mathrm{O}(N_i,N_j)-\lambda\cdot(d_{ij}/\bar{d})^{\gamma} \tag{S7}$$

where $O(N_i, N_j)$ denotes the overlap ratio of the k-nearest-neighbour (kNN) sets, and $\lambda$ and $\gamma$ control the strength of distance penalisation. The algorithm computed scores within each node's kNN candidate set and selected the global top $M$ edges.

(2) Homophily model[9]. We defined a score combining spatial distance and feature similarity:

$$S_{ij}=d_{ij}^{-\alpha}\cdot\exp(-\eta\cdot|z_i-z_j|/\sigma_z) \tag{S8}$$

where z is the neuron coordinate along the cortical-depth axis (or another phenotypic feature), and $\eta$ controls homophily strength.

(3) Fully random model. Following the Erdős–Rényi random-graph assumption, we fixed node and edge counts, and sampled edges uniformly at random from all node pairs. All baseline models were matched to NIGC in target connection density (mean degree) to enable fair comparisons.

S5.3. ESN implementation details

We constructed an echo state network (ESN) of leaky-integrator units[74], with reservoir topology specified by the NIGC-generated anatomical connectome. The implementation details of the network dynamics and training procedure were as follows.

The reservoir comprised N=12966 neurons, corresponding to the NIGC-generated mouse auditory-pathway connectome. The internal reservoir weight matrix $W_{res} \in R^{N\times N}$ was set to the generated weighted adjacency matrix[75]. To enforce the echo-state property and operate in a high-capacity regime, we performed spectral scaling on $W_{res}$. Specifically, we computed the mean eigenvalue modulus of the unscaled matrix, denoted $\overline{|\lambda|}_{raw}$, and rescaled $W_{res}$ as[76]:

$$W_{res} \leftarrow W_{res} \cdot \frac{\rho_{target}}{\overline{|\lambda|}_{raw}} \tag{S9}$$

After grid-search optimization, the target mean eigenvalue modulus was set to $\rho_{target}$=0.8.

The input-node set $S_{in}$ contained $N_{in}$=216 neurons, randomly selected from the cochlear nucleus (CN). The readout-node set $S_{out}$ contained $N_{out}$=1024 neurons, randomly selected from prefrontal cortex (PFC)-related regions (PL, IL, and OFC). The input weight matrix $W_{in} \in R^{N\times Nin}$ was nonzero only on rows corresponding to $S_{in}$, and its nonzero elements were drawn from a uniform distribution $U(\mu-\delta, \mu+\delta)$ with $\mu$=0.1 and $\delta$=0.05.

For each 13-dimensional mel-frequency cepstral coefficient (MFCC) vector $u(t) \in R^{13}$, we assigned its components to the 216 input nodes in a cyclic manner: input node $i$ received component $u(i \bmod 13)(t)$. The reservoir state $x(t) \in R^{N}$ evolved in discrete time according to the leaky-integrator update rule:

$$x(t+1) = (1-\alpha)x(t) + l \cdot \tanh(W_{in}u(t+1) + W_{res}x(t)) \tag{S10}$$

where $l$ is the leak rate controlling the effective time constant of neuronal states. Based on parameter sweeps, we set $l$=0.1. The nonlinearity was the hyperbolic tangent, tanh(·).

Task training followed the standard ESN paradigm: the reservoir weights $W_{res}$ and input weights $W_{in}$ were fixed, and only the readout weights were optimised. For each sample sequence, we used the subvector of reservoir states on $S_{out}$ at the final time $T$, $x_{out}(T) \in R^{1024}$, as the feature representation. The classifier was multinomial logistic regression, optimised with L-BFGS. The objective was the cross-entropy loss with $L_2$ regularization, and the maximum number of iterations was 200. Training was implemented on a CPU (Intel Core i9) and GPU (NVIDIA RTX 4080) using scikit-learn, with CuPy for acceleration. All reported accuracies were evaluated on an independent test set.

S5.4. Definition of functional metrics

Our functional analyses used time series of reservoir neuronal states. We first aggregated single-neuron activity within each brain region into a regional series and then defined spectral fingerprints, hierarchical delays, and inter-regional dependence metrics on these signals. Let region $r$ contain neuronal indices $V_r$, and denote the reservoir state of neuron $i \in V_r$ by $x_i(t)$. The region-level LFP was defined as the time-point mean of neuronal states within the region[24]:

$$\mathrm{LFP}_r(t) = \frac{1}{|V_r|} \sum_{i \in V_r} x_i(t) \tag{S11}$$

When computing delay-related temporal statistics, we used $|\mathrm{LFP}_r(t)|$ to avoid polarity-dependent effects on threshold-crossing detection, and used the peak amplitude within the analysis window $\Omega$ for normalization.

Spectral fingerprints. Spectral fingerprints were represented by the relative power of classical frequency bands of the regional LFP. We band-pass filtered $\mathrm{LFP}_r(t)$ into five bands—$\delta$:[0.5, 4] Hz, $\theta$:[4, 8] Hz, $\alpha$:[8, 13] Hz, $\beta$:[13, 30] Hz, and $\gamma$:[30, 100] Hz—using a fourth-order Butterworth filter with forward–backward (zero-phase) implementation[77]. This yielded band-resolved signals $\mathrm{LFP}_{r,b}(t)$ ($b \in \{\delta, \theta, \alpha, \beta, \gamma\}$). Band power was defined as the mean squared amplitude over the analysis window $\Omega$:

$$E_{r,b} = \frac{1}{|\Omega|} \sum_{t \in \Omega} \mathrm{LFP}_{r,b}(t)^2 \tag{S12}$$

Relative power was then defined as:

$$P_{r,b} = \frac{E_{r,b}}{\sum_{b'} E_{r,b'}} \tag{S13}$$

The spectral fingerprint vector of region $r$ was $\mathbf{P}_r=[P_{r,\delta}, P_{r,\theta}, P_{r,\alpha}, P_{r,\beta}, P_{r,\gamma}]$, and was used to compare band composition across regions or across model conditions.

Hierarchical delays. Hierarchical delays were quantified using threshold-crossing times of the regional LFP. For each region $r$, the peak amplitude within $\Omega$ was $A_r=\max_{t\in\Omega}|\mathrm{LFP}_r(t)|$. The onset latency was the first time point at which $|\mathrm{LFP}_r(t)|\geq 0.1A_r$. The rise-to-peak latency was defined analogously using $|\mathrm{LFP}_r(t)|\geq 0.9A_r$. The 10% and 90% thresholds help reduce sensitivity to baseline offsets and post-peak decay in latency estimation[78]. In implementation, threshold crossings were detected on discrete samples and converted to milliseconds using the sampling rate.

To characterize intrinsic temporal properties without task drive, we generated spontaneous activity using low-amplitude noise input[23] and estimated within-region timescales from autocorrelation decay of the regional LFP. Specifically, we drove the ESN with independent Gaussian noise for

$T_{noise}$=2000 steps (noise standard deviation $\sigma$=0.02), obtained reservoir states, and aggregated them to a regional LFP $L_R(t)$. For each region, $L_R(t)$ was demeaned and the normalized autocorrelation $AC_R(l)$ was computed for lags $l \in [0, l_{max}]$ with $l_{max}$=400, normalized by $AC_R(0)$. The intrinsic timescale $\tau_R$ was estimated as the discrete integral over non-negative lags, with negative autocorrelation values truncated to zero:

$$\tau_R = \sum_{l=0}^{l_{max}} \max\left(0, AC_R\left(l\right)\right) \quad \text{(S14)}$$

Inter-areal directional dependence and phase coupling. Directional inter-regional dependence was quantified using Granger causality. For any pair of regions $i$ and $j$ with regional LFP series $LFP_i(t)$ and $LFP_j(t)$, we compared a restricted autoregressive model (lags of $LFP_j$ only) with a full model (lags of both $LFP_i$ and $LFP_j$). With maximum lag order $L$, the restricted and full models were[79]:

$$LFP_j(t) = \sum_{k=1}^{L} b_k LFP_j(t-k) + \varepsilon_R(t) \quad \text{(S15)}$$

$$LFP_j(t) = \sum_{k=1}^{L} a_k LFP_i(t-k) + \sum_{k=1}^{L} b_k LFP_j(t-k) + \varepsilon_F(t) \quad \text{(S16)}$$

If $Var(\varepsilon^{(F)}) << Var(\varepsilon^{(R)})$, we considered the predictive gain from $i \rightarrow j$ to be positive and defined the Granger causal strength as the log variance ratio:

$$GC_{i \rightarrow j} = \ln \frac{Var(\varepsilon_R)}{Var(\varepsilon_F)} \quad \text{(S17)}$$

In practice, we computed Granger-causal strengths across lags from 1 to $L$ and aggregated them into a single edge weight. We used $L$=10 by default and set negative estimates to 0 to retain interpretable directed gains.

Inter-regional phase coupling was quantified using the phase-locking value (PLV). For a given band $b$, we band-pass filtered the regional LFP and applied the Hilbert transform to obtain the analytic signal $z(t)=A(t)e^{j\Phi(t)}$. From the instantaneous phases $\Phi_i(t)$ and $\Phi_j(t)$, we computed the phase difference $\Delta\Phi(t)=\Phi_i(t)-\Phi_j(t)$. PLV was defined as the magnitude of the time-averaged unit phase-difference vector[80]:

$$\mathrm{PLV}_{i,j} = \left| \frac{1}{|\Omega|} \sum_{t \in \Omega} e^{j\Delta\phi(t)} \right| \in [0,1] \quad \text{(S18)}$$

Low-dimensional trajectories and embedding quality. Low-dimensional spatiotemporal trajectory analysis was based on manifold embedding of within-region neuronal population activity. For each region , we assembled neuronal states within the analysis window into a matrix $X_r \in R^{T \times |V_r|}$ (rows: time points; columns: neurons) and applied UMAP to obtain $Y_r \in R^{T \times d}$. This defines the time-evolving trajectory $y_r(t) \in R^d$. Unless otherwise specified, we used d=3. To reduce the impact of coordinate-scale differences on geometric statistics, we applied per-dimension min–max normalization to the embedded coordinates before computing trajectory angles and orthogonality.

Embedding quality was evaluated using neighbourhood preservation metrics. With sample size $n=T$, we defined the k-nearest-neighbour sets in the original space and embedding space as $N_k^H(i)$ and $N_k^L(i)$, respectively. Trustworthiness was computed as[29]:

$$T(k)=1-\frac{2}{nk(2n-3k-1)}\sum_{i=1}^{n}\sum_{j\in N_k^L(i)\setminus N_k^H(i)}\left(r_H(i,j)-k\right) \tag{S19}$$

where $r_H(i, j)$ is the rank of sample $j$ among the neighbours of sample $i$ in the original space. Continuity was defined as the dual metric[29]:

$$C(k)=1-\frac{2}{nk(2n-3k-1)}\sum_{i=1}^{n}\sum_{j\in N_k^H(i)\setminus N_k^L(i)}\left(r_L(i,j)-k\right) \tag{S20}$$

where $r_L(i, j)$ is the neighbour rank in the embedding space. The kNN retention rate was defined as the average fraction of overlap between the two kNN sets[30]:

$$P(k)=\frac{1}{n}\sum_{i=1}^{n}\frac{\left|N_k^H(i)\cap N_k^L(i)\right|}{k} \tag{S21}$$

These three metrics were implemented consistently with those used for dimensionality selection in Section S3.1, and the same $k$ settings were used across target dimensions and conditions for fair comparison.

Trajectory geometric relations were quantified by tangent-direction angles and orthogonality. For any two regions $r$ and $s$, we approximated the local tangent direction at time $t$ using a centred difference with time-window half-width $w$, defining the direction vector $d_r(t)=y_r(t+w)-y_r(t-w)$ (and analogously $d_s(t)$). The cosine of the inter-trajectory angle was:

$$\cos\theta_{r,s}(t)=\frac{d_r(t)\cdot d_s(t)}{\left\|d_r(t)\right\|\left\|d_s(t)\right\|} \tag{S22}$$

Orthogonality was defined as $O_{r,s}(t)=1-|\cos\theta_{r,s}(t)|$, which attains its maximum when the two directions approach 90°.

Structure–function coupling via projection patterns and induced causal responses. We evaluated coupling between structural projection patterns and stimulus-evoked causal responses to quantify structure–function correspondence in NIGC-generated connectomes. We first constructed a region-to-region weighted adjacency matrix $A\in R^{R\times R}$ from the neuron-level reservoir weight matrix $W_{res}$. Specifically, $A_{ij}$ was defined as the sum of absolute connection weights from source region $i$ to target region $j$, and was normalized by the source/target neuron counts divided by $N_iN_j$, yielding the average strength per neuron pair. On the region graph, we computed a k-hop structural projection field from each source region using a row-normalized random-walk form[81]: letting $P$ be the row-normalized version of $A$, and letting the initial vector $v$ be one-hot at the source region (1 at the source and 0 elsewhere), the structural projection vector was $s=\Sigma\lambda^{h-1}vP^h$, with decay factor $\lambda=0.05$ and $k=2$ in experiments. To avoid self-projection effects, the source component (diagonal term) was set to zero in subsequent comparisons.

The corresponding causal functional strength was obtained by applying regional pulse stimulation and measuring whole-network responses[82]. For each source region, we constructed an external input sequence of length $T=400$, and applied a square-wave pulse (amplitude 0.5) at $t=50$ for two time steps. We simulated reservoir states and re-aggregated them into regional LFPs. For each target region, we used the mean of the pre-stimulation window as the baseline, defined the response segment as the deviation of LFP from baseline during the stimulation window, and quantified response strength by the absolute integral $\Sigma_t|L_R(t)-L_R^{base}|$, yielding a source-to-target dynamical response vector $d$ (with the source self-term set to zero).

To compare distributions at the matrix level, we normalized the structural projection matrix and response matrix to [0, 1] by dividing each by its global maximum. Final structure–function coupling was evaluated by edge-wise correlations (Pearson $r$ and Spearman $\rho$)[83] after removing diagonal terms. We additionally computed Spearman correlations row-wise (by source) and column-wise (by target) to characterize coupling heterogeneity across source and target regions.

### S6. Supplementary Discussion

#### S6.1. Cross-scale geometric laws

At the neuronal scale, we find that, within the measurable range in mouse V1, connection probability decays with distance in a manner better captured by a power law than by exponential or Gaussian kernels commonly used for mesoscopic connectivity modelling. In contrast, when connections are aggregated to the level of brain regions or large-scale parcels, structural datasets across multiple species are more often well described by an exponential distance decay[2]. This contrast suggests scale dependence in the statistical form of the distance–connection probability relationship, such that microscopic and mesoscopic levels may not share the same effective kernel class.

This neuronal-level power-law versus mesoscopic-exponential contrast naturally suggests a scale-separation picture. At the microscopic level, networks remain sparse overall, yet the heavy tail of a power law assigns relatively higher probability to a small number of long-range edges, preserving sparse channels needed for long-range integration. At the coarser region scale, aggregation-induced averaging—together with anatomical boundaries and metabolic and wiring costs—drives the effective mean connectivity toward a faster, exponential-like decay. In other words, a microscopic heavy tail does not require the mesoscopic level to retain the same heavy-tailed form; instead, mesoscopic statistics may reflect an aggregate effective kernel shaped jointly by spatial averaging and resource constraints.

More broadly, this scale separation is unlikely to be specific to mouse V1. At the neuronal-scale connectome level, power-law distance dependence has also been reported (e.g., in Drosophila)[1], whereas exponential distance decay is more common in multi-species region- or parcel-scale datasets[84]. Taken together, these observations point to a cross-species regularity: microscopic scales preserve sparse but important long-range channels via power-law tails, while mesoscopic scales exhibit faster exponential decay under anatomical and cost constraints. NIGC implements this perspective with a two-step design. It adopts a power-law distance prior at the neuronal level, while applying interpretable resource-based constraints to long-range edges via node-level connection-propensity modulation and an energy budget. This preserves the long-range integration potential of heavy tails while avoiding excessive and physiologically implausible long-range edges, providing a testable mechanistic bridge between neuronal-scale power-law scaling and mesoscopic exponential decay.

#### S6.2. Empirical sufficiency and non-uniqueness of the constraint set

Main-text results together with supplementary controls indicate that, given neuronal-scale geometric coordinates, the constraint combination used in NIGC supports key structural statistics and multidimensional functional phenotypes under the same fixed, unified dynamical regime, and that agreement decreases systematically under stepwise ablation. This provides a compact and testable empirical constraint set for connectome generation, enabling structure–function correspondence to be evaluated within a single evidence chain.

However, empirical sufficiency does not imply uniqueness or strict minimality. In principle, other equivalent or near-equivalent generative schemes may achieve comparable agreement across different collections of structure–function metrics. We do not attempt an exhaustive search of the broader model space. Instead, we start from a small set of physiologically interpretable constraints and test whether they can support a falsifiable bridge from microscopic connectivity to system-level function.

Accordingly, the value of constraint ablation is not only in comparing outcomes, but also in enabling a falsifiable mode of mechanistic localisation. When removing a constraint induces a functional degradation pattern that aligns with its structural role, this suggests that the functional phenomena are less likely to arise from incidental dynamical choices, task readouts or analysis conventions, and instead depend on structural priors within the unified dynamical substrate. In this way, the work provides actionable empirical constraints and a clear baseline for incremental extensions towards richer mechanisms (e.g., cell types, laminar structure[85], delay distributions[86]).

#### S6.3. Spontaneous emergence of function under unified dynamics

A key functional observation is that the multidimensional functional features analysed here were not obtained by directly fitting functional data; instead, they emerged under task drive in the spoken-digit classification task, given structural priors and a fixed, unified dynamical regime. This is central to the proposition of structural-prior sufficiency: it shifts the focus from how to train a model to match activity to whether structural priors can support consistent dynamical organisation without explicitly fitting functional data. Under this setting, structure–function consistency can be interpreted more directly in terms of the structural priors, rather than compensatory effects of high-dimensional learning or many tunable parameters.

From this standpoint, our functional controls and ablations emphasise qualitative patterns and cross-metric consistency over isolated significance tests for each metric. With relatively few constraints and strong correlations among functional metrics, repeatedly applying highly similar quantitative measures can double-count the same error sources, making it harder to identify where a model deviates from empirical observations. By contrast, treating spectral fingerprints, hierarchical delays, directional causality and phase coupling, low-dimensional population-trajectory geometry, and responses to pathological perturbations as complementary observational windows helps localise where systematic deviations first emerge when a structural prior is removed or disrupted, especially when NIGC, baselines and ablations are compared under identical dynamics.

This multi-window strategy supports our goal of identifying empirically critical structural constraints. Rather than drawing conclusions from a single metric, we aim to determine, via a consistent evidence chain across scales and phenotypes, which structural constraints are critical for biologically consistent functional organisation. Functional analysis therefore serves both as a check of basic task-related function and as a stress test of the interpretability and transferability of structural priors, providing a reference frame for extensions to more complex tasks or more realistic dynamics.

#### S6.4. Predictions under pathological perturbations

Pathological perturbation simulations illustrate how this framework can be used to probe disease-related functional reconfiguration. In these experiments, we neither retune parameters nor change generative rules or task settings. Instead, we impose lesion-like weakening by directed attenuation of edges incident to selected regions, and examine whether the resulting changes in functional metrics match the direction of effects reported in lesion studies and disease animal models. This design enables more transparent attribution of perturbation effects to structural changes, maintaining interpretability from structural priors through dynamical responses to observable functional metrics.

In hippocampal-related interventions, attenuating HPC connectivity decreases $\theta/\gamma$ power and increases $\delta$ power, consistent with spectral reorganisation trends reported across hippocampal damage contexts[87]. Obtaining directionally consistent responses without specialised fitting suggests that, within this framework, structural priors can support not only task-relevant dynamics under healthy conditions but also qualitatively consistent predictions under perturbation. By applying local perturbations at the neuronal-scale connectome level, we can predict system-level changes in functional metrics for comparison with existing or future experimental data.

Compared with comprehensive disease models, this minimal-intervention approach prioritises controllability and traceability. It emphasises directionality and mechanistic localisation rather than adding degrees of freedom to accommodate complex disease progression, while leaving room to introduce plasticity, compensation and more realistic dynamics.

#### S6.5. Limitations and future extensions

Although NIGC indicates structure–function consistency under the current data and settings, several limitations remain. First, the empirical benchmarks in the structure–function evidence chain are not all drawn from perfectly matched circuits and task paradigms. Structural calibration mainly relies on neuronal-scale connectome data from visual cortex, whereas functional validation spans both auditory and visual paradigms. Even within the auditory analyses, some evidence comes from broader auditory cognition and pathway studies rather than being restricted to the spoken-digit classification task. This design tests consistency and transferability across sensory channels and task conditions; however, it also requires a cautious delimitation of scope when circuits and tasks differ.

Second, at the dynamical level we use a continuous-valued ESN reservoir under a fixed, unified dynamical regime. While effective for testing how structural priors support task execution, it does not include spiking, short-term synaptic plasticity[88], long-term learning[89] or neuromodulation[90]. This limits its ability to address behaviours that require long-term learning, rule switching[91] or strategy exploration[92]. Third, the current constraint set does not explicitly encode cell types, laminar organisation, E/I ratios, conduction-delay distributions, or local extracellular space and matrix[93] (and other microenvironmental factors). We also observe that certain structural or functional statistics (e.g., the extreme tail of the degree distribution, precise amplitudes in specific bands, or behavioural extremes under certain paradigms) may be more sensitive to such microscopic details. This choice helps attribute the observed functional signatures to the structural prior, rather than to cellular-level tuning or synaptic plasticity.

Future work can proceed along several directions. (i) Keep the current geometric and energy assumptions fixed, and incorporate cell-type and laminar annotations[85] to construct type-specific connection probabilities and energy budgets; test whether these additions explain remaining deviations in higher-order statistics and functional phenomena. (ii) Integrate more realistic axon–dendrite morphology and conduction-delay distributions into the generative rules[94] to support finer time-scale characterisation of phase coupling and cross-frequency interactions, and to link these dynamics to slower physiological regulatory processes. (iii) Leverage increasingly rich multi-species 3D coordinate datasets to extend NIGC to primate and ultimately human cortex, assessing whether geometry and node propensity modulation remain effective priors under different brain volumes and energy budgets. (iv) Integrate NIGC as a structural prior into neural-activity foundation models or large-scale digital-brain platforms[95]. For example, use NIGC to generate neuronal-scale connectivity within digital-brain frameworks, or to constrain representations of spatial structure and sparse wiring in data-driven models, thereby strengthening links among high-fidelity simulation, task-level behaviour and interpretable structure.

### S7. Supplementary Figures

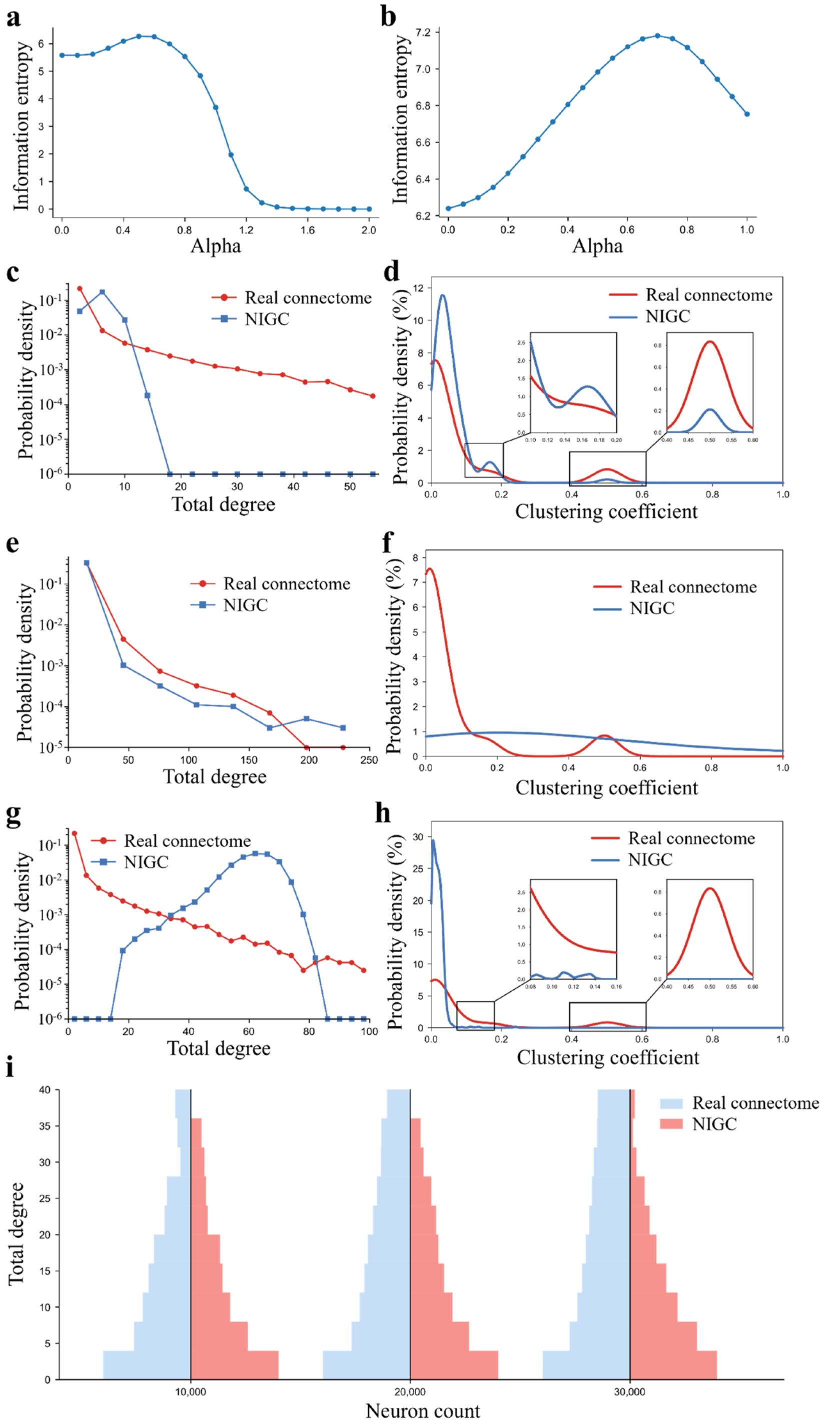

**Supplementary Fig. 1. Maximum-entropy choice of the power-law exponent and effects of constraint ablations on structural agreement. a,** To select the exponent $\alpha$ of the geometric power-law prior, we swept $\alpha$ and computed the structural information entropy of the generated connectome[1], yielding the entropy–$\alpha$ curve. **b,** For the multi-region auditory-pathway connectome used in the functional analyses, we repeated the sweep and computed structural information entropy, yielding the entropy–$\alpha$ curve. **c–h,** Structural-constraint ablations. While matching the node set and mean degree, we removed the node-level connection-propensity modulation (**c,d**), the global energy-budget constraint (**e,f**) or the information-entropy maximisation constraint (**g,h**), regenerated connectomes, and compared them with the empirical V1 microcircuit connectome using two structural statistics: the degree distribution (**c,e,g;** y axis, log-probability density) and the clustering-coefficient distribution (**d,f,h;** insets, enlargements of the low- and high-clustering regimes). Red curves denote the empirical connectome and blue curves denote the generated connectomes under the corresponding ablation conditions. **i,** Sensitivity to sampling scale. For different sample sizes (10k, 20k and 30k neurons), we extracted subsamples from the empirical connectome and the NIGC-generated connectome and compared their degree histograms to assess finite-sampling consistency and bias trends (red, empirical; blue, generated).

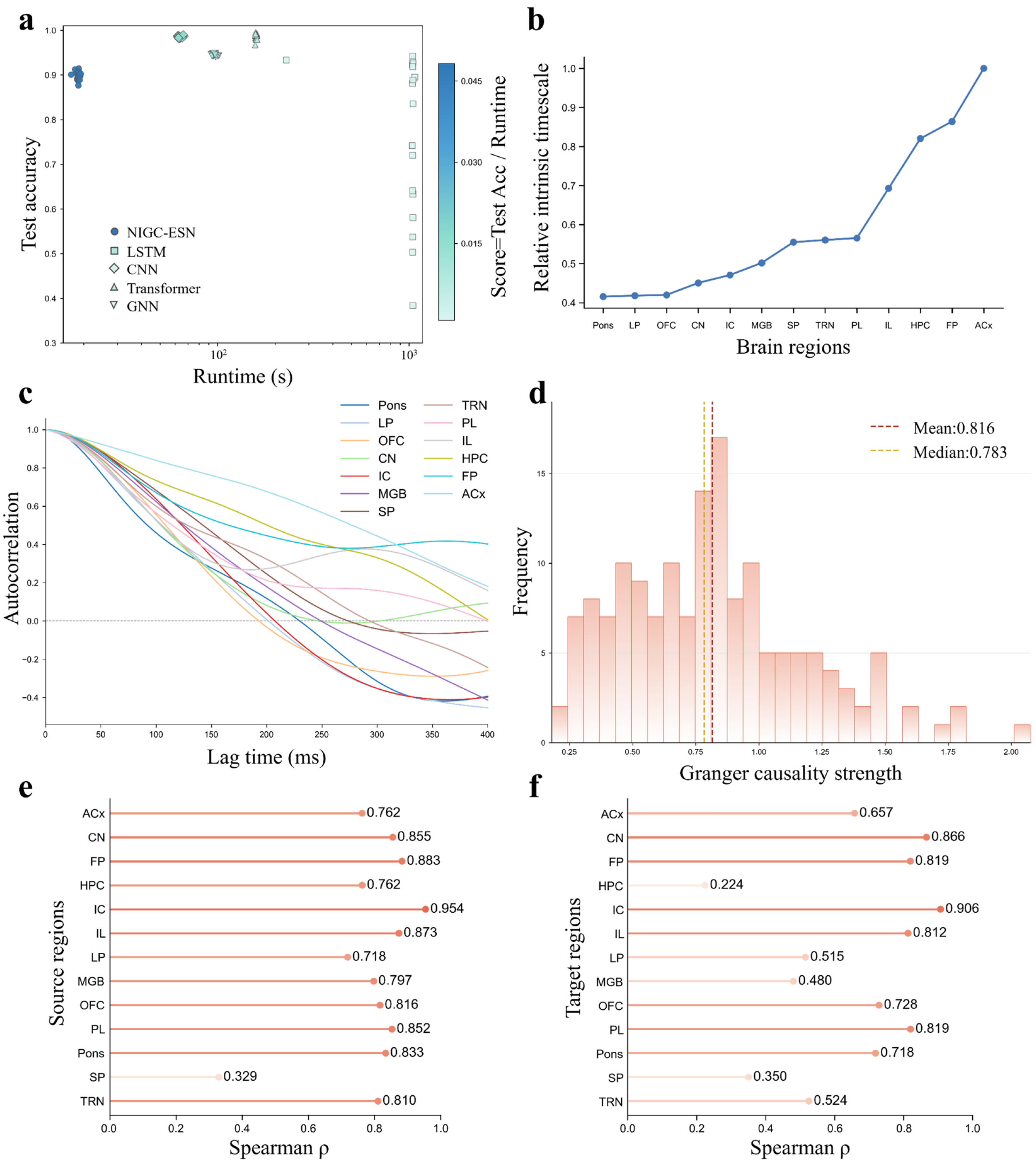

**Supplementary Fig. 2. Additional quantification of the performance–efficiency trade-off, intrinsic timescales and structure–function coupling in the spoken-digit task. a,** Under a matched total parameter budget (fixed reservoir parameters plus trainable readout parameters), we compared the efficiency–performance trade-off between NIGC–ESN and end-to-end baselines (LSTM, CNN, Transformer and GNN). The x axis shows runtime and the y axis shows test accuracy; colour encodes a composite score (accuracy divided by runtime). Each point denotes an independent repeat. **b–c,** Regional statistics of intrinsic timescales and autocorrelation. **b,** Hierarchical gradient of relative intrinsic timescales across regions (normalised to 0–1). **c,** Decay of the regional LFP autocorrelation function as a function of lag. **d,** Distribution of Granger-causality strengths aggregated across all directed region pairs; dashed lines indicate the mean and median. **e–f,** Hierarchy-stratified Spearman correlations for structure–function coupling. After pairing structural projection strength with pulse-evoked causal response strength for each region pair, we computed Spearman's ρ by fixing the source region and pooling across targets (**e**), or by fixing the target region and pooling across sources (**f**).

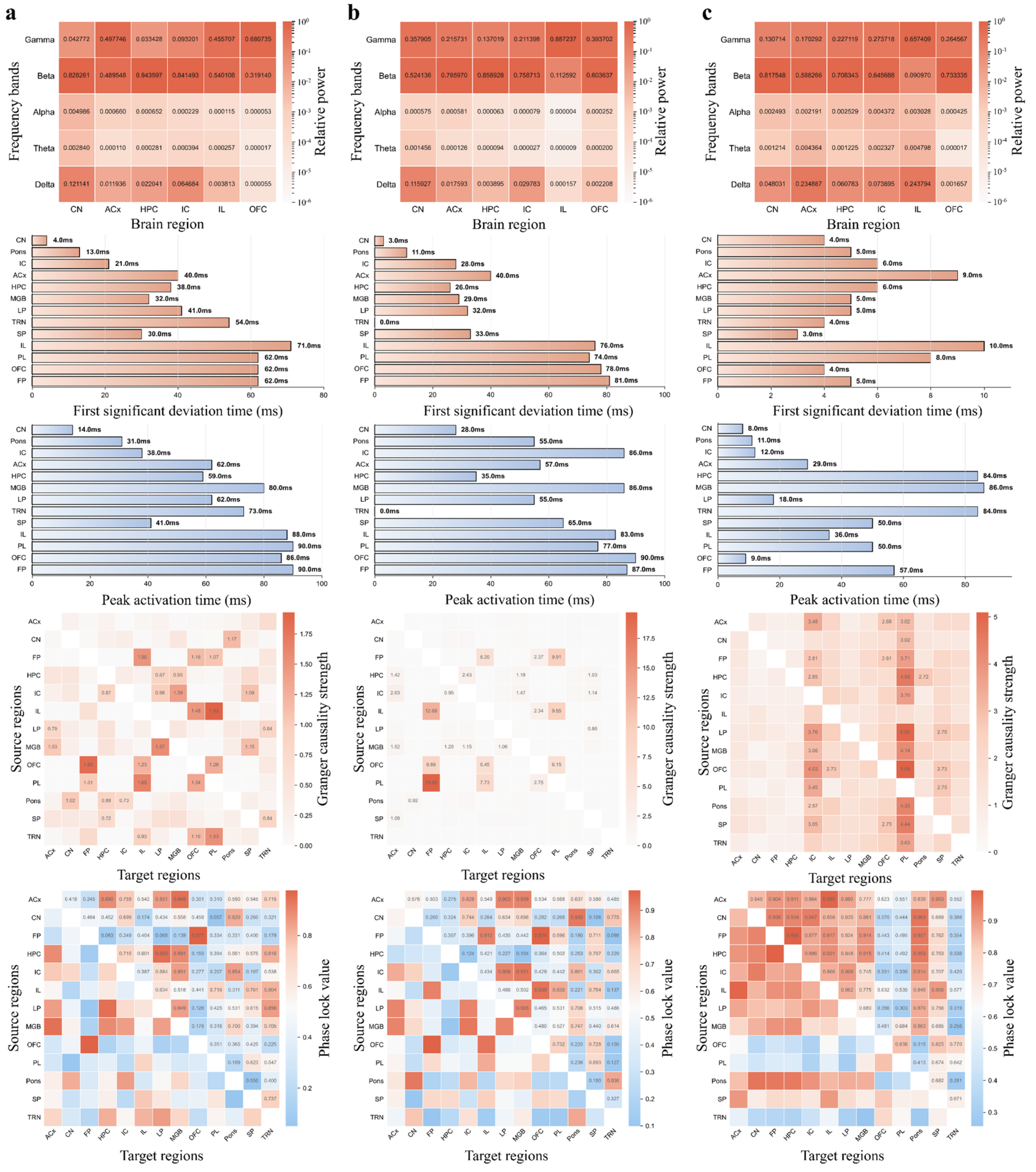

**Supplementary Fig. 3. Basic functional phenotypes for three structural baselines. a–c,** Results for the economical wiring, homophily and mean-degree-matched random baselines under the same spoken-digit task drive and analysis pipeline as in the main text. In each column (top to bottom): relative power across five frequency bands (δ/θ/α/β/γ) for six representative regions (CN, ACx, HPC, IC, IL and OFC); time for each region's LFP to reach 10% of its peak (onset latency); time to reach 90% of its peak (near-peak latency); cross-regional Granger-causality matrix; and cross-regional PLV phase-locking matrix (source × target).

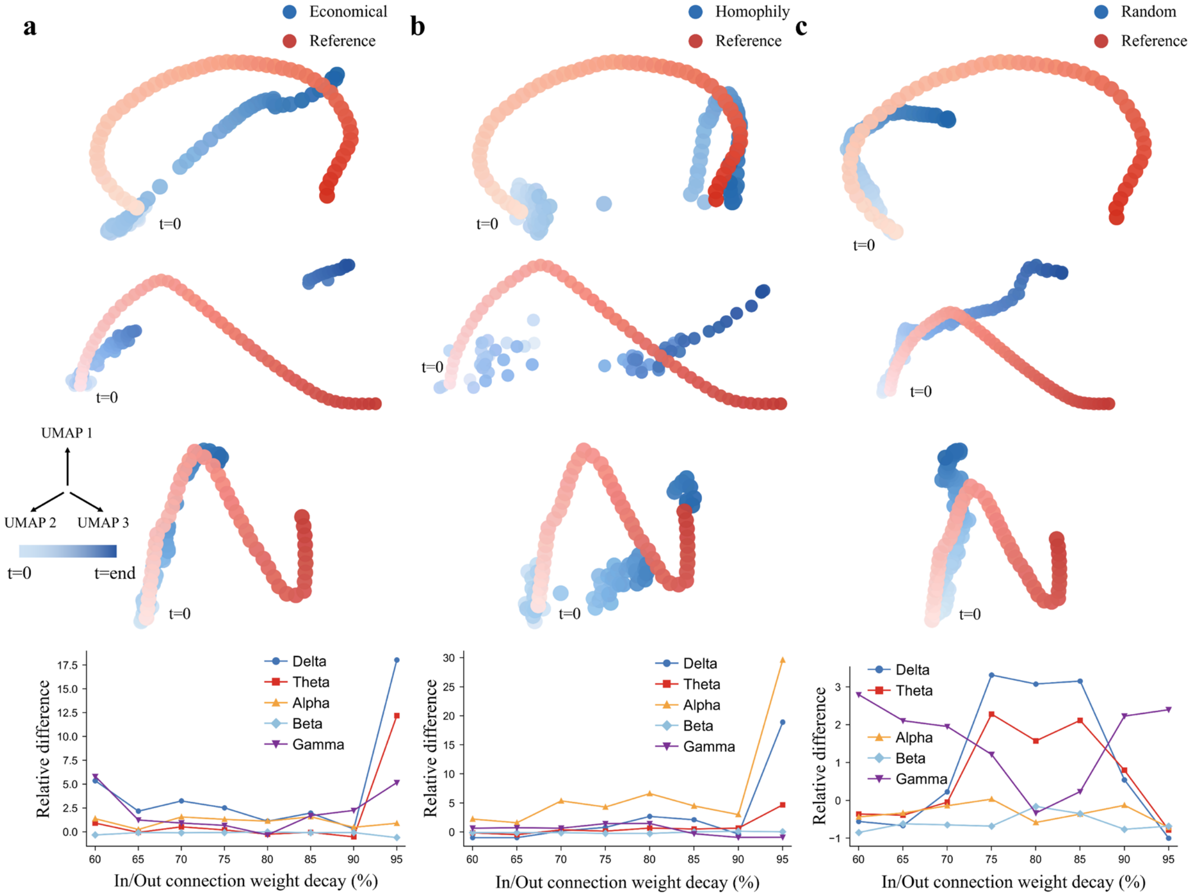

**Supplementary Fig. 4. Intermediate and high-level functional phenotypes for three structural baselines. a–c,** Results for the economical wiring, homophily and fully random baselines. In each column (top to bottom): low-dimensional spatiotemporal trajectories of ACx, PL and OFC compared with reference trajectories from previous studies (red, previous study; blue, baseline network; colour intensity indicates temporal progression; t=0, start); and, after graded attenuation of incoming/outgoing HPC connections, the dependence of HPC relative power across five LFP bands on attenuation strength.

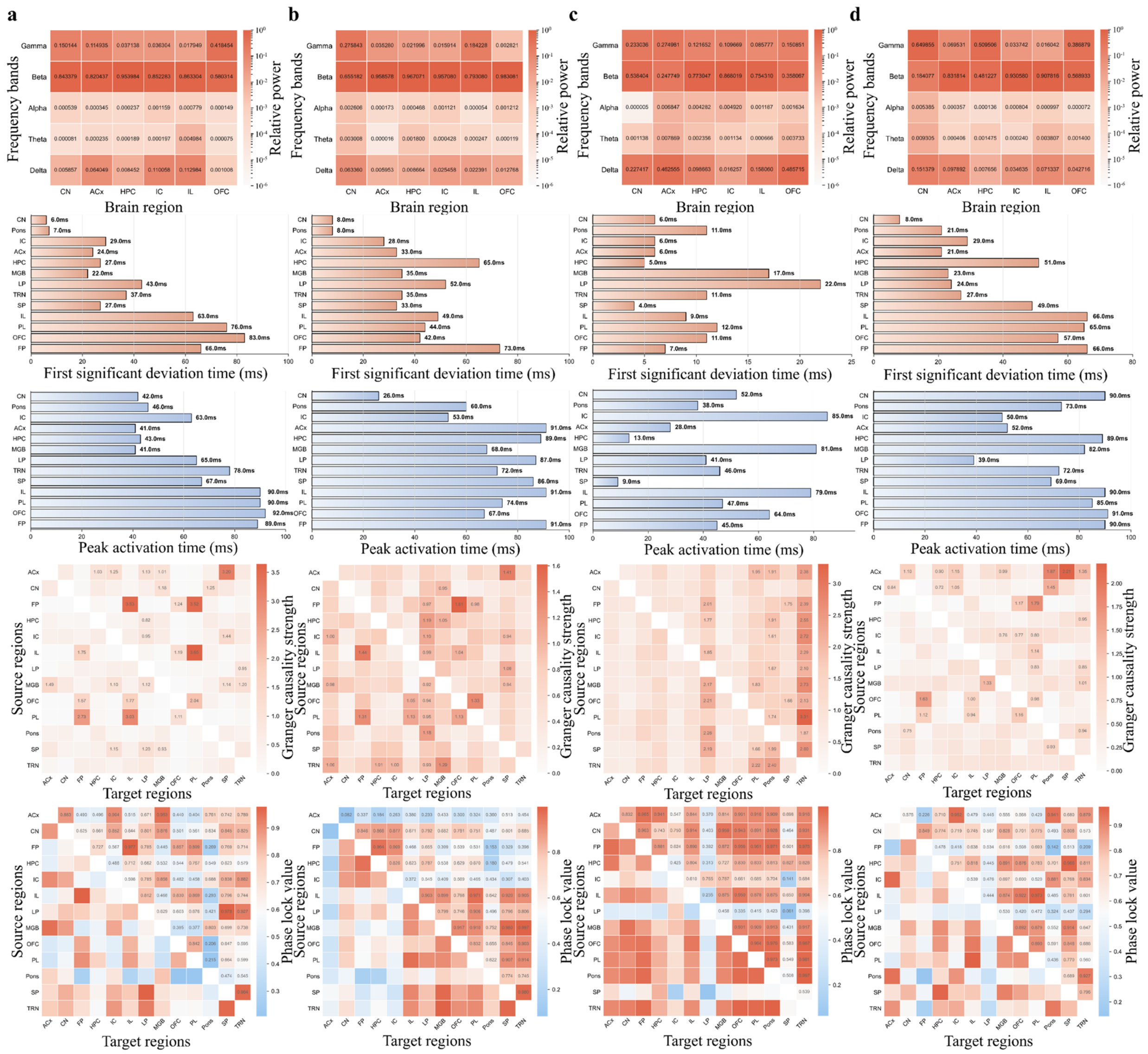

**Supplementary Fig. 5. Basic functional phenotypes under ablations of four structural constraints. a–d,** Results after removing, within the NIGC generation framework, the geometric power-law constraint (**a**), the node-level connection-propensity modulation constraint (**b**), the energy-budget constraint (**c**) and the information-entropy maximisation constraint (**d**) (all other procedures match the main text). In each column (top to bottom): relative power across five frequency bands for six representative regions (CN, ACx, HPC, IC, IL and OFC); time to reach 10% of peak; time to reach 90% of peak; cross-regional Granger-causality matrix; and cross-regional PLV phase-locking matrix (source × target).

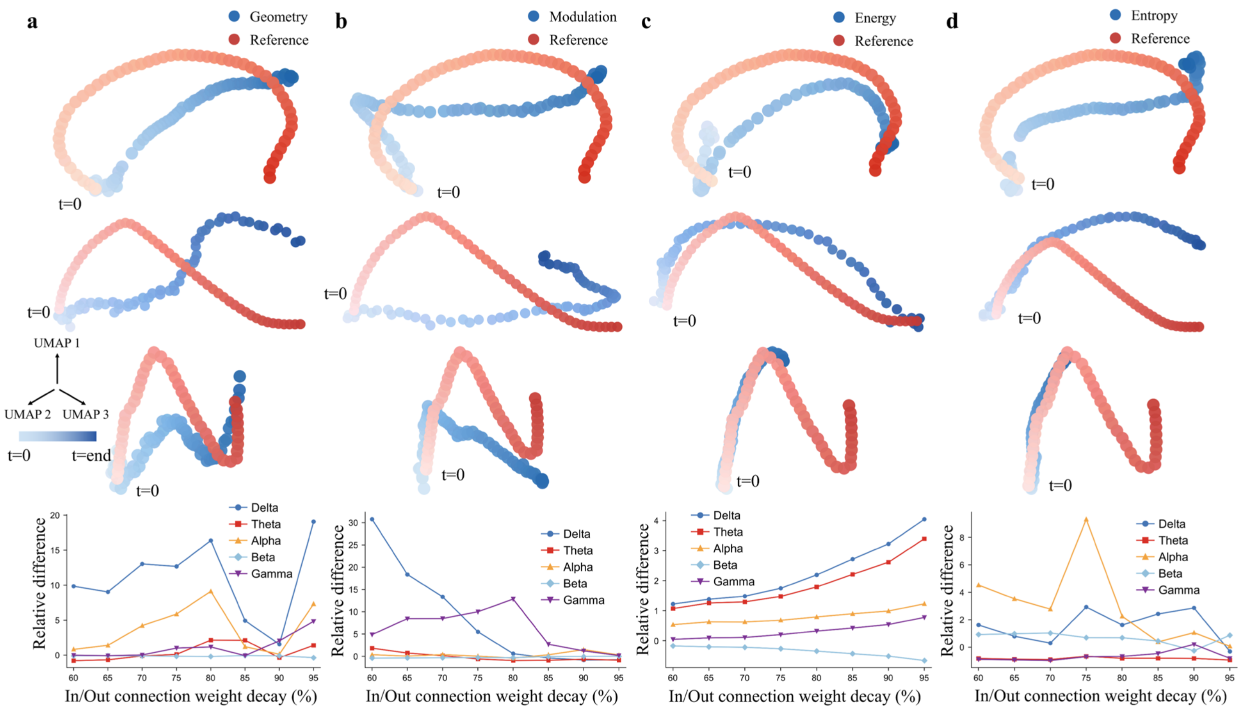

**Supplementary Fig. 6. Intermediate and high-level functional phenotypes under ablations of four structural constraints. a–d,** Ablations of the geometric power-law (**a**), node-level connection-propensity modulation (**b**), energy-budget (**c**) and information-entropy maximisation (**d**) constraints. In each column (top to bottom): low-dimensional spatiotemporal trajectories of ACx, PL and OFC compared with reference trajectories from previous studies (red, previous study; blue, ablated network; colour intensity indicates temporal progression; t=0, start); and, after graded attenuation of incoming/outgoing HPC connections, the dependence of HPC relative power across five LFP bands on attenuation strength.

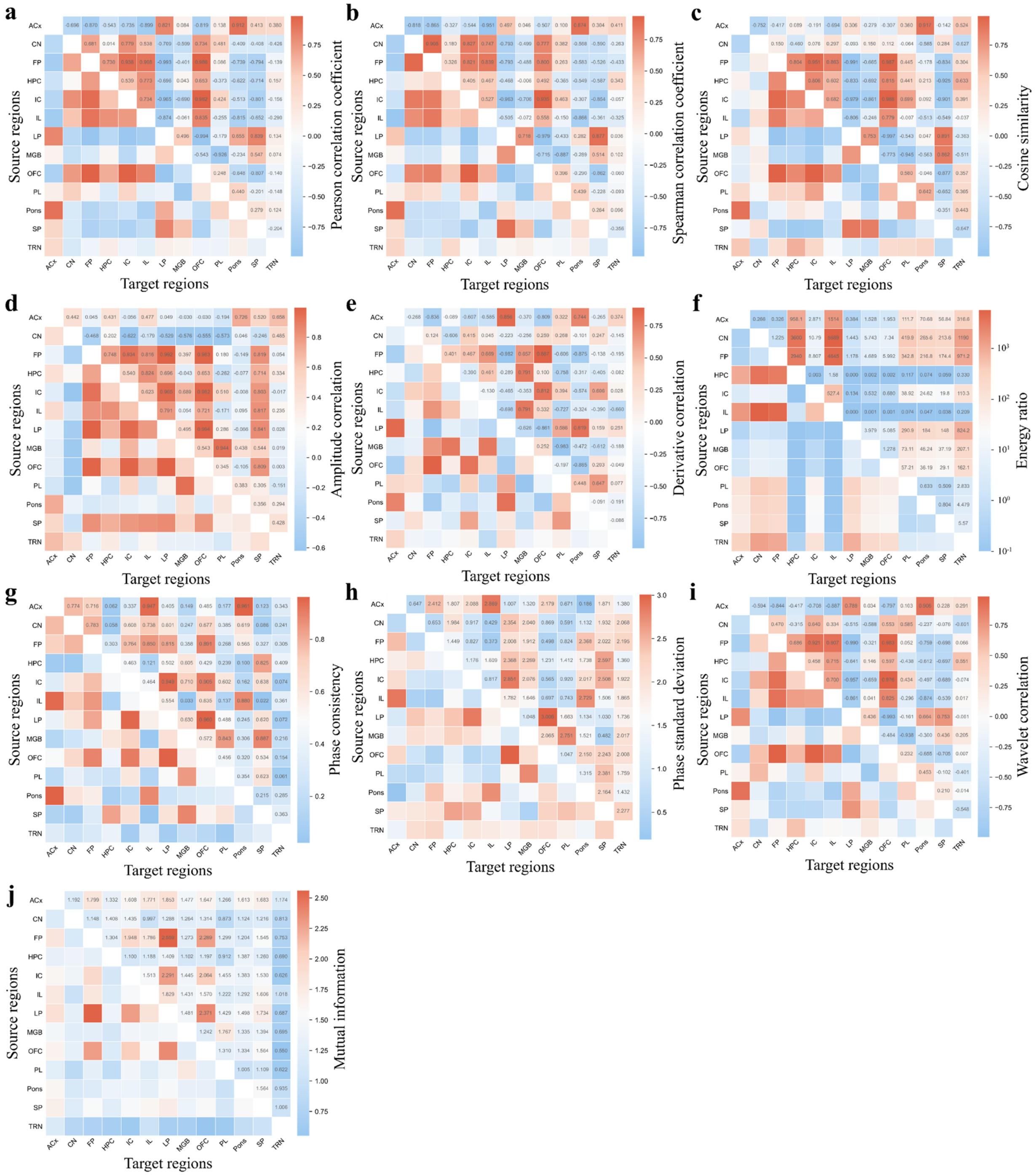

**Supplementary Fig. 7. Additional metrics for cross-regional rhythmic coupling in the spoken-digit task.** Beyond PLV reported in the main text, we summarised additional similarity and coupling measures that characterise cross-regional rhythmic coordination. Each panel is shown as a source × target matrix. **a–j** correspond to Pearson correlation, Spearman correlation, cosine similarity, amplitude correlation, derivative correlation, energy ratio, phase consistency, phase standard deviation, wavelet correlation and mutual-information, respectively.

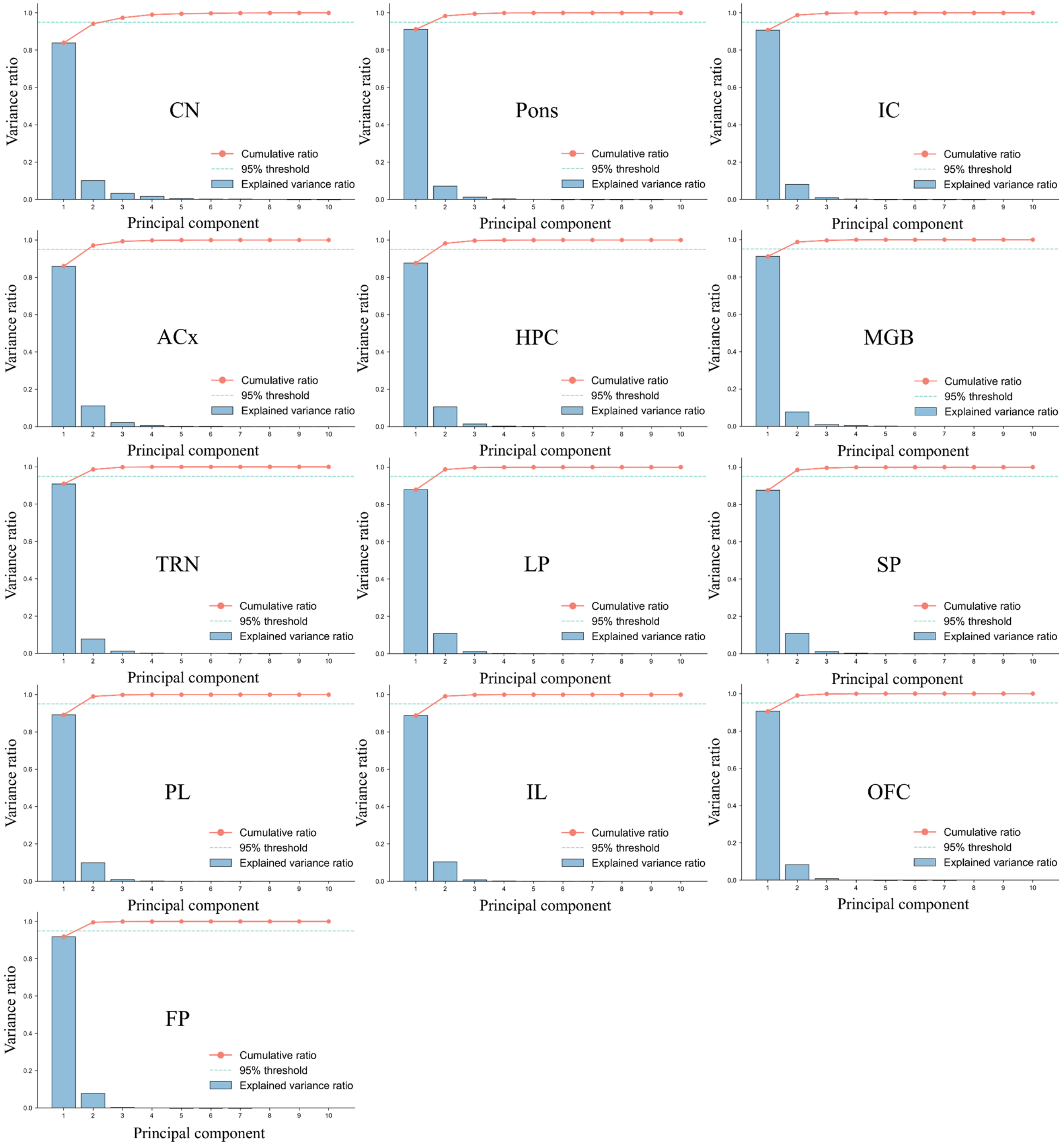

**Supplementary Fig. 8. Explained-variance spectra from PCA for each region in the auditory pathway.** PCA explained-variance analyses for each region (ordered left to right and top to bottom: CN, Pons, IC, ACx, HPC, MGB, TRN, LP, SP, PL, IL, OFC and FP). In each panel, blue bars indicate the explained-variance ratio of each principal component (PC1–PC10), the red line indicates the cumulative explained-variance ratio, and the cyan dashed line marks a reference threshold for cumulative explained variance. The x axis denotes principal component number and the y axis denotes explained-variance ratio.

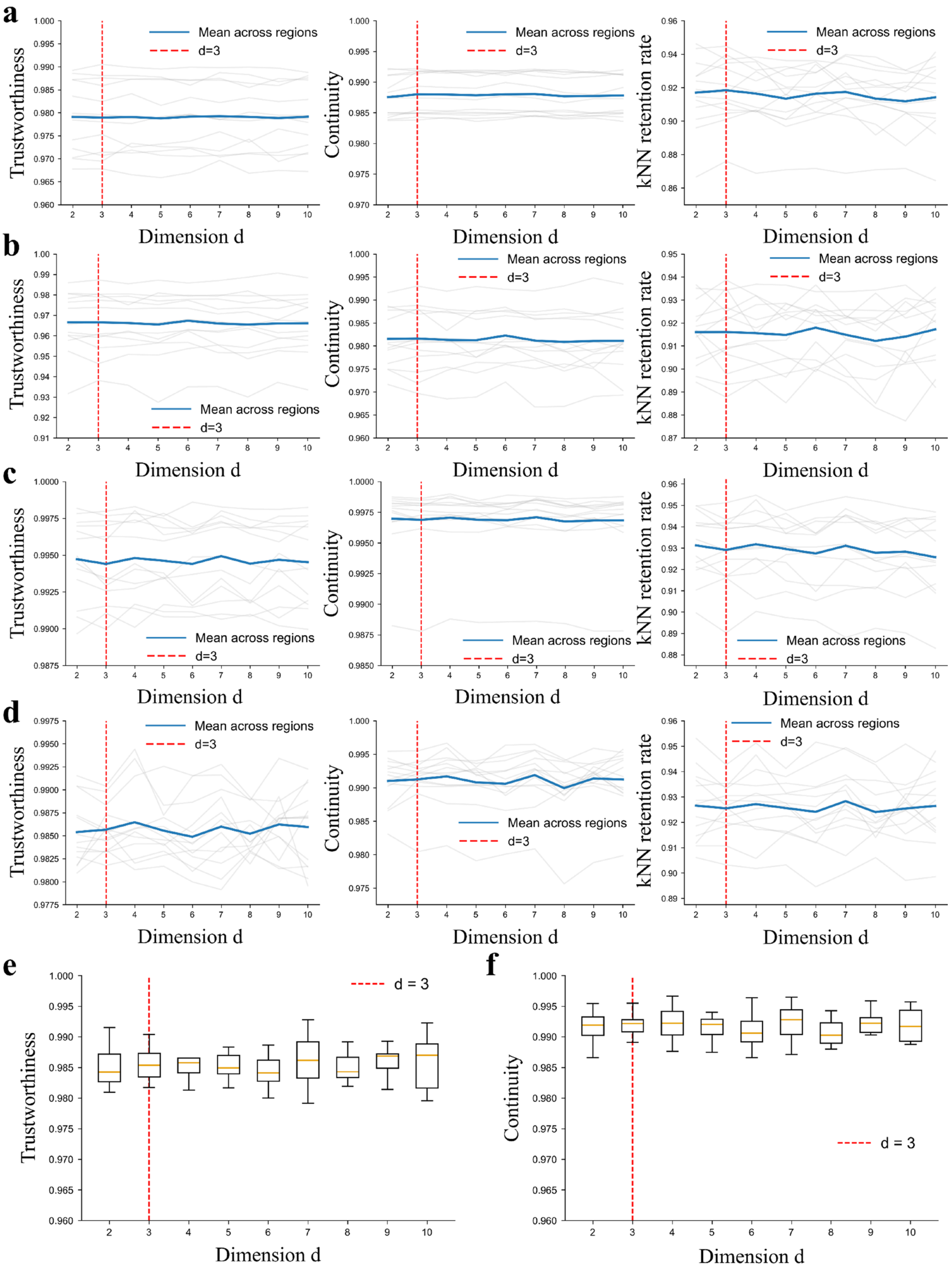
a
b
c
d
e
f
Trustworthiness
Continuity
kNN retention rate
Dimension d
Mean across regions
d=3
d = 3

**Supplementary Fig. 9. Quality metrics for selecting UMAP embedding dimensionality. a–d,** Metric values (y axis) as a function of UMAP embedding dimensionality d (x axis, 2–10) under different neighbourhood sizes k and distance metrics. In each panel, grey lines denote region-specific curves and the blue line denotes the mean across regions; the red dashed line marks the adopted dimensionality (d=3). From left to right, columns show Trustworthiness, Continuity and kNN preservation. **a,** k=15 with Euclidean distance. **b,** k=30 with Euclidean distance. **c,** k=15 with cosine distance. **d,** k=30 with cosine distance. **e-f,** Boxplots summarising Trustworthiness (**e**) and Continuity (**f**) across all regions for each dimensionality d.

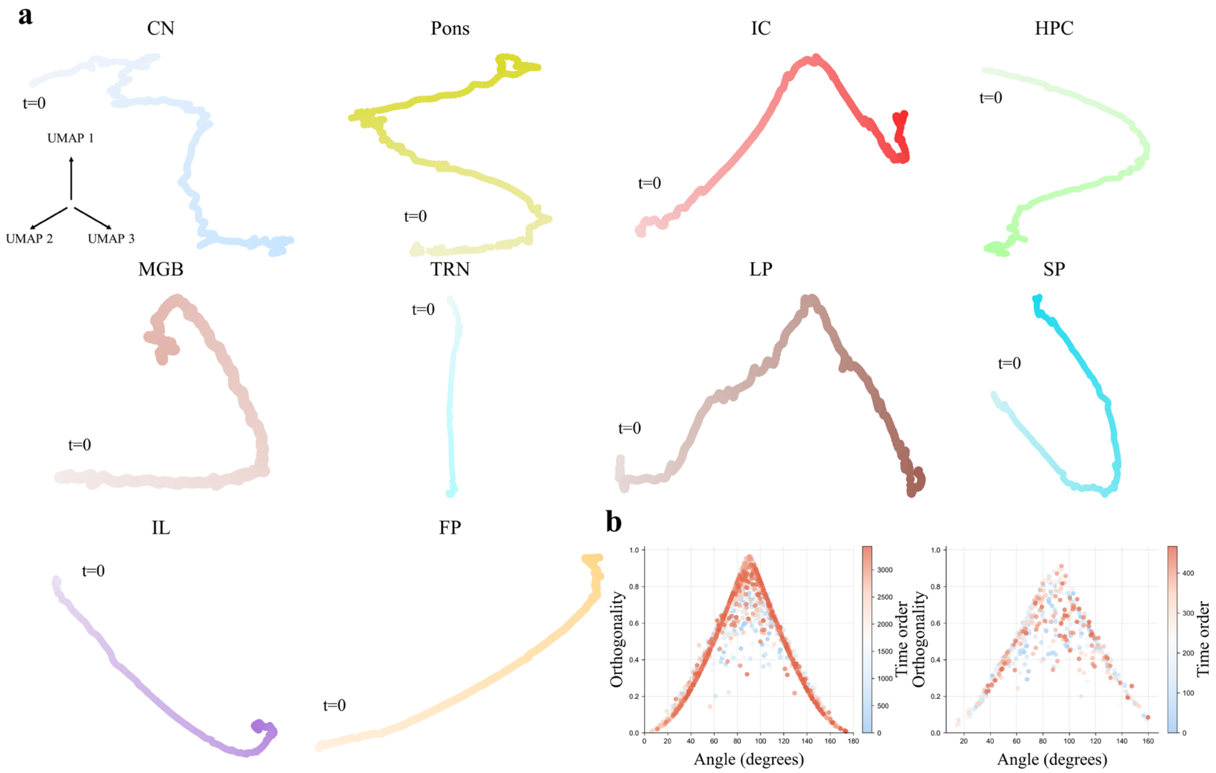

**Supplementary Fig. 10. UMAP spatiotemporal trajectories for additional auditory regions and dependence on analysis window length. a,** UMAP trajectories for the remaining 10 regions beyond ACx, PL and OFC highlighted in the main text (ordered left to right and top to bottom: CN, Pons, IC, HPC, MGB, TRN, LP, SP, IL and FP) during the auditory task. Colour intensity indicates temporal progression, with t=0 marking the start. **b,** Relationship between trajectory angle and orthogonality under different window lengths (left, 5-step window; right, 25-step window). The x axis denotes angle and the y axis denotes orthogonality; point colour encodes temporal order.

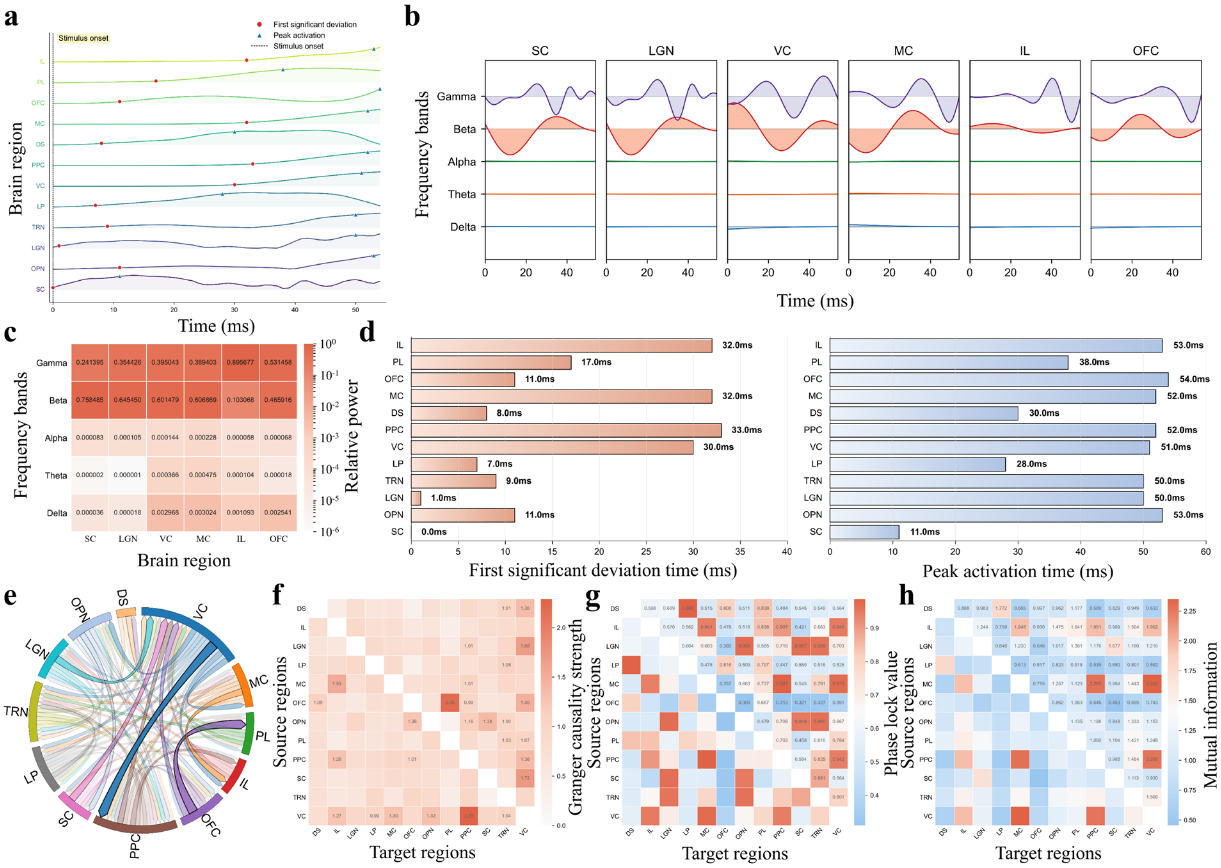

**Supplementary Fig. 11. Basic dynamical fingerprints for the visual pathway in the visual cognitive task.** Using the same analysis pipeline as in the spoken-digit task, we report basic dynamical fingerprints of the visual pathway under video-task drive. **a,** Regional LFP time series for 12 regions; red circles and blue triangles mark the time points at which the LFP first reaches 10% and 90% of its peak, respectively. **b,** Examples of band-pass-filtered LFP components in five bands (δ/θ/α/β/γ) for six representative regions (SC, LGN, VC, MC, IL and OFC). **c,** Relative power across the five bands corresponding to **b**. **d,** Hierarchical delays: time to first reach 10% of peak (left) and 90% of peak (right). **e,** Chord diagram of Granger-causality strengths showing the top 50% of cross-regional links ranked by strength; arrows indicate direction (source to target) and line width encodes causality strength. **f,** Cross-regional Granger-causality matrix (source × target). **g,** Cross-regional PLV phase-locking matrix (source × target). **h,** Cross-regional mutual-information matrix (source × target).

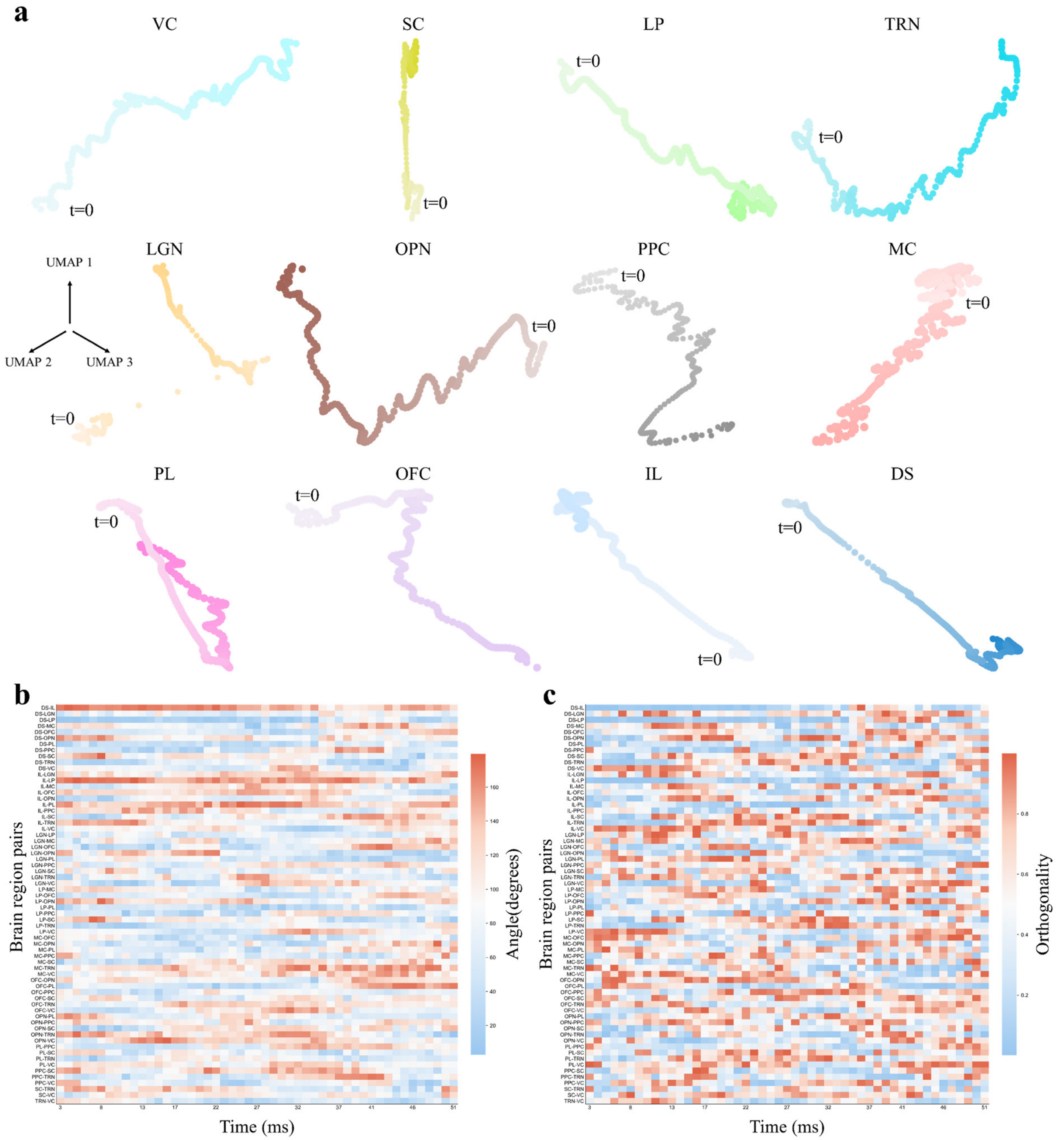

**Supplementary Fig. 12. Intermediate functional analysis for the visual cognitive task: low-dimensional trajectories and cross-regional trajectory geometry. a,** Spatiotemporal trajectories of 12 regions in the visual pathway in a low-dimensional space (ordered left to right and top to bottom: VC, SC, LP, TRN, LGN, OPN, PPC, MC, PL, OFC, IL and DS); colour intensity indicates temporal progression, with t=0 marking the start. **b,** Heatmap of the time-varying angle between trajectories for each region pair. The x axis denotes time, the y axis lists region-pair identities, and the colour bar indicates angle values. **c,** Heatmap of the time-varying orthogonality between trajectories for each region pair. The x axis denotes time, the y axis lists region-pair identities, and the colour bar indicates orthogonality.

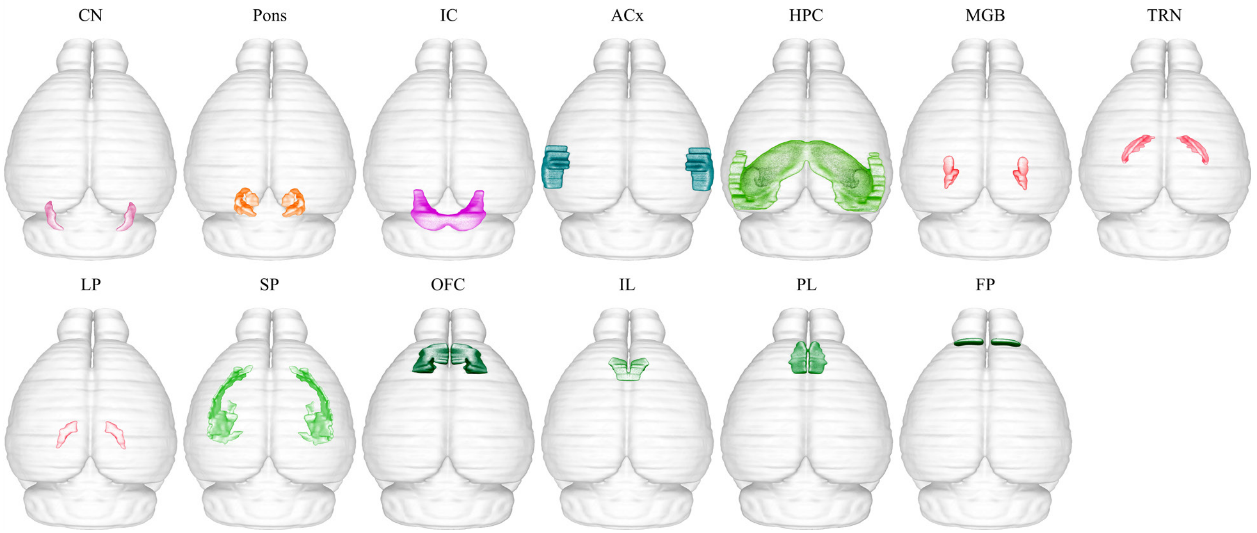

**Supplementary Fig. 13. Anatomical locations of the 13 regions used in the auditory cognitive task[46].** Panels are ordered left to right and top to bottom as: CN, Pons, IC, ACx, HPC, MGB, TRN, LP, SP, OFC, IL, PL and FP. Region abbreviations: CN, cochlear nuclei; IC, inferior colliculus; ACx, auditory cortex; HPC, hippocampus; MGB, medial geniculate body; TRN, thalamic reticular nucleus; LP, lateral posterior nucleus; SP, cortical subplate; OFC, orbitofrontal cortex; IL, infralimbic cortex; PL, prelimbic cortex; FP, frontal pole.

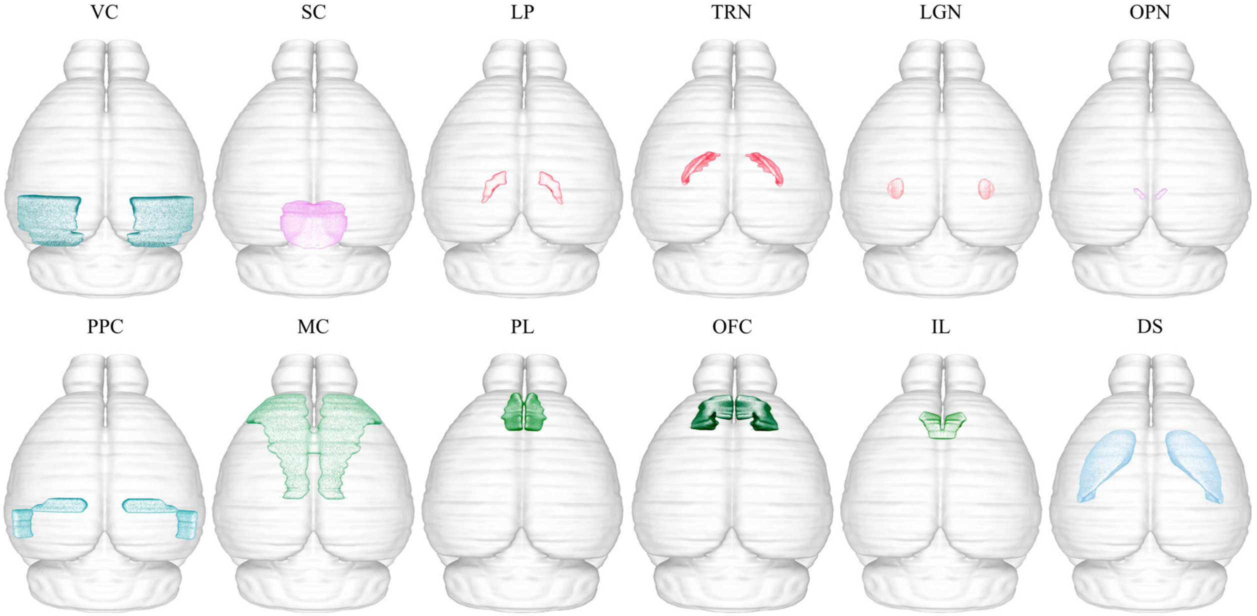

**Supplementary Fig. 14. Anatomical locations of the 12 regions used in the visual cognitive task[46].** Panels are ordered left to right and top to bottom as: VC, SC, LP, TRN, LGN, OPN, PPC, MC, PL, OFC, IL and DS. Region abbreviations: VC, visual cortex; SC, superior colliculus; LP, lateral posterior nucleus; TRN, thalamic reticular nucleus; LGN, lateral geniculate nucleus; OPN, olivary pretectal nucleus; PPC, posterior parietal cortex; MC, motor cortex; PL, prelimbic cortex; OFC, orbitofrontal cortex; IL, infralimbic cortex; DS, dorsal striatum.

### S8. Supplementary Tables

**Supplementary Table 1. Model fits and information-criterion comparisons for four candidate distance-decay models of the neuronal-scale distance–connection-probability relationship.** This table summarises evaluation metrics obtained by fitting four commonly used distance-decay models (exponential, power-law, Gaussian and logistic) to neuron-pair connectivity data in mouse V1. All models were fitted over the same distance range; parameters were estimated by maximum likelihood using the raw neuron-pair data. We report goodness-of-fit $R^2$, log-likelihood, and information criteria (AIC and BIC). Larger $R^2$ and log-likelihood indicate better fits, whereas smaller AIC and BIC indicate better model support. Bold values denote the best-performing model for each metric.

| Metric | Exponential | Power law | Gaussian kernel | Logistic |
|---|---|---|---|---|
| $R^2$ | 0.84 | **0.94** | 0.61 | 0.82 |
| AIC | 145.87 | **105.21** | 180.23 | 153.00 |
| BIC | 149.20 | **108.53** | 183.56 | 157.99 |
| Log-likelihood | -70.94 | **-50.60** | -88.11 | -73.50 |

**Supplementary Table 2. Structural similarity of degree distributions: NIGC-generated connectomes versus three baseline models.** For each model, we generated a connectome, computed its degree distribution, and compared it with the empirical degree distribution using three metrics: cosine similarity (higher indicates closer agreement), Jensen–Shannon (JS) divergence and Bray–Curtis dissimilarity (lower indicates closer agreement). Bold values denote the best model for each metric.

| Metric | NIGC | Economical wiring | Homophily | Fully random |
|---|---|---|---|---|
| Cosine similarity | **0.997** | 0.958 | 0.954 | 0.952 |
| JS divergence | **0.133** | 0.242 | 0.251 | 0.260 |
| Bray–Curtis dissimilarity | **0.059** | 0.209 | 0.218 | 0.226 |

**Supplementary Table 3. Structural similarity of clustering-coefficient distributions: NIGC-generated connectomes versus three baseline models.** For each model, we computed the node-wise clustering-coefficient distribution and compared it with the empirical distribution using three metrics: the Kolmogorov–Smirnov (KS) statistic, Jensen–Shannon (JS) divergence and Bray–Curtis dissimilarity (lower values indicate closer agreement). Bold values denote the best model for each metric.

| Metric | NIGC | Economical wiring | Homophily | Fully random |
|---|---|---|---|---|
| KS statistic | **0.480** | 0.906 | 0.900 | 0.685 |
| JS divergence | **0.413** | 0.800 | 0.796 | 0.609 |
| Bray–Curtis dissimilarity | **0.526** | 0.939 | 0.935 | 0.945 |

**Supplementary Table 4. Effects of constraint ablations on degree-distribution similarity.** We quantified how ablation of individual NIGC constraints affects agreement with the empirical degree distribution. Connectomes were generated under three ablation conditions (removing node-level connection-propensity modulation, the energy-budget constraint, or the information-entropy maximisation constraint) and compared with the empirical distribution using cosine similarity, Jensen–Shannon (JS) divergence and Bray–Curtis dissimilarity. Results for the full NIGC model are shown for reference. In the column headers, w/o denotes without.

| Metric | NIGC | w/o Node propensity | w/o Energy | w/o Max-entropy |
|---|---|---|---|---|
| Cosine similarity | **0.997** | 0.326 | **0.999** | 0.002 |
| JS divergence | **0.133** | 0.559 | **0.048** | 0.814 |
| Bray–Curtis dissimilarity | **0.059** | 0.729 | **0.013** | 0.982 |

**Supplementary Table 5. Effects of constraint ablations on clustering-coefficient-distribution similarity.** We evaluated the role of key NIGC constraints in shaping local clustering organisation. Connectomes were generated under three ablation conditions (removing node-level connection-propensity modulation, the energy-budget constraint, or the information-entropy maximisation constraint), and node-wise clustering-coefficient distributions were computed. Each distribution was compared with the empirical distribution using the Kolmogorov–Smirnov (KS) statistic, Jensen–Shannon (JS) divergence and Bray–Curtis dissimilarity (lower values indicate closer agreement). Results for the full NIGC model are shown for reference. In the column headers, w/o denotes without.

| Metric | NIGC | w/o Node propensity | w/o Energy | w/o Max-entropy |
|---|---|---|---|---|
| KS statistic | **0.480** | 0.573 | 0.612 | **0.259** |
| JS divergence | **0.413** | 0.533 | 0.558 | 0.426 |
| Bray–Curtis dissimilarity | **0.526** | 0.571 | 0.884 | 0.608 |

**Supplementary Table 6. Spoken Arabic Digit classification: performance and efficiency of NIGC–ESN versus end-to-end networks with matched trainable parameter counts.** The NIGC-generated connectome was embedded as a fixed recurrent reservoir in an ESN (NIGC–ESN) and compared with four end-to-end networks (LSTM, CNN, Transformer and GNN). Each method was repeated 20 times under identical experimental settings. Test accuracy and runtime (s) are reported as mean ± s.d. The composite score is defined as test accuracy divided by runtime[15] (higher values indicate higher accuracy per unit time). Bold values denote the best method for each metric.

| Metric | NIGC–ESN | LSTM | CNN | Transformer | GNN |
|---|---|---|---|---|---|
| Test accuracy | **0.898±0.009** | 0.419±0.119 | 0.818±0.026 | 0.749±0.043 | 0.892±0.008 |
| Runtime (s) | **18.6±0.56** | 21.8±0.03 | 61.0±1.36 | 44.2±0.26 | 69.6±1.16 |
| Composite score | **0.048** | 0.019 | 0.013 | 0.017 | 0.013 |

**Supplementary Table 7. Spoken Arabic Digit classification: performance and efficiency of NIGC–ESN versus end-to-end networks with matched total parameter counts.** For NIGC–ESN, the total parameter count equals the number of non-trainable reservoir parameters (specified by the reservoir connectivity) plus the number of trainable readout parameters; this total was matched to the parameter counts of end-to-end networks (LSTM, CNN, Transformer and GNN) to enable comparisons under a unified definition of model capacity. Each method was repeated 20 times under identical experimental settings. Bold values denote the best method for each metric.

| Metric | NIGC–ESN | LSTM | CNN | Transformer | GNN |
|---|---|---|---|---|---|
| Test accuracy | 0.898±0.009 | 0.783±0.172 | 0.985±0.002 | **0.987±0.006** | 0.945±0.002 |
| Runtime (s) | **18.7±0.43** | 1006.1±178.47 | 63.5±1.56 | 158.9±1.16 | 96.2±2.31 |
| Composite score | **0.048** | 0.001 | 0.016 | 0.006 | 0.010 |

**Supplementary Table 8. Comparative summary of representative approaches corresponding to the three routes discussed in the Discussion.** Route indicates the conceptual route (this work; Route 1–3). Journal lists the venue of the representative study. Framework provides a brief methodological descriptor. Constrain indicates whether the approach uses a concise and separable set of wiring constraints. Structural denotes the extent to which key neuronal-scale structural statistics of connectivity are reproduced. Functional denotes the extent to which biologically consistent functional phenotypes are reproduced under controlled dynamics. Reproducibility denotes the relative practical ease of reproducing the core results (considering accessibility of data and typical computational requirements). √ indicates that the criterion in Constrain is explicitly satisfied; × indicates that it is not satisfied or not directly established in the study's core formulation. Star ratings are reported on a five-level ordinal scale (more stars indicate higher structural concordance, higher functional concordance, or greater reproducibility, respectively) and are intended for relative comparison across entries. For this work, the listed route corresponds to a direct test of the microscopic structure–function "concise-constraint sufficiency" hypothesis: under a fixed geometric framework and controlled dynamics, a small, explicit and ablatable set of biophysical wiring constraints is sufficient, empirically, to reproduce key neuronal-scale structural statistics and to yield multiple biologically consistent functional phenotypes without fitting functional matrices or time courses.

| Route | Ref. | Framework | Constrain | Structural | Functional | Reproducibility |
|---|---|---|---|---|---|---|
| this work | - | Explicit, concise, ablatable wiring constraints; unified evaluation under geometric embedding and controlled dynamics | √ | ★★★★★ | ★★★★★ | ★★★★ |
| Route 1 | [96] **Nature 2025** | Large-scale, data-driven foundation model; end-to-end training for neural response prediction and generalisation | × | ★★★ | ★★★★★ | ★★ |
| Route 1 | [97] **Nature Machine Intelligence 2023** | Spatially embedded RNN; sparse connectivity and cost regularisation shaping structure and representations | × | ★★★ | ★★★ | ★★★ |
| Route 1 | [98] | Genomic-bottleneck weight compression and generation; priors encoded for task performance | × | ★★ | ★★★ | ★★★★ |

| | | | | | | |
|---|---|---|---|---|---|---|
| | **PNAS 2024** | | | | | |
| Route 2 | [94] **Nature Communications 2025** | Synthetic axonal morphologies for brain-wide connectivity; constrained primarily by morphology and projection statistics | × | ★★★★★ | ★★ | ★★★ |
| Route 3 | [95] **Nature Computational Science 2024** | Measurement-scaffolded simulation and assimilation; large-scale spiking modelling to approximate macroscopic signals | × | ★★★ | ★★★★ | ★ |
